# Mapping Cerebellar Anatomical Heterogeneity in Mental and Neurological Illnesses

**DOI:** 10.1101/2023.11.18.567647

**Authors:** Milin Kim, Esten Leonardsen, Saige Rutherford, Geir Selbæk, Karin Persson, Nils Eiel Steen, Olav B. Smeland, Torill Ueland, Geneviève Richard, Christian F. Beckmann, Andre F. Marquand, Alzheimer’s Disease Neuroimaging Initiative (ADNI), Ole A. Andreassen, Lars T. Westlye, Thomas Wolfers, Torgeir Moberget

**Affiliations:** Norwegian Centre for Mental Disorders Research (NORMENT), Division of Mental Health and Addiction, Oslo University Hospital & Institute of Clinical Medicine, University of Oslo, Oslo, Norway; Department of Psychology, Faculty of Social Sciences, University of Oslo, Norway; KG Jebsen Centre for Neurodevelopmental Disorders, University of Oslo, Oslo, Norway; Department of Behavioral Science, School of Health Sciences, Oslo Metropolitan University - OsloMet, Oslo, Norway; Department of Cognitive Neuroscience, Radboud University Medical Centre, Nijmegen, Netherlands; Donders Institute, Radboud University, Nijmegen, Netherlands; Department of Psychiatry, University of Michigan, Ann Arbor, MI, United States; Department of Psychiatry and Psychotherapy, Tübingen Center for Mental Health, University of Tübingen, Germany; German Center for Mental Health (DZPG); Department of Geriatric Medicine, Oslo University Hospital, Oslo, Norway; The Norwegian National Centre for Ageing and Health, Vestfold Hospital Trust, Tønsberg, Norway; Department of Psychiatric Research, Diakonhjemmet Hospital, Oslo, Norway; Department of Neuroimaging, Centre of Neuroimaging Sciences, Institute of Psychiatry, King’s College London, London, UK; Centre for Functional MRI of the Brain, Nuffield Department of Clinical Neurosciences, Wellcome Centre for Integrative Neuroimaging, University of Oxford, Oxford, UK

**Keywords:** Cerebellum, Normative modelling, Magnetic Resonance Imaging, Individual-level Inference, Heterogeneity Mapping, Schizophrenia, Alzheimer, Autism Spectrum Disorder, Mental Disorders, & Neurological Diseases

## Abstract

The cerebellum has been linked to motor coordination, cognitive and affective processing, in addition to a wide range of clinical illnesses. To enable robust quantification of individual cerebellar anatomy relative to population norms, we mapped the normative development and aging of the cerebellum across the lifespan using brain scans of > 54k participants. We estimated normative models at voxel-wise spatial precision, enabling integration with cerebellar atlases. Applying the normative models in independent samples revealed substantial heterogeneity within five clinical illnesses: autism spectrum disorder, mild cognitive impairment, Alzheimer’s disease, bipolar disorder, and schizophrenia. Notably, individuals with autism spectrum disorder and mild cognitive impairment exhibited increased numbers of both positive and negative extreme deviations in cerebellar anatomy, while schizophrenia and Alzheimer’s disease predominantly showed negative deviations. Finally, extreme deviations were associated with cognitive scores. Our results provide a voxel-wise mapping of cerebellar anatomy across the human lifespan and clinical illnesses, demonstrating cerebellum’s nuanced role in shaping human neurodiversity across the lifespan and in different clinical illnesses.

## Introduction

The cerebellum accounts for 10 to 15% of the brain’s volume^1^ and contains approximately 80% of all neurons^2^. The cerebellum has long been recognized for its involvement in motor functions^3^, but has also been linked to human cognitive capacities such as language and social intelligence^4–7^. In line with this expanded view of cerebellar function, the cerebellum has also been implicated in a wide range of mental and neurological illnesses characterized by both motor and cognitive deficits^8^. However, empirical findings on cerebellar structure and function in different mental and neurological illnesses are varied^9,10^. A growing literature^11,12^ suggests a key role for cerebellum in autism spectrum disorder (ASD). However, a meta-analysis^13^ of human MRI studies revealed no significant group differences in cerebellar volume between individuals with ASD and individuals without diagnosis, suggesting the importance of replication and the inherent uncertainty that comes with small studies^14^. Such inconsistent findings may also be due to the high degree of heterogeneity within clinical groups, a feature which can be explicitly investigated through the construction of normative models.

Normative modelling draws its inspiration from paediatric growth charts, which chart key aspects of a child’s development, such as weight or height, over the early years of life. As a machine learning framework, normative modelling is designed to estimate a prototypical and representative developmental trajectory, by mapping different types of variables onto each other, e.g. such as mapping age and sex onto different brain measures^15^. Once such models are established, individuals can be placed in reference to the resulting norms, from which individual-level normative probability maps can be derived. These represent the extent to which an individual deviates from the estimated norm in locations across the brain. In this respect the use of the normative models enables comparison across studies, scanner sites and lifespan stages by binding to a common reference framework and thus makes studies more comparable^16^. In earlier applications of this approach, we charted the normative trajectory of cortical thickness and subcortical volume^17^ across the lifespan. When applied to clinical samples, these normative models revealed a high degree of heterogeneity for individual-level profiles within same condition. This pattern has now been demonstrated and replicated^18^ across many different clinical illnesses, such as ASD^19–21^, attention deficit hyperactivity disorder (ADHD)^22^, dementia including AD (Alzheimer’s Disease)^23,24^, first-episode psychosis^25^, bipolar disorder (BD) and schizophrenia (SZ)^18,26^. Of note, a recent paper demonstrated that while phenotypic heterogeneity within same order may be due to heterogeneity in regional deviations, phenotypic similarities can be coupled to common functional circuits and networks^27^. Together, these findings highlight the need for mapping heterogeneity of complex clinical illnesses using voxel-wise, regional, and network-based approaches.

In the current study, we chart the normative development and aging of the human cerebellum at lobular and voxel-wise spatial precision in a sample of > 54k individuals. We map the heterogeneity of five distinct clinical phenotypes accounting for 132 scanning sites (Table 1 and Supplementary Table 1) and > 143k voxels, providing a generalisable and state-of-the-art reference model for cerebellar measures across the lifespan. Further, and based on cytoarchitectonic, functional and regional parcellations of the cerebellum, we link percentage of extreme deviations to cognitive measures and symptom scores across the groups and in this way shed light on the role of individual-level deviations from normative cerebellar anatomy across five mental and neurological illnesses.

**Table 1.** Sample description and demographics.

|  |  | N (Participants) | N (Scanners) | Age (Mean, S.D.) | Sex (%F:%M) |
| --- | --- | --- | --- | --- | --- |
| Full | All | 54102 | 132 |  |  |
|  | Training set | 27117 | 132 | 54.36 (20.31) | 0.53:0:47 |
|  | Testing set | 26985 | 132 | 54.52 (20.19) | 0.53:0:47 |
| Clinical | Testing set | 1757 | 53 | 29.40 (20.84) | 0.30 0:70 |
|  | Alzheimer's Disease | 146 | 13 | 72.42 (7.65) | 0.53:0:47 |
|  | ASD | 900 | 37 | 16.20 (9.00) | 0.14:0:86 |
|  | Bipolar Disorder | 277 | 3 | 32.73 (11.67) | 0.60:0:40 |
|  | Mild Cognitive<br>Impairment | 122 | 3 | 67.25 (9.27) | 0.42:0:58 |
|  | Schizophrenia | 312 | 3 | 29.58 (9.52) | 0.33:0:67 |

## Results

### Lifespan trajectories of cerebellar morphology

Normative models were trained on MRI-derived cerebellar features from the training portion of the overall dataset (Table 1), comprising more than 27k individuals without diagnosis across the lifespan (Fig. 1A). All normative models included age and sex as covariates, while controlling for scanning sites related variability. Cerebellar morphology was quantified as lobular volumes, or voxel-wise grey matter probability values (multiplied by the Jacobian of the transformation matrix to preserve volumetric information after normalization) (see Online Methods and Supplementary Fig. 2 and 3 for details). Figure 1B shows selected normative trajectories of cerebellar lobules. Most of the lobules exhibit a pattern of volume increase until around age 19, followed by a gradual decrease. Notably, we observed flatter trajectories in specific lobules, namely Left X, Left VIIIB, Corpus Medullare, Right VIIIB, Vermis VI, and Vermis VIII. To validate our model fit, we report the performance of these models, in terms of explained variance, plotted on the cerebellar flat maps (Figure 1B and 1C). The lobule with the best model fit was in Right V, while the worst fit was found in Vermis X. Additional evaluation metrics such as kurtosis, skew, and mean squared logarithmic loss (MSLL) are shown in Supplementary Figure 1. Additionally, Figure 1C presents voxel-wise trajectories, as chosen from the 143k models estimated for individual cerebellar grey matter voxels.

**Figure 1.**
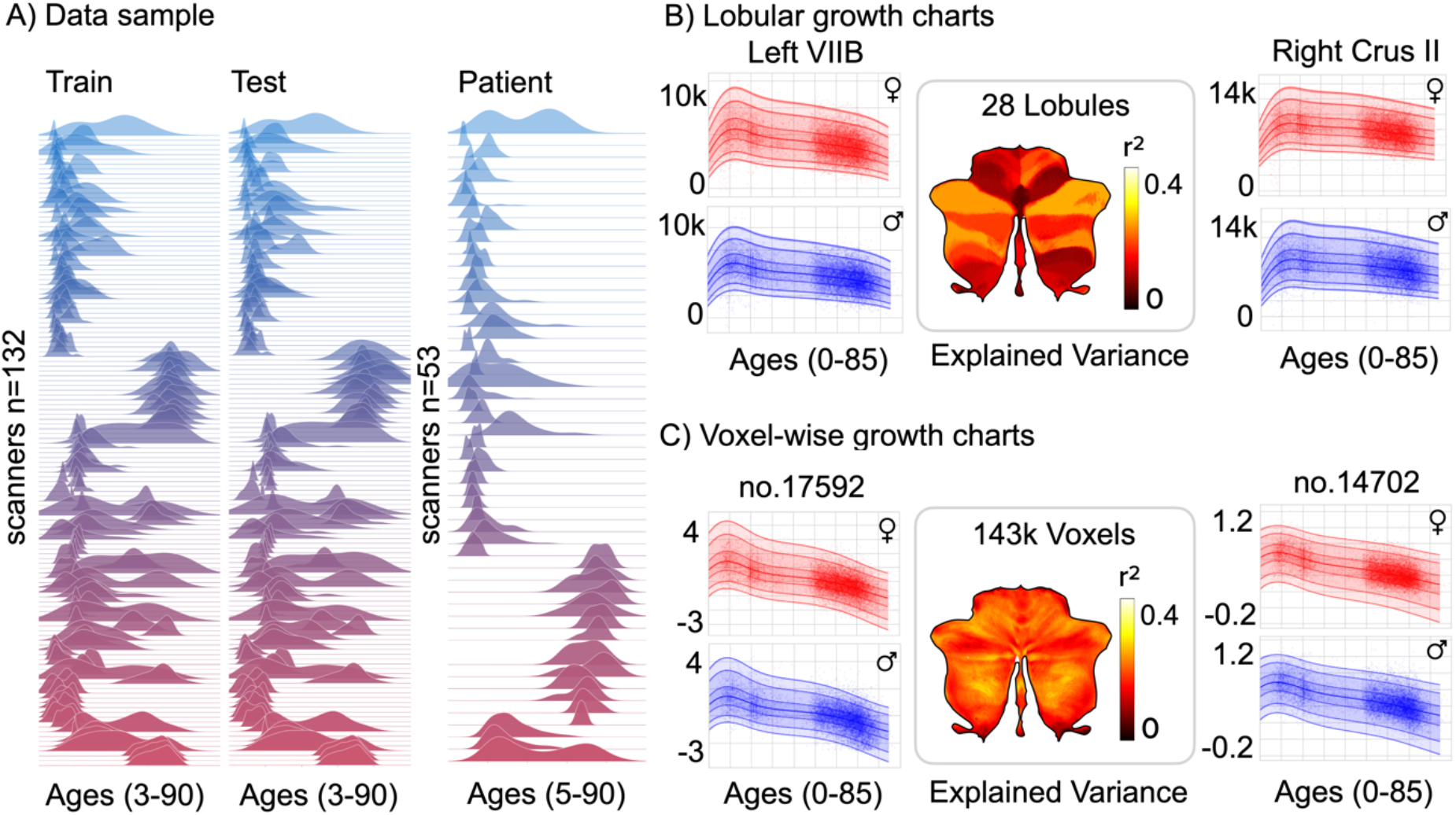
Normative models based on MRI data from > 54k participants describe the lifespan trajectories of cerebellar lobules and individual voxels. In panel (A), the age density distribution is displayed for each scanning sites in the training, test, and clinical sets. Panel (B-C) showcase two of the 28 regions representing the lobular growth charts and two of the 143k voxel-wise growth charts for each sex. The x-axis represents age, ranging from 3 to 85, while the y-axis represents the predicted cerebellar volume and grey matter probability values. Additionally, the figure includes the explained variance, indicating the goodness of fit.

### Altered cerebellar morphology across clinical groups

Figure 2A display nonparametric comparison, Mann-Whitney U-test, of voxel-wise model-derived z-scores between patient groups and individuals without diagnosis in the test datasets (for lobule-wise results, see Supplementary Figure 4A and Supplementary Table 3). All reported effects in the result section are corrected for multiple comparisons using Bonferroni correction (across regions of interest or voxels, number of clinical groups and directions) and only significant results are shown in the figures (corrected p < 0.05). Of note, we replicated previous reports of reduced cerebellar volumes in SZ by observing significant negative effect sizes (rank biserial correlations) of deviations in the negative direction, indicating greater strength of SZ than individuals without diagnosis. Similar patterns were seen for SZ in lobular volumes, while AD and MCI exhibited a notably different pattern of negative deviations in lobular volumes compared to voxel-wise maps and furthermore in uncorrected maps (Supplementary Fig. 5). Significant negative effects were observed in Right V for MCI and AD in addition to Right Crus I for AD when corrected. ASD showed negative deviations in the anterior regions for lobular volumes while showed small negative effects in voxel-wise maps in the posterior regions. BD did not show noticeable effects in terms of z-scores with the individuals without diagnosis after multiple comparisons.

**Figure 2.**
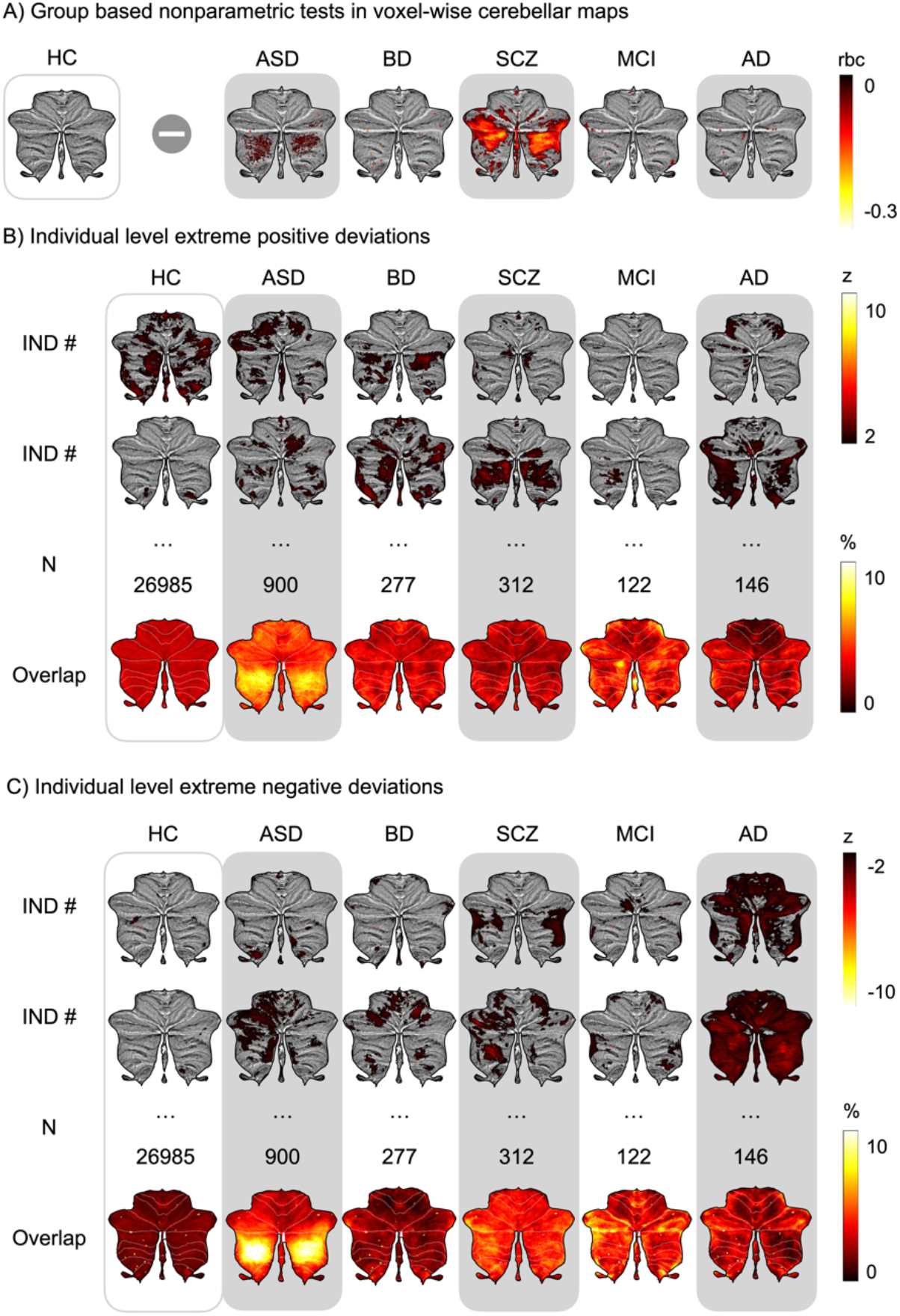
The voxel-wise deviations from estimated norms show high levels of heterogeneity within diagnostic groups. (A) depict Mann-Whitney U-test rank biserial correlation (rbc) of z-scores between the clinical and the individuals without diagnosis in voxel-wise. The effects are corrected for multiple comparisons using Bonferroni correction (corrected p < 0.05). Specifically, patients diagnosed with schizophrenia (SZ) and autism spectrum disorder (ASD) exhibited significant effects compared to individuals without diagnosis (HC) in the voxel-wise maps. (B-C) The z-scores of extreme positive and negative deviations (|z| > 1.96) are shown for two individuals per cohort. Overlap maps calculate percentage of extreme deviations occurred in the same group. The clinical groups displayed a significantly higher occurrence of percentage of extreme deviations, even in cases where the group based nonparametric tests did not differ significantly. These results indicate that within the clinical groups, there were individuals who exhibited significant deviations from the normative patterns, regardless of the overall group effects.

### Individual-level extreme deviations are patient specific and transdiagnostic

The normative modelling approach allows us to quantify extreme positive and negative deviations, here defined as |z| > 1.96, at the voxel-wise and lobular levels for each individual. Figures 2B-C show examples of normative probability maps for different individuals across clinical diagnostic groups. These maps reveal high cerebellar heterogeneity within and across groups, especially when we look at the overlaps of these individual-level probability maps. The overlap maps calculate the percentage of extreme deviations within the same group for each voxel. Even within the same diagnostic group only a marginal percentage of individuals show extreme deviations in the same cerebellar regions. Moreover, some cerebellar regions (e.g., in posterior lobules) show increased numbers of extreme deviations across several diagnostic groups (e.g., SZ, ASD and AD). Analyses of lobular volumes yielded similar results (see Supplementary Fig. 4B-C).

### Norms summed across existing cerebellar atlases aid functional interpretation

Voxel-wise normative models can be projected onto any existing - or future - cerebellar atlas morphed into Montreal Neurological Institute (MNI) space (Fig. 3A). Here, the voxel-wise normative probability maps are summed across 28 lobules (anatomical atlas), 10 functionally defined cerebellar regions (task-based atlas)^28^, and 17 regions defined by their resting-state connectivity with functional networks of the cerebral cortex (resting-state atlas)^29,30^. Percentage of deviations for each region is calculated across parcellations for each individual. The figures (Fig. 3B-C and Supplementary Table 4-6) show effect sizes (rank biserial correlation) of percentage of both positive and negative extreme deviations for various clinical cohorts compared to the individuals without diagnosis. As the percentage of negative deviations increases, the strength of the clinical groups also increases. SZ, mild cognitive impairment (MCI) and AD groups showed small to medium effects in the negative deviations compared to the individuals without diagnosis globally across all three atlases. For all three groups, the effects are visibly greater in the posterior regions of the cerebellum and display consistent patterns across the atlases. However, ASD and MCI groups reveal significant small effects in positive deviations across numerous regions. For ASD, the medium effects in the negative deviations are seen in the region related to action observation and divided attention (region 4) in the task-based atlas and linked to the dorsal attention network region (network 6) in the resting state atlas.

**Figure 3.**
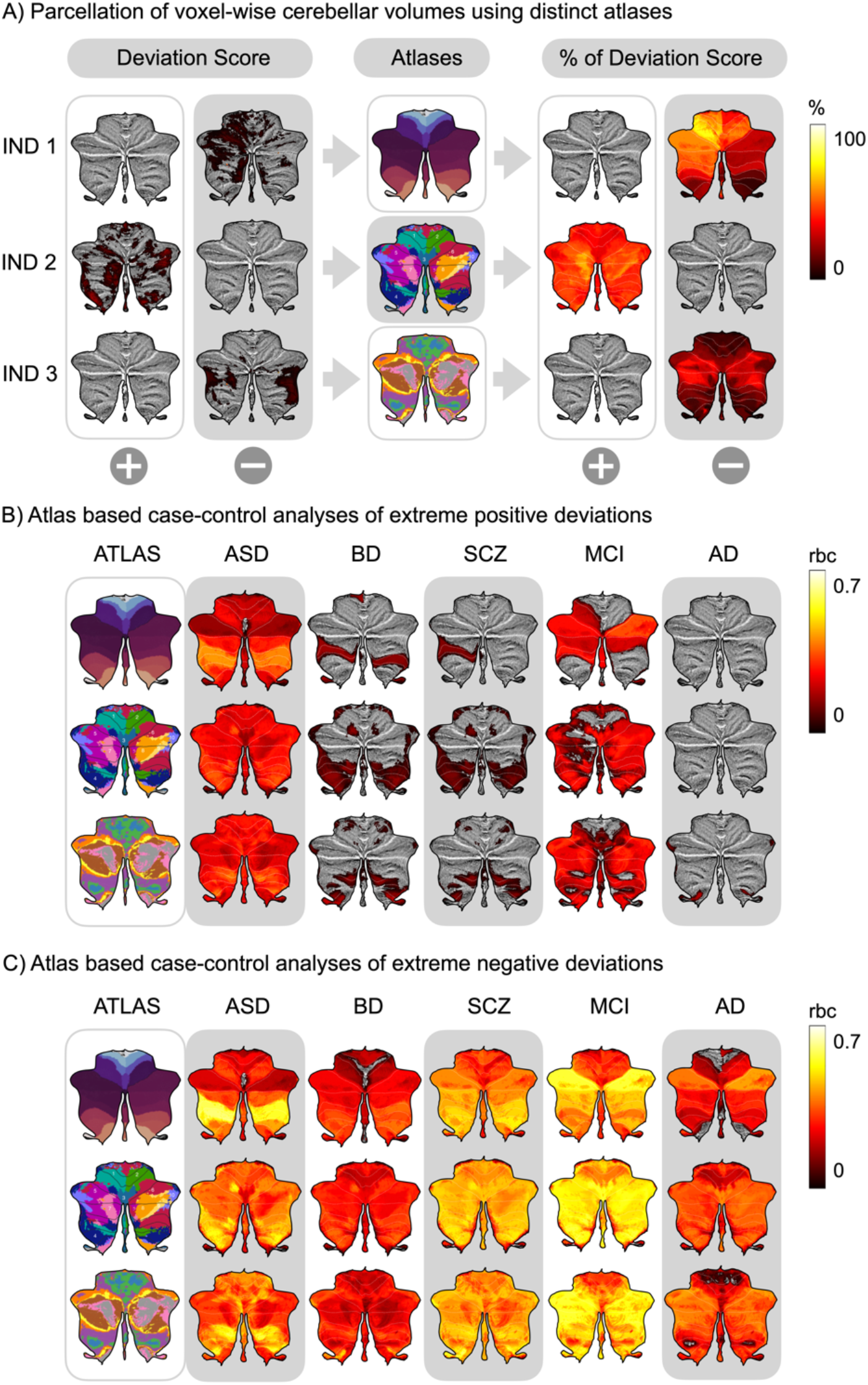
Voxel-wise normative models can be applied to existing or future cerebellar atlases. (A) The outputs from normative model, 143k features normative probability maps of an individual, are applied onto existing atlases of traditional anatomical regions, task-based regions, and resting state connectivity atlases. Panel (B-C) depict effect size of comparison of percentage of extreme positive and negative deviations of clinical cohorts to the individuals without diagnosis in voxel-wise per participant based on the three atlases. Scale indicates Mann-Whitney U-test rank biserial correlation (rbc) and only shows significant regions after multiple comparison corrections.

### Extreme deviations are associated with intelligence

Figure 4 shows correlations obtained from three distinct atlases when examining the relationship between intelligence scores. In ASD, regions associated with Right IV-VI in anatomical atlas and somatomotor B (network 4) in resting state atlas showed positive correlations between percentage of extreme positive deviations per participant and performance IQ (PIQ) (Supplementary Table 7-9) that survived multiple comparisons of number of tests, regions of interest and directions. Weak positive correlations between the percentage of extreme positive deviations per participant and intelligence scores were shown in patients with SZ across three atlases and vice versa for negative deviations (Figure 4B and Supplementary Table 13-15). However, symptom scores Autism Diagnostic Observation Schedule (ADOS) for ASD and Positive and Negative Syndrome Scale (PANSS) for SZ did not show significant associations with percentage of extreme deviations per participant after correcting for multiple comparisons (Supplementary Table 10-12 and 16-18).

**Figure 4.**
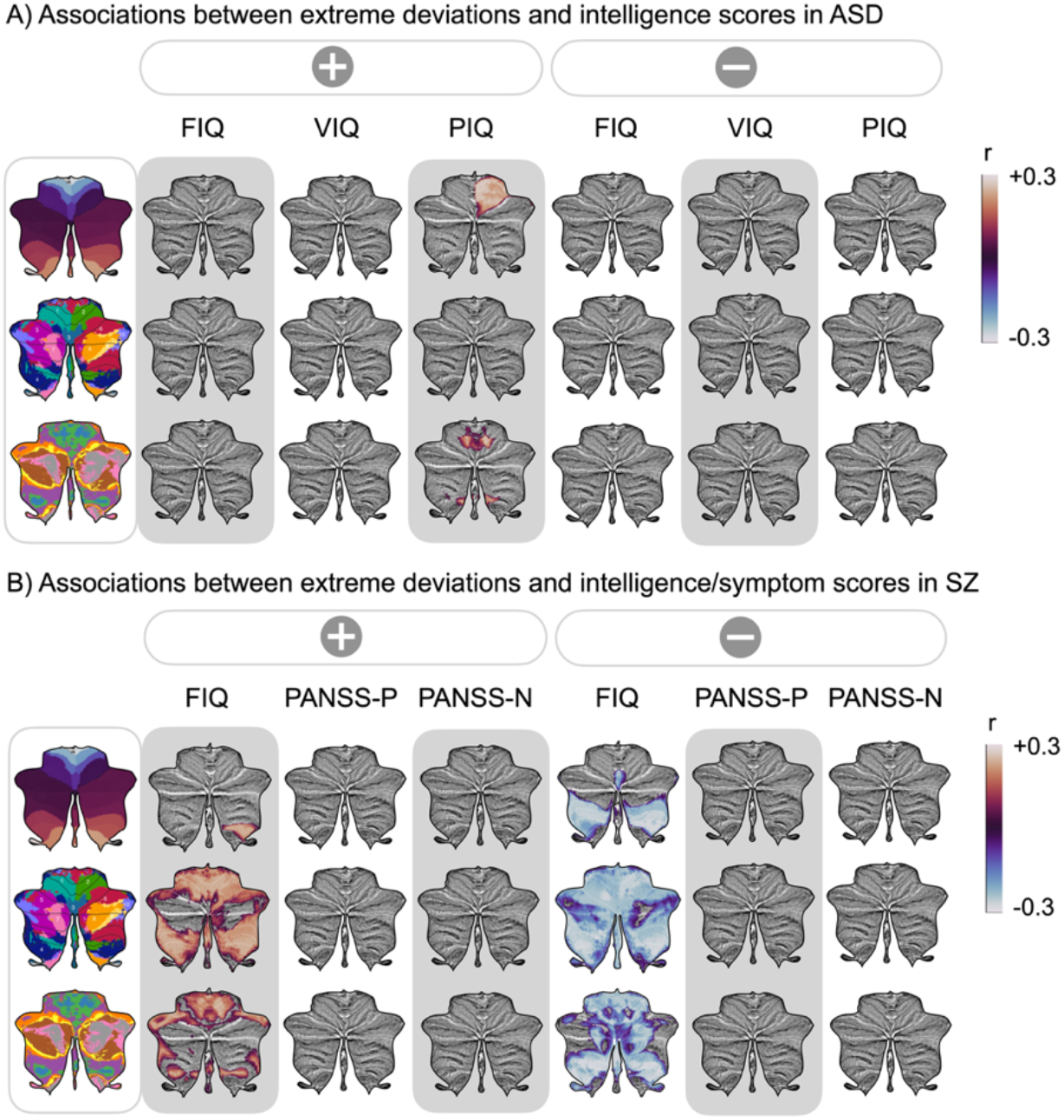
When applied to different atlases, significant correlations were observed between the percentage of extreme deviations per participant and IQ scores. The panels (A-B) show significant correlations between extreme positive (left) or negative (right) deviations per participant and intelligence or symptom scores mapped onto three atlases. Panel (A) displays significant correlations between performance intelligence scores (PIQ) and the percentage of extreme positive deviations per participant in autism spectrum disorder (ASD). (B) In schizophrenia (SZ), positive associations are shown in percentage of extreme positive deviation while negative associations in percentage of extreme negative deviations with full-scale IQ (FIQ).

## Discussion

Leveraging brain MRI data from > 54k individuals, we chart cerebellar lifespan trajectories with voxel-wise precision using normative modelling. These models are estimated across 132 scanning sites and contribute to the field by: i) providing > 143k normative models of the cerebellar lobules and voxels across the lifespan, ii) confirming and extending previous reports of altered cerebellar structure in various mental and neurological illnesses, iii) demonstrating considerable structural cerebellar heterogeneity within all clinical groups, iv) coupling normative models with existing cerebellar atlases to enhance interpretability and v) demonstrating functional significance of extreme deviations in terms of associations with measures of cognitive function. Conditions marked by cognitive deficits, like schizophrenia, ASD, and dementia, exhibit cerebellar differences, although not universally across all individuals with such illnesses. The substantial individual differences even within the same diagnostic groups underscore the multifaceted role of the cerebellum in these clinical phenotypes and highlight the need for fine grained analytical procedures at scale.

As prolonged developmental and aging windows may render the cerebellum susceptible to cellular, morphological and circuit abnormalities^28^, understanding normal and abnormal development of the cerebellum is a major research priority. The lifespan normative models developed here allow us to combine datasets and perform analyses in reference to a common population cohort, making further structural investigations of the cerebellum more comparable. Our results are generally in line with previous reports on cerebellar aging^29,30^ showing a rapid growth of most cerebellar regions during childhood (with volumes typically peaking in late adolescence), followed by a more gradual decline In addition to boosting the sample size relative to these previous studies about 10-fold, we also increase the spatial precision by analysing lifespan trajectories at voxel-wise spatial resolution. This allows for a more fine-grained understanding of cerebellar structure and more precise delineation of functional units. Importantly, these voxel-wise models can readily be integrated with any existing or future cerebellar atlas, and may thus be sensitive to structural deviations that do not necessarily align well with traditional anatomical borders (i.e., lobules)^31^.

Group comparisons using model-derived z-scores confirmed previous reports of altered cerebellar morphology in certain mental and neurological illnesses, while normative probability maps revealed high cerebellar heterogeneity within these same illnesses. First, patients with AD exhibited a significant reduction in lobular measures compared to the individuals without diagnosis while voxel-wise did not survive multiple comparison correction. In line with meta-analysis study on grey matter loss^9^, group effects were particularly pronounced in the Right Crus I which have previously been associated with cognitive processing^32^. A recent study^33^ reported that cerebellum volume is associated with cognitive decline in individuals with MCI but not in those with AD. Conversely, in MCI, the presence of extreme bidirectional deviations in the cerebellum might suggest a possible compensatory mechanism in some individuals and decline in others during the initial phase of the disease^34^, potentially serving as a cognitive reserve^35^. Such “cerebellar reserve”^36^, might thus mitigate some of the cognitive decline associated with neurodegeneration through compensatory reorganization. Our findings further suggest that only a proportion of all diagnosed individuals with AD exhibit extreme negative normative deviations in the cerebellum. These findings highlight the heterogeneity in phenotypes and pathophysiology in AD^37^ and the importance of looking into the variability of the disease.

In line with a previous mega-analysis^10^, we observed significantly lower normative model z-scores (indicating lower volumes) in patients with SZ relative to individuals without a diagnosis. Compared to individuals without diagnosis, some patients with SZ showed evidence of smaller regional volumes in reporting small but reliable reduction in cerebellar volume in SZ, particularly in areas associated with high-level cognitive function. Across atlases, our analyses revealed extreme negative deviations primarily in the posterior regions. Recent meta-analysis study^38^ found brain-predicted age difference in SZ by average of +3.55 years compared to the individuals without diagnosis. In this study, findings of SZ stand out as brain age study^39^ display more pronounced changes in full brain and cerebellar subcortical in comparison to AD. Moreover, disruptions in cerebello-thalamo-cortical circuit may lead to impairment in synchrony of mental processes, possibly generating symptoms of schizophrenia^40^.

The present results are in line with previous findings of substantial heterogeneity in ASD^13,41,42^. We observed medium effects in percentage of negative deviations associated with action observation in task-based atlas and dorsal attention network in resting-state atlas. This is of interest as individuals with ASD frequently report difficulties with social interaction and restricted, repetitive behaviour^43^. D’Mello and colleagues^11^ reported that reduced regional and lobular grey matter volumes in right VII (Crus I/II) correlated with the severity of social, communication and repetitive behaviours, based on ADOS scores. While we observe some nominally significant associations with ADOS scores om the current study, these did not survive correction for multiple comparisons.

Previous studies have established an association between intelligence and larger brain size or greater grey matter volume^44^ as well as total brain volume^45^. Likewise, cerebellar morphology has been associated with cognitive ability and psychiatric symptoms^46^. In patients with SZ, weak yet significant positive and negative associations were found with FIQ across the cerebellum, clearly showing that positive deviations are associated with higher scores and negative deviations with lower scores. In ASD, higher performance intelligence scores were associated with positive deviation in the regions related to Right IV-VI in anatomical atlas and somatomotor network (network 4) in task-based atlas.

While our work represents one of the largest investigations into the cerebellar structures and heterogeneity at the level of the individual, and provides a resource for researchers for future investigations, some limitations require consideration. First, we lack coherent and detailed behavioural, cognitive, genetic, phenotypic, and medical history information for both individuals without diagnosis and clinical samples. This is usually the case when combining information across multiple datasets from different projects as documentation and assessment varies which may preclude certain associations from being tested. Additionally, the coverages of lifespan normative models in the very young age and in age range of 30 to 40 were relatively low. Moreover, our data primarily represents western populations, potentially limiting its generalizability to other populations. Due to its location in the posterior fossa, its intricate arrangement, and motions artifacts, the cerebellum has posed challenges in imaging studies and therefore inferior regions of the cerebellum may result in poorer model fits. The cerebellar topography shows substantial individual differences^47^, and therefore does not perfectly align with existing “average” atlases. Therefore, there are limitations to the interpretation of functional implications. The resource established here can be used in future studies, to further elucidate the complex aetiology of mental and neurological illnesses. By including comprehensive longitudinal datasets or dividing the heterogeneous clinical population into subtypes, we can aim to enhance biomarker development using cognitive and behavioural measures^21,48–50^.

## Conclusion

We report the largest multi-site investigation of heterochronic development and aging of the cerebellum at both voxel-wise and lobular spatial precision. Through normative modelling, we observe individualized patterns of deviation across five different mental and neurological illnesses. Several clinical phenotypes exhibited negative deviations at the group level, but with notable individual differences even within the same clinical groups. Overall, this study charts cerebellar morphology across the lifespan, provides evidence for the differential involvement of the cerebellum across brain illness, and links extreme deviations from population norms to cognitive function.

## Online Methods

### Population data

The cohort of individuals without diagnoses was obtained from 132 scanning sites, including various studies such as ABIDE, ADHD200, AOMIC ID1000, Beijing Enhanced, CAMCAN, CoRR, DLBS, DS000119, DS000202, DS000222, Fcon1000, HBN, HCP, MPI Lemon, NKI-Rockland, OASIS-3, PING, SALD, SLIM, and UK Biobank. Data used in the preparation of this article were obtained from the Alzheimer’s Disease Neuroimaging Initiative (ADNI) database (adni.loni.usc.edu). The ADNI was launched in 2003 as a public-private partnership, led by Principal Investigator Michael W. Weiner, MD. The primary goal of ADNI has been to test whether serial magnetic resonance imaging (MRI), positron emission tomography (PET), other biological markers, and clinical and neuropsychological assessment can be combined to measure the progression of mild cognitive impairment (MCI) and early Alzheimer’s disease (AD). Further details about each study can be found in the associated publications. (Supplementary Table 1). The total number of participants in the individuals without diagnosis population was 54102 (53% females), and the clinical set was 1757, encompassing > 56k in total. The age range spanned from 3 to 85 years (Figure 1). Detailed descriptions of each site, including sample size, mean age, standard deviation, and sex ratio, can be found in Supplementary Table 2. If longitudinal scans were available in the studies, only the baseline scans were used. Participants who had withdrawn from the studies or had missing demographic information and T1-weighted MRI data were excluded from the analyses.

### Clinical data

As for the clinical datasets, we combined data from ABIDE, ADNI, AIBL, DEMGEN, and TOP (Figure 1). Apart from the requirement of having available clinical diagnoses, clinical groups with more than 100 participants were included in the study. Among the clinical groups, we selected AD (Alzheimer’s disease), ASD (autism spectrum disorder), BD (bipolar disorder), MCI (mild cognitive impairment), and SZ (schizophrenia) as the clinical cohorts.

### Lobular-level processing

The T1-weighted images were skull stripped using the FreeSurfer 5.3 auto-recon pipeline^51^ and reoriented to the standard FSL orientation using the *fslreorient2std*^52^. The linear registration was performed using *flirt*^53^, which utilized linear interpolation and the default 1 mm FSL template (version 6.0). The borders were cropped in [6:173, 2:214, 0:160] coordinates to minimize size while retaining the complete volume. Lastly, the voxel intensity values were normalized to the range of [0,1], adjusting the intensity values of each voxel to a standardized scale.

In our study, we utilized the ACAPULCO algorithm^29^, which is a state-of-the-art cerebellum parcellation algorithm based on convolutional neural networks. This algorithm is part of the ENIGMA Cerebellum Volumetric Pipeline and provides speedy and accurate quantitative in-vivo regional assessment of the cerebellum at the highest fidelity^54^. We used pre-processed T1-weighted image for better quality control and alignment. The inhomogeneity of the images was corrected using the N4^55^ and registered to the 1mm isotropic ICBM 2009c template in MNI space using the ANTs registration suite^56^. The ACAPULCO algorithm was trained using 15 expert manual delineations of an adult cohort^54^. It performs per-voxel labelling of the cerebellum and applies post-processing to remove isolated pieces, ensuring accurate segmentation. The algorithm divides the cerebellum into 28 cerebellar lobules, including bilateral Lobules I–VI; Crus I and II; Lobules VIIB, VIIIA, VIIIB, and IX-X; Vermis VI, VII, VIII, IX, and X; and Corpus Medullare (CM). It also calculates the volume (mm^3^) of each region. The automated quality control generates segmented images for each participant, and we removed the extreme outliers where the number of lobules exceeded a certain threshold (e.g., n > 2).

### Voxel-level processing

In our study, we utilized the SUIT (Spatially Unbiased Infratentorial Toolbox) toolbox to perform segmentation of cerebellar grey and white matter voxel-based morphometry (VBM) maps. This segmentation process involved using the outputs from ACAPULCO, which include the N4 bias-corrected and MNI-aligned T1 image^57,58^, as well as an averaged mask derived from randomly selected 300 individuals without diagnosis. The ACAPULCO mask plays a crucial role in correcting and refining overinclusion errors in the segmentation process due to variations in the segmentation algorithm. Following segmentation, the grey matter maps were normalized and resliced to align with a standardized space. This normalization step ensures that the data can be compared across different individuals and studies. Additionally, the grey matter maps were modulated by the Jacobian to preserve the value of each voxel in proportion to its original volume. This modulation accounts for individual differences in brain size and helps to retain the relative intensity values within the mapped brain regions. By using the SUIT toolbox and incorporating the ACAPULCO outputs, we were able to obtain accurate and spatially unbiased segmentation of cerebellar grey and white matter, enabling further analysis and comparison of VBM maps within and across individuals.

### Normative modelling

We split our individuals without diagnosis sample into training and test sets based on scanning site, sex, and age. This split is important to account for the potential confounding effects of MRI scanners on the data^17,59,60^. The individuals without diagnosis were first stratified based on the scanning sites. This ensures that the training and testing datasets include a representative distribution of participants from each location. To achieve this, we evenly split the control participants from each scanning site between the training and testing sets. A minimum requirement of 5 participants from the same scanners sites was required. However, only the test set consisted of diagnostic groups (e.g., AD, ASD, BD) with minimum of 100 participants. This criterion ensures that there is an adequate number of participants in each diagnostic group to provide reliable statistical analyses. By employing this stratification approach, we aimed to create balanced and representative training and testing datasets that account for MRI scanners and sex, while also including enough participants in each diagnostic group.

We used the PCNtoolkit package (version 0.24)^15,61^ in Python 3.8 to estimate a normative model for predicting regional cerebellar volumes and voxel-wise intensity based on sex and age, while correcting for scanning site. Results that deviated more than 5 standard of deviation were imputed by the mean. We employed Bayesian Linear Regression (BLR) with likelihood warping approach^62^, specifically using the ’sinarcsinsh’ transformation^60,63^. This approach is well-suited for handling non-linear basis functions and non-Gaussian predictive distributions for large datasets as well as correcting for outer centiles. A detailed mathematical description on BLR for normative modelling can be found in the following paper Fraza et al. (2021)^60^. To account for scanner effects, we treated the scanning site as a fixed effect in our analysis^16,64^. This approach has been shown to yield relatively good performance, as demonstrated in previous work^16^. To assess how each participant’s (*i*) deviate from the individuals without diagnosis pattern at each lobule or voxel (*j*) in the cerebellum, we calculated the z-score:

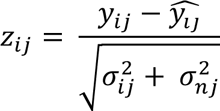

The computation of the z-score includes predicted mean 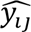(lobule or voxel), true response *y*_*ij*_, predicted variance *σ*_*ij*_ and normative variance *σ*_*nj*_. For model fit, the normative model provided point estimates and evaluation metrics, including explained variance, mean squared log-loss, skew, and kurtosis^63^. These evaluation metrics were computed in the test set that did not include any clinical groups. To determine participants with extreme deviations, we set a threshold at *z* > |1.96|, corresponding to the 95% confidence interval. For instance, deviations with z-scores greater than 1.96 were identified as extreme positive deviations, indicating significantly increased volume compared to the control pattern and vice versa for extreme negative deviations.

### Group comparisons

We performed classical nonparametric test on the z-scores of clinical cohorts and individuals without diagnosis. To assess the statistical significance, we performed Mann-Whitney U-tests^65^, a non-parametric test that is suitable for comparing two independent samples that are not normally distributed. To account for multiple comparisons, the resulting p-values were corrected, using Bonferroni correction^66^ and calculated rank biserial correlation to see its effect.

### Atlas-based analyses

Additionally, the normative model can be applied to various research questions and is compatible with existing atlases by using registration methods such as FSL *flirt* and *fnirt* ^53,67^. This makes it an attractive versatile tool that can be utilized in different studies and across different brain regions. By mapping the deviations onto specific anatomical regions, such as the 28 cerebellar regions, King’s 10 regions of interest from the multi-domain task battery (MDTB)^31^, and 17 regions of interest from resting-state connectivity^68,69^, we gain insights into the specific areas where deviations occur. We separately calculated the percentage of extreme positive and negative deviations for each participant in the regions of interest in reference to the existing atlases. We divided this by the size of the region and multiplied the resulting proportion by 100 (i.e., deriving a percentage of extreme positive or negative deviations per region). To compare the extreme deviations observed in different cohorts to the individuals without diagnosis group, we used Mann-Whitney U-tests and calculated the rank biserial correlation (r) for significant results.

To investigate potential associations between measured intelligence and symptom scores and clinical cohorts, Spearman correlation analyses were performed using voxel-wise extreme deviation scores that were mapped onto the existing atlases. The Spearman correlation coefficient is used to quantify the strength and direction of the association between the variables, allowing for the examination of potential relationship (see Supplementary Methods for tests used). Only correlations with a corrected p-value below 0.05 (p < 0.05) were considered statistically significant and reported.

### Data availability

In this study we used brain imaging from ABIDE, ADHD200, AOMIC ID1000, Beijing Enhanced, CAMCAN, CoRR, DLBS, DS000119, DS000202, DS000222, Fcon1000, HBN, HCP, MPI Lemon, NKI-Rockland, OASIS-3, PING, SALD, SLIM and UK Biobank, ADNI, AIBL, DEMGEN, PNC, and TOP. The ROI models from this work are available on via PCNportal^70^: https://pcnportal.dccn.nl/.

### Code availability

All code used in this work is publicly available at FreeSurfer (https://surfer.nmr.mgh.harvard.edu), FSL (https://fsl.fmrib.ox.ac.uk/fsl/fslwiki/FslInstallation), ACAPULCO (https://gitlab.com/shuohan/acapulco), and SUIT (https://github.com/jdiedrichsen/suit). Code for normative model is available as open-source python package, Predictive Clinical Neuroscience (PCN) toolkit (https://github.com/amarquand/PCNtoolkit). Further codes are available on https://github.com/milinkim/mapping_cerebellar_hetereogeneity.

## Acknowledgements

Data collection and sharing for this project was funded by the Alzheimer’s Disease Neuroimaging Initiative (ADNI) (National Institutes of Health Grant U01 AG024904) and DOD ADNI (Department of Defense award number W81XWH-12-2-0012). ADNI is funded by the National Institute on Aging, the National Institute of Biomedical Imaging and Bioengineering, and through generous contributions from the following: AbbVie, Alzheimer’s Association; Alzheimer’s Drug Discovery Foundation; Araclon Biotech; BioClinica, Inc.; Biogen; Bristol-Myers Squibb Company; CereSpir, Inc.; Cogstate; Eisai Inc.; Elan Pharmaceuticals, Inc.; Eli Lilly and Company; EuroImmun; F. Hoffmann-La Roche Ltd and its affiliated company Genentech, Inc.; Fujirebio; GE Healthcare; IXICO Ltd.; Janssen Alzheimer Immunotherapy Research & Development, LLC.; Johnson & Johnson Pharmaceutical Research & Development LLC.; Lumosity; Lundbeck; Merck & Co., Inc.; Meso Scale Diagnostics, LLC.; NeuroRx Research; Neurotrack Technologies; Novartis Pharmaceuticals Corporation; Pfizer Inc.; Piramal Imaging; Servier; Takeda Pharmaceutical Company; and Transition Therapeutics. The Canadian Institutes of Health Research is providing funds to support ADNI clinical sites in Canada. Private sector contributions are facilitated by the Foundation for the National Institutes of Health (www.fnih.org). The grantee organization is the Northern California Institute for Research and Education, and the study is coordinated by the Alzheimer’s Therapeutic Research Institute at the University of Southern California. ADNI data are disseminated by the Laboratory for Neuro Imaging at the University of Southern California. Also, data used in preparation of this article were obtained from the Australian Imaging Biomarkers and Lifestyle Study of Ageing (AIBL) databases (adni.loni.usc.edu), and the Pediatric Imaging, Neurocognition and Genetics (PING) study database (chd.ucsd.edu/research/ping-study.html, now shared through the NIMH Data Archive (NDA). This publication is solely the responsibility of the authors and does not necessarily represent the views of the National Institutes of Health or PING investigators. The authors of this manuscript gratefully acknowledge the following funding bodies. TW acknowledges funding by the German Research Foundation (DFG; Projectnumber: 513851350) as well as starting funding from the faculty of medicine at the University of Tübingen.

The European Research Council under the European Union’s Horizon 2020 research and Innovation program (ERC StG, Grant 802998), the Research Council of Norway (300767, 324499), the South-Eastern Norway Regional Health Authority (2019101). We performed this work on the *Services for sensitive data* (TSD), University of Oslo, Norway, with resources provided by UNINETT Sigma2 - the National Infrastructure for High-Performance Computing and Data Storage in Norway. We want to acknowledge the Norwegian registry of persons assessed for cognitive symptoms (NorCog), for providing access to patient data. We conducted this research using the UK Biobank Resource under Application Number 27412.

## Authorship Contributions

T.M., T.W., and M.K. originally conceived of the project. M.K., T.W., T.M., and E.L. performed the analyses. M.K. wrote the initial draft of the manuscript. O.A., G.R., K.P., G.S., N.E.S., O.B.S., A.F.M., C.F.B., T.U., and L.W. contributed to data curation. All authors discussed the results and contributed to the final manuscript.

## Ethics declarations

The authors declare no competing interests.

