## Supplementary File for "Mapping Cerebellar Anatomical Heterogeneity in Mental and Neurological Illnesses"

*Supplementary Information*

**Kim et al.**

### Supplementary Methods

#### *Quality control*

We executed quality control by running the ENIGMA Cerebellum Volumetric Pipeline QC Scripts with Singularity. The output of the quality control, "QC_Images.html", provides visual representations of the ACAPULCO segmented images in coronal, sagittal and transverse sections. These images allow for a visual inspection of the segmentation quality and help identify any mis-segmentations or abnormalities in the cerebellar lobules. In addition to the visual assessment, the QC pipeline also provides quantitative and visual information about the cerebellum volumetrics of the segmented regions. This includes measures such as volume, outliers, and box plots of the outliers. These metrics can be used to identify extreme outliers in terms of the number of lobules, indicating potential segmentation errors or other anomalies. By incorporating the QC pipeline into our analysis, we were able to identify and address any mis-segmentations or outliers, improving the reliability and accuracy of our results.

#### *Atlases*

The complex organization of the cerebellum suggests that different subregions of the cerebellum play specialized roles in various cognitive and motor functions. Stoodley and Schmahmann[1] showed functions based on lobules where lobule V/adjacent VI/lobule VIII were activated by sensorimotor tasks, lobule VIIIA/B for motor activation, VIIIB for somatosensory activation, posterior lobes for higher level tasks, VI and Crus I for language and verbal working memory, lobule VI for spatial tasks, VI/Crus I/VIIB for executive functions, and VI/Crus I/medial VII for emotional processing.

However, the functional subregions of the cerebellum do not align with its lobular boundaries, as demonstrated in the MDTB map[2], where regions 1 and 2 are associated with hand movements, 3 with visual memory, 4 with attention, regions 5 and 6 with working memory, regions 7 and 8 to narrative comprehension, regions 8 and 9 to language functions and region 10 to autobiographical recall. These functional mappings illustrate that the boundaries of cerebellar functions do not strictly adhere to the anatomical lobular divisions. Instead, functional boundaries cross lobules, indicating a more nuanced organization of cerebellar functions that may cut across traditional anatomical landmarks.

The work by Buckner et al. [3,4] depicted the complex interconnectivity between the cerebellum and the and the cerebral cortex during resting state. The resting state networks exhibits common functional activations even in the absence of a specific task. By assigning cerebellar voxels to distinct cerebral networks based on their highest temporal correlation, the study revealed the rich functional connections of the cerebellum with various regions of the cortex. This indicates that the cerebellum is involved in ongoing functional interactions with the cerebral cortex that contribute to overall brain functioning. The global topographic organization of the cerebellum, as revealed by the study, shows specific patterns of connectivity between different cerebellar lobules and cortical regions. For instance, lobule VIII and the anterior lobe exhibit connectivity with cortical and premotor areas belonging to networks 3, 4, and 7. Lobule VII and IX, on the other hand, demonstrate connectivity with prefrontal and parietal association areas within networks 8, 12, 13, 14, 16, and 17. The regions for the 17 networks are N1: Visual A, N2: Visual B; Network 3: Somatomotor A; N4: Somatomotor B; N5: Dorsal Attention A; N6: Dorsal Attention B; N7: Salience/ Ventral Attention A; N8: Salience/ Ventral Attention B; N9: Limbic B; N10: Limbic A; N11: Control A; N12: Control B; N13: Control C; N14: Default A; N15: Default B; N16: Default C; N17: Temporal Parietal. Understanding the resting state functional connectivity of the cerebellum provides valuable insights into its contributions to overall brain function and its involvement in both task-based and intrinsic functional networks.

#### *Cognition and Symptom Scores*

The ASD scores examined in the correlation analyses included measures (communication, reciprocal social interaction, and stereotyped behaviours) from testing batteries that calculated Autism Diagnostic Observation Schedule (ADOS)[5] and full-scale IQ (FIQ), verbal IQ (VIQ) and performance IQ (PIQ) from ABIDE dataset. Positive and Negative Syndrome Scale (PANSS)[6] examines the severity of patients with SZ which was divided into positive and negative scale from Oslo TOP dataset. There were some participants with missing cognition and symptom scores resulting in number of ASD of 236 and SZ of 269.

### List of Supplementary Figures

1. Evaluation metrics of lobular and voxel-wise growth charts
2. Cerebellar lobular growth charts
3. Cerebellar voxel-wise growth charts
4. Group based nonparametric tests in lobular cerebellar maps and individual level extreme deviations
5. Group based nonparametric tests in voxel-wise and lobular maps

### List of Supplementary Tables

1. Sources of all the studies used in the study
2. Overview of the dataset
3. Group based nonparametric tests of cerebellar lobules
4. Anatomical atlas based case-control analyses of extreme deviations
5. Task atlas based case-control analyses of extreme deviations
6. Resting-state atlas based case-control analyses of extreme deviations
7. Association of extreme deviations and IQ in ASD using anatomical atlas
8. Association of extreme deviations and IQ in ASD using task-based atlas
9. Association of extreme deviations and IQ in ASD using resting-state atlas
10. Association of extreme deviations and ADOS in ASD using anatomical atlas
11. Association of extreme deviations and ADOS in ASD using task-based atlas
12. Association of extreme deviations and ADOS in ASD using resting-state atlas
13. Association of extreme deviations and IQ in SZ using anatomical atlas

14. Association of extreme deviations and IQ in SZ using task-based atlas
15. Association of extreme deviations and IQ in SZ using resting-state atlas
16. Association of extreme deviations and PANSS in SZ using anatomical atlas
17. Association of extreme deviations and PANSS in SZ using task-based atlas
18. Association of extreme deviations and PANSS in SZ using resting-state atlas

### Supplementary Figures

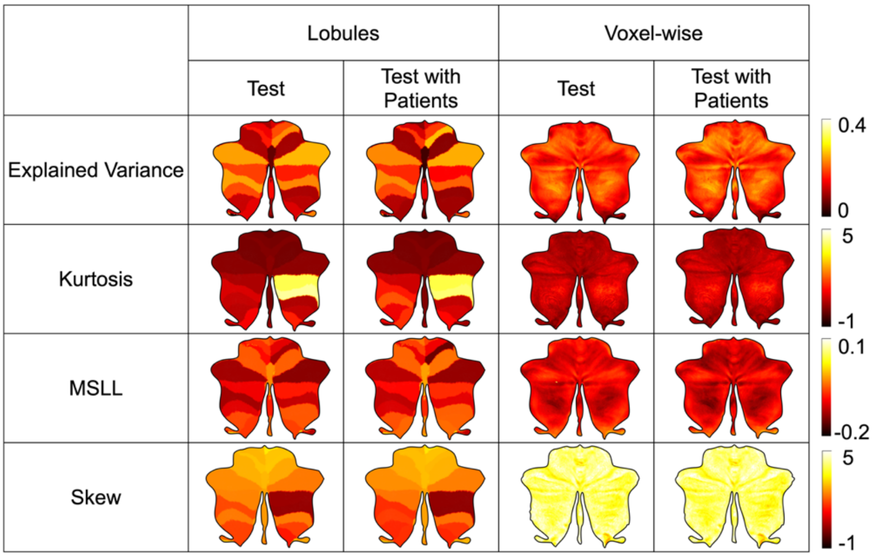


**Supplementary Figure 1.** Evaluation metrics across test set and test set with clinical sets show a good fit of the model in central tendency and variance (explained variance, kurtosis) and shape of the model (MSLL and skew).

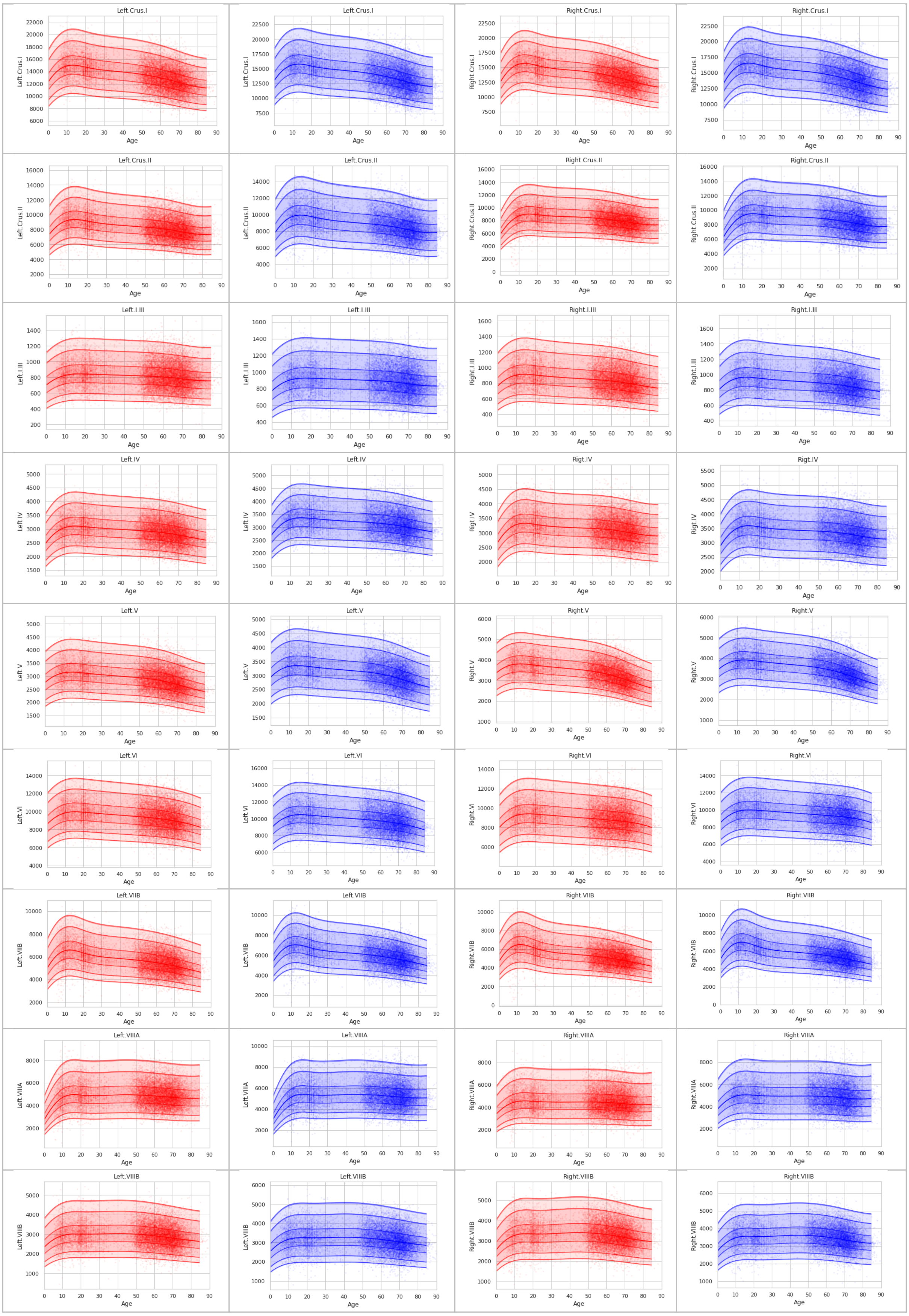
Left.Crus.I
Left.Crus.I
Right.Crus.I
Right.Crus.I
Left.Crus.II
Left.Crus.II
Right.Crus.II
Right.Crus.II
Left.I.III
Left.I.III
Right.I.III
Right.I.III
Left.IV
Left.IV
Rigt.IV
Rigt.IV
Left.V
Left.V
Right.V
Right.V
Left.VI
Left.VI
Right.VI
Right.VI
Left.VIIB
Left.VIIB
Right.VIIB
Right.VIIB
Left.VIIIA
Left.VIIIA
Right.VIIIA
Right.VIIIA
Left.VIIIB
Left.VIIIB
Right.VIIIB
Right.VIIIB
Age

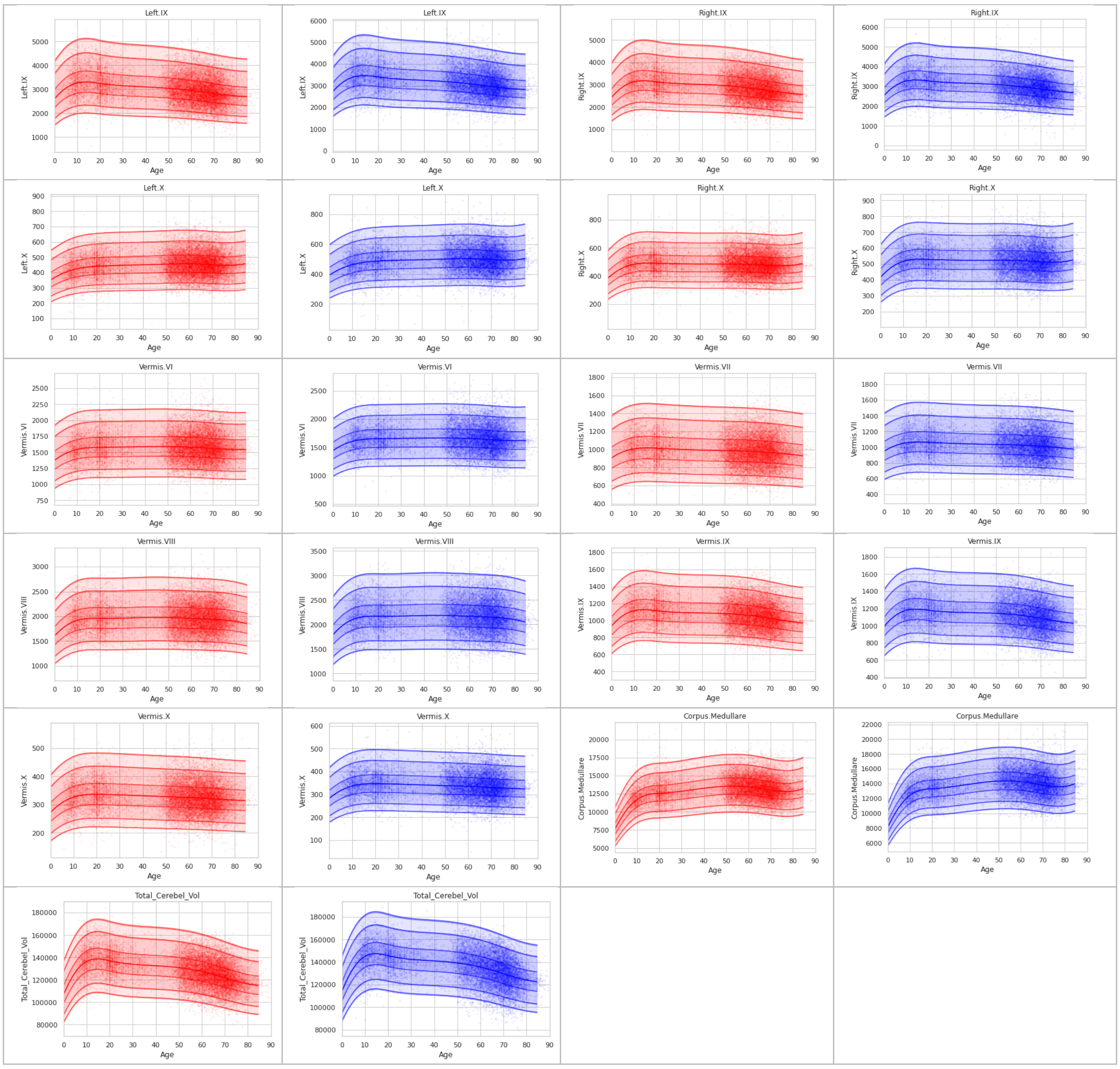


**Supplementary Figure 2. Cerebellar lobular growth charts.**

Total of 28 anatomical regions of the cerebellum are mapped throughout the lifespan, including total cerebellum volume.

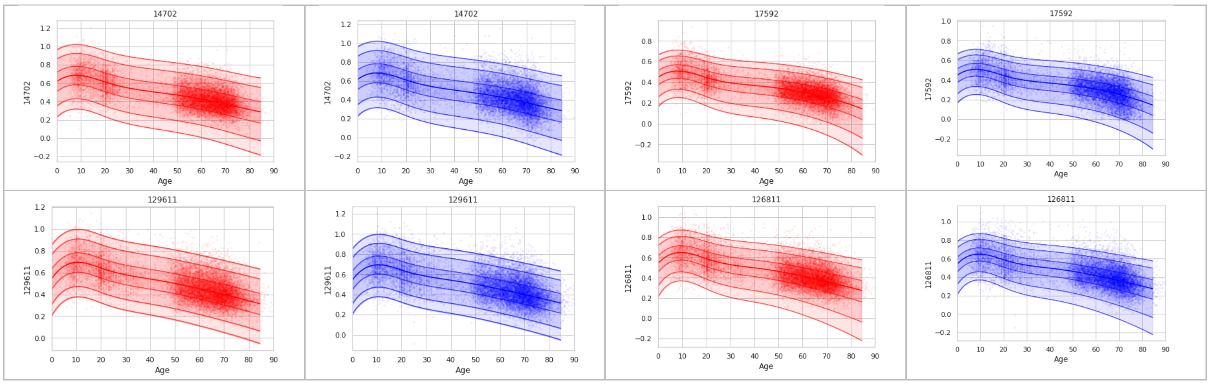


**Supplementary Figure 3. Cerebellar voxel-wise growth charts.**

This shows selected trajectories out of 143k voxels that were mapped.

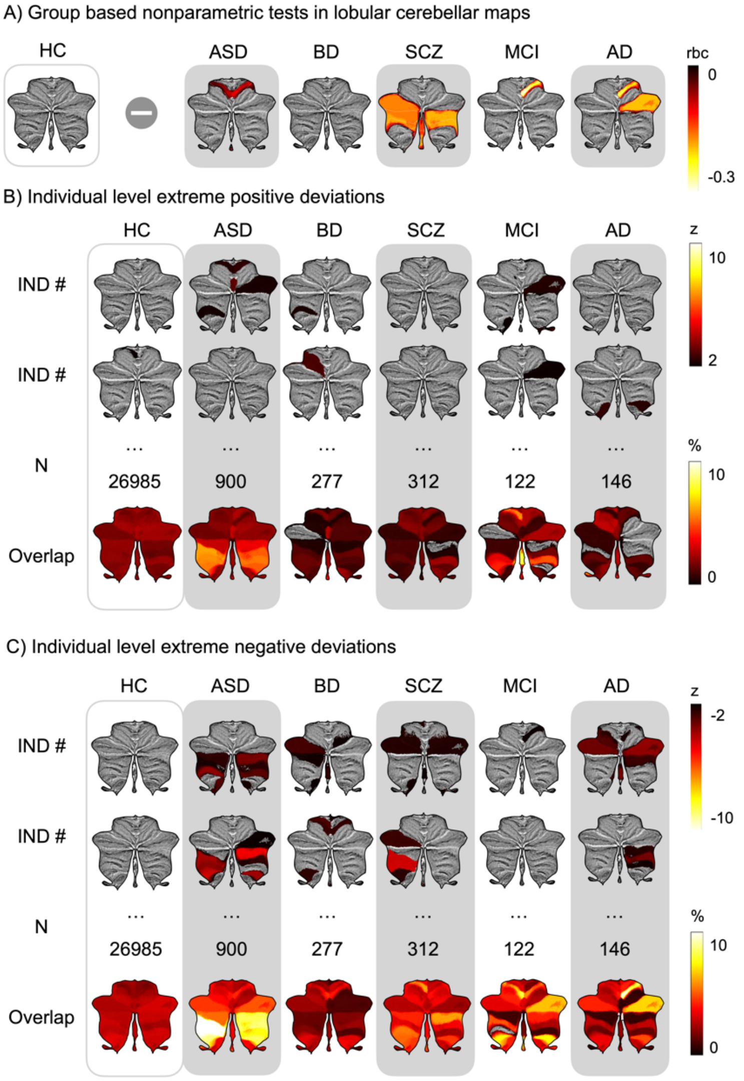


**Supplementary Figure 4. Group based nonparametric tests in lobular cerebellar maps and individual level extreme deviations**

(A) The rank differences in the z-score maps have been adjusted for multiple comparisons using Bonferroni correction (corrected $p < 0.05$). The scale indicates Mann-Whitney U-test rank biserial correlation (rbc) reflecting the statistical significance of the findings. (B) The overlap maps visually represent these extreme deviations in percentage, highlighting the regions in which the deviations occurred. The clinical groups displayed a significantly higher occurrence of extreme deviations, even in cases where the corrected group comparisons did not differ significantly.

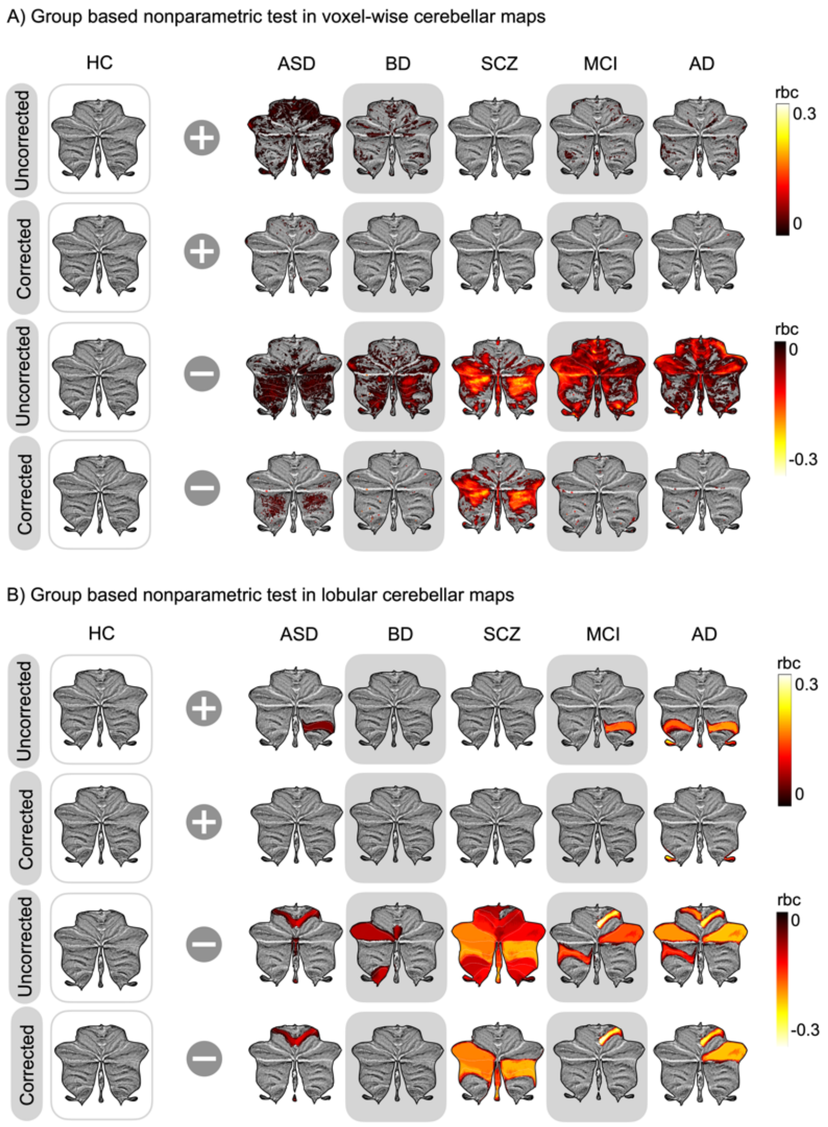


**Supplementary Figure 5. Group based nonparametric tests in voxel and lobular cerebellar maps**

(A-D) The rank differences in the z-score maps shows both uncorrected and corrected for multiple comparisons using Bonferroni correction (corrected $p < 0.05$) in voxel-wise and lobular maps. The scale indicates Mann-Whitney U-test rank biserial correlation (rbc) reflecting the statistical significance of the findings. The uncorrected comparison show that the likelihood of the clinical groups decreases when z-score increases in more regions compared to the corrected comparisons.

### Supplementary Tables

#### Supplementary Table 1. Sources of all the studies used in the study

| Datasets | Sources | Comments | References |
|---|---|---|---|
| Autism Brain Imaging Dataset Exchange | http://fcon_1000.projects.nitrc.org/ | Primary support for the work by Adriana Di Martino was provided by the NIMH (K23MH087770) and the Leon Levy Foundation. Primary support for the work by Michael P. Milham and the INDI team was provided by gifts from Joseph P. Healy and the Stavros Niarchos Foundation to the Child Mind Institute, as well as by an NIMH award to MPM (R03MH096321). | [10] |
| Autism Brain Imaging Dataset Exchange II | http://fcon_1000.projects.nitrc.org/ | Primary support for the work by Adriana Di Martino and her team was provided by the National Institute of Mental Health (NIMH 5R21MH107045). Primary support for the work by Michael P. Milham and his team provided by the National Institute of Mental Health (NIMH 5R21MH107045); Nathan S. Kline Institute of Psychiatric Research). Additional Support was provided by gifts from Joseph P. Healey, Phyllis Green and Randolph Cowen to the Child Mind Institute. | [11] |
| ADHD200 | http://fcon_1000.projects.nitrc.org/ | F. Xavier Castellanos, David Kennedy, Michael Milham, and Stewart Mostofsky are responsible for the initial conception of the ADHD-200 Consortium. Consortium steering committee includes Jan Buitelaar, F. Xavier Castellanos, Dan Dickstein, Damien Fair, David Kennedy, Beatriz Luna, Michael Milham (Project Coordinator), Stewart Mostofsky, and Julie Schweitzer. Data aggregation and organization was coordinated by the INDI team, which included Saroja Bangaru, David Gutman, Maarten Mennes, and Michael Milham. Web infrastructure and data storage were coordinated by Robert Buccigrossi, Albert Crowley, Christian Hasselgrove, David Kennedy, Kimberly Pohland, and Nina Preuss. The ADHD-200 Global Competition Coordinators were Damien Fair (Chair of Selection Committee, Editor in Chief for Global Competition Special issue) and Michael Milham | [12,13] |
| Alzheimer's Disease Neuroimaging Initiative | http://adni.loni.usc.edu/ | The ADNI was launched in 2003 as a public-private partnership, led by Principal Investigator Michael W. Weiner, MD. ADNI consists of 4 waves, the later is still ongoing (ADNI 3). A complete listing of ADNI investigators can be found at http://adni.loni.usc.edu/wpcontent/uploads/how_to_apply/ADNI_Acknowledgement_List.pdf. Data collection and sharing for this project was funded by the Alzheimer's Disease Neuroimaging Initiative (ADNI) (National Institutes of Health Grant U01 AG024904) and DOD ADNI (Department of Defense award number W81XWH-12-2-0012). ADNI is funded by the National Institute on Aging, the National Institute of Biomedical Imaging and Bioengineering, and through generous contributions from the following: AbbVie, Alzheimer's Association; Alzheimer's Drug Discovery Foundation; Araclon Biotech; BioClinica, Inc.; Biogen; Bristol-Myers Squibb Company; CereSpir, Inc.; Cogstate; Eisai Inc.; Elan Pharmaceuticals, Inc.; Eli Lilly and Company; EuroImmun; F. Hoffmann-La Roche Ltd and its affiliated company Genentech, Inc.; Fujirebio; GE Healthcare; IXICO Ltd.;Janssen Alzheimer Immunotherapy Research & Development, LLC.; Johnson & Johnson Pharmaceutical Research & Development LLC.; Lumosity; Lundbeck; Merck & Co., Inc.;Meso Scale Diagnostics, LLC.; NeuroRx Research; Neurotrack Technologies; Novartis Pharmaceuticals Corporation; Pfizer Inc.; Piramal Imaging; Servier; Takeda Pharmaceutical Company; and Transition Therapeutics. The Canadian Institutes of Health Research is providing funds to support ADNI clinical sites in Canada. Private sector contributions are facilitated by the Foundation for the National Institutes of Health (www.fnih.org). The grantee organization is the Northern California Institute for Research and Education, and the study is coordinated by the Alzheimer's Therapeutic Research Institute at the University of Southern California. ADNI data are disseminated by the Laboratory for Neuro Imaging at the University of Southern California. | |

| | | | |
|---|---|---|---|
| The Australian Imaging, Behaviour and Lifestyle Flagship Study | https://aibl.csiro.au/ | Australian Imaging Biomarkers and Lifestyle flagship study of ageing (AIBL) was funded by the Commonwealth Scientific and Industrial Research Organisation (CSIRO), which was made available at the ADNI databas (http://www.loni.usc.edu/ADNI). The AIBL researchers contributed data but did not participate in analysis or writing of this report. AIBL researchers are listed at http://www.aibl.csiro.au. Correspondence should be addressed to Christopher Rowe. | 14 |
| The Amsterdam Open MRI Collection | https://nilab-uva.github.io/AOMIC.github.io/ | We thank all research assistants and students who helped collecting the data of the three projects, Jasper Wijnen and Marco Teunisse for advice and guidance with respect to anonymization and GDPR-related concerns, and Jos Bloemers, Sennay Ghebeab, Adriaan Tuiten, Joram van Driel, Christian Olivers, Ilja Sligte, Sara Jahfari, Guido van Wingen, and Suzanne Oosterwijk for help with designing the paradigms, Marcus Spaan for technical support, and Franklin Feingold and Joe Wexler for help with uploading the datasets to Openneuro. | 15 |
| Beijing Normal University Enhanced Sample | http://fcon_1000.projects.nitrc.org/ | Financial support for the data used in this project was provided by a grant from the National Natural Science Foundation of China: 30770594 and a grant from the National High Technology Program of China (863): 2008AA02Z405. | 16,17 |
| Cam-CAN | https://camcan-archive.mrc-cbu.cam.ac.uk/dataaccess/ | Data collection and sharing for this project was provided by the Cambridge Centre for Ageing and Neuroscience (CamCAN). CamCAN funding was provided by the UK Biotechnology and Biological Sciences Research Council (grant number BB/H008217/1), together with support from the UK Medical Research Council and University of Cambridge, UK. | 18,19 |
| Consortium for Reliability and Reproducibility | http://fcon_1000.projects.nitrc.org/ | | 20 |
| Demgen | Authors | Supported by the Norwegian National Advisory Unit on Aging and Health | 21,22 |
| Dallas Lifespan Brain Study | http://fcon_1000.projects.nitrc.org/ | | 23 |
| ds00119 | https://openfmri.org/ | Supported by the National Institutes of Mental Health (NIMH RO1 MH067924). Enami Yasui provided assistance with data collection | 24 |
| ds000202 | https://openfmri.org/ | | 25,26 |
| ds000222 | https://openfmri.org/ | | 27 |
| 1000 Functional Connectomes Classic Sample | http://fcon_1000.projects.nitrc.org/ | Collected at 33 independent sites by J.J. Pekar, S.H. Mostofsky, S, Colcombe, Y.F. Zang, D. Marguiles, R.L. Buckner, M.J Low, B. Rypma, D.J. Madden, A.C. Evans, S.A.R.B. Rombouts, A. Villringer, S.J. Li, C.Sorg, V. Riedel, B. Biswal, M. Hampson, M.P. Milham, F.X. Castellanos, P. Williamson, M. Hoptman, V.J. | |

| | | | |
|---|---|---|---|
| | | Kiviniemi, J. Veijiola, S.M. Smith, C. Mackay, M. Greicius, G. Siegle, K. McMahon, B. Schlaggar, S. Petersen, C.P. Lin, H.S. Mayberg, C.S. Monk, R.D. Seidler, S.J. Peltier | |
| Human Connectome Project | https://www.humanconnectome.org/ | Data were provided [in part] by the Human Connectome Project, MGH-USC Consortium (PIs: Bruce R. Rosen, Arthur W. Toga and Van Wedeen; U01MH093765) funded by the NIH Blueprint Initiative for Neuroscience Research grant; the National Institutes of Health grant P41EB015896; and the Instrumentation Grants S10RR023043, 1S10RR023401, 1S10RR019307. | 28 |
| Healthy Brain Network | http://fcon_1000.projects.nitrc.org/ | | 29 |
| Max Planck Institut Leipzig Mind-Brain-Body Dataset | http://fcon_1000.projects.nitrc.org/ | | 30 |
| Enhanced Nathan Kline Institute - Rockland Sample | http://fcon_1000.projects.nitrc.org/ | Principal support for the enhanced NKI-RS project is provided by the NIMH BRAINS R01MH094639-01 (PI Milham). Funding for key personnel also provided in part by the New York State Office of Mental Health and Research Foundation for Mental Hygiene. Funding for the decompression and augmentation of administrative and phenotypic protocols provided by a grant from the Child Mind Institute (1FDN2012-1). Additional personnel support provided by the Center for the Developing Brain at the Child Mind Institute, as well as NIMH R01MH081218, R01MH083246, and R21MH084126. Project support also provided by the NKI Center for Advanced Brain Imaging (CABI), the Brain Research Foundation, and the Stavros Niarchos Foundation. | 31 |
| Open Access Series of Imaging Studies 3 | http://www.oasis-brains.org/ | Data were provided by OASIS 3: Longitudinal Multimodal Neuroimaging: Principal Investigators: T. Benzinger, D. Marcus, J. Morris; NIH P30 AG066444, P50 AG00561, P30 NS09857781, P01 AG026276, P01 AG003991, R01 AG043434, UL1 TR000448, R01 EB009352. AV-45 doses were provided by Avid Radiopharmaceuticals, a wholly owned subsidiary of Eli Lilly. Supported by grants P50 AG05681, P01 AG03991, R01 AG021910, P50 MH071616, U24 RR021382, R01 MH56584. | 32 |
| Pediatric Imaging, Neurocognition and Genetics | http://pingstudy.ucsd.edu/ | Data used in the preparation of this article were obtained from the Pediatric Imaging, Neurocognition and Genetics (PING) Study database (www.chd.ucsd.edu/research/ping-study.html, now shared through the NIMH Data Archive (NDA)). PING was a multisite, cross-sectional study that recruited more than 1,700 participants aged 3 to 20 years. The study was supported by award number RC2DA029475 from the National Institute on Drug Abuse with additional support for data sharing provided by the Eunice Kennedy Shriver National Institute of Child Health & Human Development under award number R01HD061414. A list of participating sites and study investigators can be found at https://ping-dataportal.ucsd.edu/sharing/Authors10222012.pdf. PING investigators designed and implemented the study and/or provided data but did not necessarily participate in analysis or writing of this report. This publication is solely the responsibility of the authors and does not necessarily represent the views of the National Institutes of Health or PING investigators. | 33 |
| Philadelphia Neurodevelopmental Cohort | https://www.med.upenn.edu/bbl/philadelphianeurodevelopmentalcohort.html | | 34 |
| Southwest University Adult Lifespan Dataset | http://fcon_1000.projects.nitrc.org/ | | 20 |

| | | | |
|---|---|---|---|
| Southwest University Longitudinal Imaging Multimodal Brain Data | http://fcon_1000.projects.nitrc.org/ | Support was provided by grant numbers 31271087; 31470981; 31571137, 31500885, SWU1509383, SWU1509451, cstc2015jcyjA10106, 151023, 2015M572423, 2015M580767, Xm2015037, 14JJD880009 | 35,36 |
| StrokeMRI | Authors | Supported by the Research Council of Norway (249795, 248238), the South-Eastern Norway Regional Health Authority (2014097, 2015044, 2015073, 2016083), and the Norwegian ExtraFoundation for Health and Rehabilitation (2015/FO5146) | 37 |
| TOP | Authors | Supported by several grants from the Research Council of Norway, and the South-Eastern Norway Regional Health Authority | 38–40 |
| UK BioBank | https://www.ukbiobank.ac.uk/ | This research has been conducted using the UK Biobank Resource (access code 27412). | 41,42 |

**Supplementary Table 2**. **Overview of the reference set divided into training, testing and clinical sets shown by site**

| | Train | | | | Test | | | | Clinical | | | |
|---|---|---|---|---|---|---|---|---|---|---|---|---|
| **site** | **N** | **Age (m)** | **Age (std)** | **Sex Ratio** | **N** | **Age (m)** | **Age (std)** | **Sex Ratio** | **N** | **Age (m)** | **Age (std)** | **Sex Ratio** |
| abide2_BNI | 15.0 | 40.0 | 15.58 | 0:1 | 14.0 | 39.14 | 15.11 | 0:1 | 28.0 | 38.04 | 16.07 | 0:1 |
| abide2_EMC | 14.0 | 8.08 | 1.04 | 0.21:0.79 | 13.0 | 8.24 | 0.97 | 0.15:0.85 | 26.0 | 8.12 | 1.16 | 0.19:0.81 |
| abide2_ETH | 12.0 | 23.47 | 4.78 | 0:1 | 12.0 | 24.3 | 4.38 | 0:1 | 13.0 | 20.56 | 3.4 | 1.00 |
| abide2_GU | 27.0 | 10.41 | 1.75 | 0.48:0.52 | 25.0 | 10.4 | 1.63 | 0.48:0.52 | 48.0 | 10.91 | 1.54 | 0.17:0.83 |
| abide2_IP | 17.0 | 23.52 | 12.15 | 0.65:0.35 | 16.0 | 23.74 | 11.62 | 0.69:0.31 | 22.0 | 15.13 | 4.93 | 0.36:0.64 |
| abide2_KKI | 39.0 | 10.36 | 1.29 | 0.44:0.56 | 37.0 | 10.38 | 1.21 | 0.43:0.57 | 19.0 | 10.04 | 1.53 | 0.37:0.63 |
| abide2_NYU_1 | 15.0 | 8.82 | 2.02 | 0.07:0.93 | 15.0 | 10.22 | 4.22 | 0.07:0.93 | 45.0 | 9.94 | 5.86 | 0.11:0.89 |
| abide2_OHSU | 28.0 | 10.32 | 1.66 | 0.54:0.46 | 27.0 | 10.44 | 1.67 | 0.52:0.48 | 37.0 | 11.81 | 2.27 | 0.19:0.81 |
| abide2_OILH | 15.0 | 23.6 | 4.08 | 0.40:0.60 | 13.0 | 23.85 | 3.6 | 0.38:0.62 | 18.0 | 22.11 | 3.86 | 0.17:0.83 |
| abide2_SDSU | 13.0 | 13.29 | 3.21 | 0.08:0.92 | 12.0 | 13.21 | 2.98 | 0.08:0.92 | 32.0 | 12.87 | 3.33 | 0.22:0.78 |
| abide2_STANFORD | 3.0 | 10.56 | 1.74 | 0:1 | 3.0 | 11.51 | 0.95 | 0:1 | 17.0 | 11.15 | 1.13 | 0.06:0.94 |
| abide2_TCD | 10.0 | 15.38 | 3.3 | 0:1 | 10.0 | 15.9 | 3.25 | 0:1 | 21.0 | 14.79 | 3.26 | 0:1 |
| abide2_UCD | 7.0 | 14.52 | 1.77 | 0.29:0.71 | 7.0 | 15.08 | 1.75 | 0.29:0.71 | 18.0 | 14.75 | 1.97 | 0.22:0.78 |
| abide2_UCLA | 9.0 | 9.93 | 2.34 | 0.33:0.67 | 7.0 | 9.46 | 1.99 | 0.29:0.71 | 16.0 | 11.67 | 2.22 | 0.06:0.94 |
| abide2_USM | 9.0 | 23.88 | 8.33 | 0.22:0.78 | 7.0 | 24.1 | 7.7 | 0.14:0.86 | 17.0 | 18.31 | 6.99 | 0.12:0.88 |
| abide2_U_MIA | 4.0 | 10.32 | 3.52 | 0.50:0.50 | 2.0 | 9.1 | 1.56 | 0.50:0.50 | 7.0 | 10.96 | 1.82 | 0.14:0.86 |
| abide_CALTECH | 8.0 | 25.0 | 8.72 | 0.25:0.75 | 8.0 | 29.54 | 12.63 | 0.25:0.75 | 16.0 | 26.32 | 9.57 | 0.25:0.75 |
| abide_CMU | 7.0 | 25.29 | 4.99 | 0.29:0.71 | 6.0 | 28.67 | 6.47 | 0.17:0.83 | 13.0 | 26.0 | 5.92 | 0.15:0.85 |
| abide_KKI | 17.0 | 10.13 | 1.38 | 0.29:0.71 | 15.0 | 10.07 | 1.12 | 0.27:0.73 | 22.0 | 10.01 | 1.45 | 0.18:0.82 |
| abide_LEUVEN_1 | 8.0 | 23.25 | 3.45 | 0:1 | 7.0 | 23.29 | 2.43 | 0:1 | 14.0 | 21.86 | 4.11 | 0:1 |
| abide_LEUVEN_2 | 10.0 | 14.25 | 1.49 | 0.30:0.70 | 9.0 | 14.68 | 1.48 | 0.22:0.78 | 15.0 | 13.92 | 1.31 | 0.20:0.80 |
| abide_LMU | 16.0 | 25.25 | 9.57 | 0.12:0.88 | 16.0 | 27.38 | 10.49 | 0.12:0.88 | 23.0 | 24.96 | 14.14 | 0.13:0.87 |
| abide_NYU | 52.0 | 15.77 | 6.32 | 0.25:0.75 | 51.0 | 15.84 | 6.21 | 0.24:0.76 | 71.0 | 14.31 | 6.65 | 0.13:0.87 |
| abide_OHSU | 8.0 | 10.06 | 1.27 | 0:1 | 7.0 | 10.07 | 0.93 | 0:1 | 13.0 | 11.66 | 2.25 | 0:1 |
| abide_OLIN | 8.0 | 16.0 | 3.7 | 0.12:0.88 | 8.0 | 17.88 | 3.64 | 0.12:0.88 | 18.0 | 16.83 | 3.38 | 0.17:0.83 |
| abide_SBL | 7.0 | 32.43 | 7.79 | 1.00 | 7.0 | 35.0 | 6.11 | 0:1 | 15.0 | 35.0 | 10.43 | 0:1 |
| abide_SDSU | 10.0 | 13.81 | 2.27 | 0.30:0.70 | 9.0 | 14.47 | 1.67 | 0.33:0.67 | 14.0 | 14.72 | 1.76 | 0.07:0.93 |
| abide_STANFORD | 8.0 | 9.71 | 1.72 | 0.25:0.75 | 8.0 | 10.36 | 1.63 | 0.25:0.75 | 16.0 | 10.16 | 1.54 | 0.25:0.75 |
| abide_TRINITY | 12.0 | 17.03 | 3.77 | 0:1 | 12.0 | 17.5 | 3.85 | 0:1 | 24.0 | 17.28 | 3.57 | 0:1 |
| abide_UCLA_1 | 16.0 | 13.08 | 2.18 | 0.12:0.88 | 15.0 | 13.13 | 1.81 | 0.13:0.87 | 41.0 | 13.1 | 2.62 | 0.15:0.85 |
| abide_UCLA_2 | 7.0 | 12.21 | 1.26 | 0.14:0.86 | 6.0 | 12.31 | 1.03 | 0.17:0.83 | 13.0 | 12.72 | 1.87 | 0:1 |
| abide_UM_1 | 27.0 | 14.01 | 3.28 | 0.33:0.67 | 26.0 | 14.2 | 3.05 | 0.31:0.69 | 52.0 | 12.84 | 2.41 | 0.15:0.85 |
| abide_UM_2 | 11.0 | 16.72 | 4.3 | 0.09:0.91 | 9.0 | 17.03 | 3.89 | 1.00 | 13.0 | 14.88 | 1.55 | 0.08:0.92 |
| abide_UPSM | 13.0 | 17.6 | 5.94 | 0.15:0.85 | 13.0 | 19.04 | 6.42 | 0.15:0.85 | 30.0 | 18.93 | 7.2 | 0.13:0.87 |
| abide_USM | 19.0 | 20.7 | 7.2 | 0:1 | 19.0 | 21.79 | 7.94 | 0:1 | 52.0 | 22.87 | 7.73 | 0:1 |
| abide_YALE | 13.0 | 12.51 | 2.97 | 0.31:0.69 | 12.0 | 12.88 | 2.76 | 0.25:0.75 | 28.0 | 12.75 | 3.05 | 0.29:0.71 |
| adhd_Brown | 14.0 | 14.0 | 2.57 | 0.64:0.36 | 12.0 | 14.0 | 2.34 | 0.67:0.33 | | | | |
| adhd_KKI | 34.0 | 9.74 | 1.26 | 0.41:0.59 | 33.0 | 9.76 | 1.23 | 0.39:0.61 | | | | |
| adhd_NYU | 46.0 | 11.91 | 3.1 | 0.54:0.46 | 45.0 | 12.0 | 3.16 | 0.53:0.47 | | | | |
| adhd_NeuroIMAGE | 19.0 | 17.79 | 3.49 | 0.68:0.32 | 18.0 | 18.39 | 3.01 | 0.67:0.33 | | | | |
| adhd_OHSU | 34.0 | 8.79 | 1.37 | 0.56:0.44 | 32.0 | 8.72 | 1.14 | 0.56:0.44 | | | | |
| adhd_Peking_0 | 14.0 | 9.36 | 1.6 | 0.50:0.50 | 13.0 | 9.92 | 2.14 | 0.54:0.46 | | | | |
| adhd_Peking_1 | 31.0 | 10.74 | 1.77 | 0.71:0.29 | 29.0 | 10.66 | 1.61 | 0.72:0.28 | | | | |

| | | | | | | | | | | | | |
|---|---|---|---|---|---|---|---|---|---|---|---|---|
| adhd_Peking_2 | 17.0 | 11.12 | 1.93 | 0.06:0.94 | 15.0 | 11.27 | 1.71 | 0:1 | | | | |
| adhd_Peking_3 | 11.0 | 12.64 | 0.92 | 0:1 | 11.0 | 12.64 | 0.92 | 0:1 | | | | |
| adhd_Pittsburgh | 46.0 | 14.41 | 2.8 | 0.48:0.52 | 46.0 | 14.63 | 2.83 | 0.48:0.52 | | | | |
| adhd_WashU | 31.0 | 10.9 | 3.73 | 0.45:0.55 | 30.0 | 11.17 | 4.07 | 0.47:0.53 | | | | |
| adni_11.0-1.5 | 4.0 | 70.6 | 4.99 | 0.25:0.75 | 3.0 | 74.63 | 9.58 | 0.33:0.67 | 4.0 | 77.0 | 4.4 | 0.50:0.50 |
| adni_116.0-1.5 | 4.0 | 73.55 | 3.05 | 0.75:0.25 | 2.0 | 73.6 | 2.83 | 1.00 | 4.0 | 76.95 | 5.49 | 0.75:0.25 |
| adni_116.0-3.0 | 4.0 | 75.28 | 8.81 | 0.50:0.50 | 2.0 | 76.45 | 7.99 | 0.50:0.50 | 5.0 | 72.38 | 8.58 | 0.20:0.80 |
| adni_128.0-1.5 | 5.0 | 74.4 | 4.39 | 0.40:0.60 | 4.0 | 74.22 | 3.67 | 0.50:0.50 | 9.0 | 78.22 | 7.07 | 0.22:0.78 |
| adni_16.0-3.0 | 3.0 | 71.9 | 2.45 | 0.67:0.33 | 3.0 | 80.6 | 9.22 | 0.67:0.33 | 8.0 | 71.69 | 9.21 | 0.62:0.38 |
| adni_2.0-1.5 | 3.0 | 75.53 | 4.37 | 0.67:0.33 | 3.0 | 81.97 | 9.54 | 0.67:0.33 | 5.0 | 75.98 | 5.01 | 0.60:0.40 |
| adni_23.0-1.5 | 5.0 | 73.68 | 3.35 | 0.60:0.40 | 3.0 | 74.17 | 3.27 | 0.67:0.33 | 7.0 | 74.59 | 4.56 | 0.71:0.29 |
| adni_29.0-3.0 | 4.0 | 72.08 | 9.12 | 0.25:0.75 | 2.0 | 72.45 | 9.83 | 0:1 | 1.0 | 78.6 | | 0:1 |
| adni_3.0-3.0 | 5.0 | 68.44 | 5.32 | 0.80:0.20 | 4.0 | 71.6 | 5.86 | 0.75:0.25 | 5.0 | 69.52 | 8.12 | 0.60:0.40 |
| adni_33.0-1.5 | 4.0 | 77.02 | 5.56 | 0.50:0.50 | 4.0 | 82.0 | 3.99 | 0.50:0.50 | 8.0 | 73.6 | 9.76 | 0.50:0.50 |
| adni_41.0-3.0 | 3.0 | 74.17 | 5.77 | 0.33:0.67 | 3.0 | 79.43 | 4.52 | 0.33:0.67 | | | | |
| adni_73.0-3.0 | 4.0 | 66.5 | 6.47 | 0.50:0.50 | 4.0 | 70.55 | 6.46 | 0.50:0.50 | 3.0 | 64.73 | 6.01 | 0.67:0.33 |
| adni_94.0-1.5 | 4.0 | 77.18 | 5.77 | 0.50:0.50 | 2.0 | 75.15 | 2.47 | 0.50:0.50 | 5.0 | 67.62 | 7.1 | 0.80:0.20 |
| aibl_Melbourne | 137.0 | 72.93 | 6.11 | 0.60:0.40 | 136.0 | 73.03 | 6.06 | 0.60:0.40 | 6.0 | 70.33 | 4.68 | 0.17:0.83 |
| aibl_Perth | 43.0 | 70.74 | 6.01 | 0.60:0.40 | 42.0 | 71.07 | 5.77 | 0.62:0.38 | 2.0 | 67.5 | 2.12 | 0.50:0.50 |
| aomic_BEST | 398.0 | 22.81 | 1.71 | 0.54:0.46 | 397.0 | 22.82 | 1.7 | 0.53:0.47 | | | | |
| beijing_BNU | 90.0 | 21.19 | 1.99 | 0.60:0.40 | 89.0 | 21.24 | 1.9 | 0.60:0.40 | | | | |
| camcan_Cambridge | 323.0 | 54.68 | 18.56 | 0.51:0.49 | 322.0 | 54.79 | 18.49 | 0.51:0.49 | | | | |
| corr_BMB | 25.0 | 30.11 | 6.23 | 0.52:0.48 | 25.0 | 31.54 | 7.93 | 0.52:0.48 | | | | |
| corr_BNU_1 | 25.0 | 22.88 | 2.26 | 0.48:0.52 | 24.0 | 23.17 | 2.51 | 0.50:0.50 | | | | |
| corr_BNU_2 | 15.0 | 21.38 | 0.87 | 0.40:0.60 | 13.0 | 21.42 | 0.73 | 0.38:0.62 | | | | |
| corr_BNU_3 | 10.0 | 22.5 | 3.27 | 0.60:0.40 | 8.0 | 22.0 | 1.69 | 0.62:0.38 | | | | |
| corr_IACAS | 14.0 | 26.29 | 6.07 | 0.50:0.50 | 13.0 | 26.46 | 3.95 | 0.54:0.46 | | | | |
| corr_IBATRT | 18.0 | 26.28 | 7.04 | 0.50:0.50 | 18.0 | 28.39 | 8.67 | 0.50:0.50 | | | | |
| corr_IPCAS_1 | 16.0 | 20.88 | 1.96 | 0.69:0.31 | 14.0 | 20.93 | 1.59 | 0.71:0.29 | | | | |
| corr_IPCAS_2 | 17.0 | 13.24 | 1.09 | 0.65:0.35 | 17.0 | 13.35 | 0.93 | 0.65:0.35 | | | | |
| corr_IPCAS_3 | 18.0 | 20.89 | 1.75 | 0.67:0.33 | 17.0 | 21.29 | 1.76 | 0.65:0.35 | | | | |
| corr_IPCAS_4 | 10.0 | 23.3 | 2.0 | 0.50:0.50 | 9.0 | 23.0 | 1.22 | 0.56:0.44 | | | | |
| corr_IPCAS_5 | 11.0 | 18.27 | 0.47 | 0:1 | 11.0 | 18.36 | 0.5 | 0:1 | | | | |
| corr_IPCAS_7 | 35.0 | 11.63 | 3.14 | 0.57:0.43 | 33.0 | 11.7 | 3.12 | 0.58:0.42 | | | | |
| corr_IPCAS_8 | 7.0 | 57.29 | 3.95 | 0.57:0.43 | 6.0 | 58.0 | 3.9 | 0.50:0.50 | | | | |
| corr_LMU_1 | 14.0 | 24.0 | 1.88 | 0.50:0.50 | 13.0 | 24.62 | 1.85 | 0.54:0.46 | | | | |
| corr_LMU_2 | 14.0 | 41.0 | 22.26 | 0.50:0.50 | 13.0 | 42.46 | 22.03 | 0.54:0.46 | | | | |
| corr_LMU_3 | 6.0 | 65.17 | 6.52 | 0.33:0.67 | 6.0 | 68.33 | 8.87 | 0.33:0.67 | | | | |
| corr_MCGILL | 40.0 | 65.05 | 6.17 | 0.72:0.28 | 40.0 | 65.68 | 6.41 | 0.72:0.28 | | | | |
| corr_NKI | 5.0 | 33.8 | 11.45 | 1:0 | 5.0 | 36.8 | 13.33 | 1:0 | | | | |
| corr_NYU_1 | 13.0 | 30.31 | 9.81 | 0.62:0.38 | 11.0 | 28.82 | 7.77 | 0.64:0.36 | | | | |
| corr_NYU_2 | 85.0 | 19.28 | 10.99 | 0.36:0.64 | 85.0 | 19.93 | 11.65 | 0.36:0.64 | | | | |
| corr_SWU_1 | 10.0 | 21.4 | 1.78 | 0.70:0.30 | 10.0 | 21.7 | 1.83 | 0.70:0.30 | | | | |
| corr_SWU_2 | 14.0 | 20.93 | 1.77 | 0.64:0.36 | 13.0 | 21.0 | 1.58 | 0.69:0.31 | | | | |
| corr_SWU_3 | 12.0 | 20.5 | 1.93 | 0.67:0.33 | 11.0 | 20.36 | 1.36 | 0.64:0.36 | | | | |
| corr_SWU_4 | 115.0 | 20.02 | 1.25 | 0.50:0.50 | 115.0 | 20.1 | 1.32 | 0.50:0.50 | | | | |
| corr_UPSM | 49.0 | 15.09 | 2.83 | 0.49:0.51 | 49.0 | 15.29 | 2.81 | 0.49:0.51 | | | | |

| | | | | | | | | | | | | |
|---|---|---|---|---|---|---|---|---|---|---|---|---|
| corr_XHCUMS | 12.0 | 51.67 | 6.97 | 0.42:0.58 | 10.0 | 51.2 | 5.57 | 0.40:0.60 | | | | |
| dlbs_UTSMC | 156.0 | 54.23 | 20.12 | 0.63:0.37 | 155.0 | 54.48 | 20.01 | 0.63:0.37 | | | | |
| ds000119 | 36.0 | 16.01 | 4.9 | 0.61:0.39 | 35.0 | 16.24 | 4.75 | 0.60:0.40 | | | | |
| ds000202 | 37.0 | 22.22 | 2.75 | 1:0 | 37.0 | 22.41 | 2.9 | 1:0 | | | | |
| ds000222 | 40.0 | 44.12 | 20.28 | 0.53:0.47 | 39.0 | 44.74 | 20.22 | 0.51:0.49 | | | | |
| fcon1000_AnnArbor | 8.0 | 20.5 | 8.33 | 0.25:0.75 | 6.0 | 20.17 | 6.94 | 0.17:0.83 | | | | |
| fcon1000_Atlanta | 12.0 | 30.75 | 10.86 | 0.50:0.50 | 11.0 | 30.09 | 8.62 | 0.55:0.45 | | | | |
| fcon1000_Baltimore | 11.0 | 29.0 | 6.16 | 0.64:0.36 | 10.0 | 30.0 | 5.19 | 0.70:0.30 | | | | |
| fcon1000_Beijing | 99.0 | 21.14 | 1.86 | 0.62:0.38 | 98.0 | 21.18 | 1.81 | 0.62:0.38 | | | | |
| fcon1000_Berlin | 12.0 | 30.08 | 5.95 | 0.50:0.50 | 10.0 | 29.3 | 4.67 | 0.50:0.50 | | | | |
| fcon1000_Cambridge | 99.0 | 21.08 | 2.39 | 0.63:0.37 | 97.0 | 21.0 | 2.25 | 0.63:0.37 | | | | |
| fcon1000_Dallas | 12.0 | 41.75 | 20.54 | 0.50:0.50 | 11.0 | 41.18 | 19.73 | 0.55:0.45 | | | | |
| fcon1000_Leiden_2200 | 10.0 | 21.7 | 2.87 | 0.40:0.60 | 9.0 | 21.67 | 2.35 | 0.44:0.56 | | | | |
| fcon1000_Milwaukee_b | 23.0 | 53.48 | 6.07 | 0.70:0.30 | 21.0 | 53.62 | 5.86 | 0.71:0.29 | | | | |
| fcon1000_Munchen | 5.0 | 69.0 | 4.06 | 0.60:0.40 | 4.0 | 69.5 | 4.65 | 0.50:0.50 | | | | |
| fcon1000_NewHaven_a | 8.0 | 29.12 | 9.99 | 0.50:0.50 | 8.0 | 34.62 | 10.81 | 0.50:0.50 | | | | |
| fcon1000_NewHaven_b | 8.0 | 25.75 | 5.55 | 0.50:0.50 | 7.0 | 28.43 | 7.59 | 0.43:0.57 | | | | |
| fcon1000_NewYork_a | 28.0 | 29.71 | 11.45 | 0.39:0.61 | 26.0 | 29.42 | 10.5 | 0.38:0.62 | | | | |
| hbn_CBIC | 319.0 | 10.75 | 3.58 | 0.66:0.34 | 317.0 | 10.73 | 3.53 | 0.66:0.34 | | | | |
| hbn_Rutgers | 408.0 | 10.5 | 3.56 | 0.64:0.36 | 407.0 | 10.51 | 3.56 | 0.64:0.36 | | | | |
| hbn_StatenIsland | 160.0 | 12.13 | 3.87 | 0.56:0.44 | 159.0 | 12.15 | 3.85 | 0.56:0.44 | | | | |
| hcp_WU | 547.0 | 28.81 | 3.71 | 0.55:0.45 | 546.0 | 28.82 | 3.7 | 0.55:0.45 | | | | |
| mpi_Leipzig | 37.0 | 26.15 | 6.94 | 0.11:0.89 | 36.0 | 27.08 | 9.81 | 0.08:0.92 | | | | |
| nki | 448.0 | 37.46 | 21.59 | 0.62:0.38 | 448.0 | 37.64 | 21.66 | 0.62:0.38 | | | | |
| oasis3_STL_2 | 289.0 | 65.46 | 9.12 | 0.61:0.39 | 287.0 | 65.47 | 8.97 | 0.61:0.39 | | | | |
| ping_PC | 35.0 | 14.27 | 3.76 | 0.54:0.46 | 34.0 | 14.57 | 3.61 | 0.56:0.44 | | | | |
| ping_PD | 47.0 | 12.72 | 6.18 | 0.49:0.51 | 46.0 | 12.96 | 6.08 | 0.48:0.52 | | | | |
| ping_PG | 53.0 | 11.62 | 3.98 | 0.51:0.49 | 51.0 | 11.51 | 3.7 | 0.51:0.49 | | | | |
| ping_PJ | 50.0 | 14.76 | 4.38 | 0.48:0.52 | 49.0 | 14.99 | 4.16 | 0.49:0.51 | | | | |
| ping_PM | 24.0 | 13.66 | 4.51 | 0.46:0.54 | 22.0 | 13.72 | 4.19 | 0.45:0.55 | | | | |
| ping_PU_1 | 47.0 | 14.59 | 4.38 | 0.45:0.55 | 46.0 | 14.82 | 4.12 | 0.46:0.54 | | | | |
| ping_PU_2 | 52.0 | 8.97 | 3.71 | 0.48:0.52 | 51.0 | 9.05 | 3.62 | 0.47:0.53 | | | | |
| ping_PY | 39.0 | 13.16 | 5.32 | 0.54:0.46 | 37.0 | 13.24 | 5.13 | 0.54:0.46 | | | | |
| pnc | 741.0 | 14.96 | 3.68 | 0.53:0.47 | 739.0 | 14.95 | 3.66 | 0.53:0.47 | | | | |
| sald_SUC | 244.0 | 45.03 | 17.34 | 0.63:0.37 | 244.0 | 45.3 | 17.4 | 0.63:0.37 | | | | |
| slim_SUC | 282.0 | 20.08 | 1.26 | 0.56:0.44 | 282.0 | 20.11 | 1.29 | 0.56:0.44 | | | | |
| top15_Oslo | 43.0 | 34.05 | 9.09 | 0.37:0.63 | 42.0 | 34.55 | 9.14 | 0.38:0.62 | 157.0 | 32.08 | 10.2 | 0.47:0.53 |
| top3_Oslo | 154.0 | 30.75 | 8.19 | 0.43:0.57 | 152.0 | 30.78 | 7.99 | 0.43:0.57 | 157.0 | 29.24 | 10.4 | 0.43:0.57 |
| top750_Oslo | 391.0 | 44.75 | 18.11 | 0.55:0.45 | 390.0 | 44.82 | 18.04 | 0.56:0.44 | 484.0 | 46.74 | 21.05 | 0.46:0.54 |
| ukb_Cheadle | 11897.0 | 63.6 | 7.55 | 0.53:0.47 | 11896.0 | 63.6 | 7.54 | 0.53:0.47 | | | | |
| ukb_Newcastle | 4998.0 | 65.11 | 7.45 | 0.54:0.46 | 4997.0 | 65.11 | 7.44 | 0.54:0.46 | | | | |
| ukb_Reading | 2744.0 | 65.87 | 7.57 | 0.54:0.46 | 2742.0 | 65.87 | 7.56 | 0.54:0.46 | | | | |

**Supplementary Table 3. Group based nonparametric tests of cerebellar lobules**

| ROI | Diagnosis | U-value | P-value | RBC | CLES | Median Control | Median Clinical |
|---|---|---|---|---|---|---|---|
| Corpus.Medullare | asd | 12356969.0 | 0.3683246482309873 | -0.01759981882939088 | 0.5087999094146954 | -0.003 | -0.03 |
| Left.Crus.I | asd | 12347572.0 | 0.3897555717908264 | -0.016825973277335038 | 0.5084129866386675 | 0.013 | -0.013 |
| Left.Crus.II | asd | 12207608.0 | 0.7864654473922184 | -0.005299899120910778 | 0.5026499495604554 | 0.008 | 0.029 |
| Left.I.III | asd | 12362835.5 | 0.3553248670890835 | -0.018082926728841153 | 0.5090414633644206 | 0.008 | -0.007 |
| Left.IV | asd | 12434119.0 | 0.22081368349177832 | -0.023953142692442198 | 0.5119765713462211 | -0.007 | -0.078 |
| Left.IX | asd | 12233453.0 | 0.7041726488046728 | -0.007428242027463838 | 0.5037141210137319 | 0.034 | 0.027 |
| Left.V | asd | 13104574.0 | 5.1978659974309194e-05 | -0.07916529759331326 | 0.5395826487966566 | -0.014 | -0.204 |
| Left.VI | asd | 12221426.0 | 0.7421042555439723 | -0.006437815247153811 | 0.5032189076235769 | -0.001 | 0.003 |
| Left.VIIB | asd | 12333418.0 | 0.4234318316092971 | -0.01566038745805276 | 0.5078301937290264 | 0.018 | -0.003 |
| Left.VIIIA | asd | 12116653.0 | 0.9108598963576383 | 0.0021902703147839153 | 0.49890486484260804 | -0.001 | 0.006 |
| Left.VIIIB | asd | 12455560.0 | 0.18863763948833345 | -0.025718814979515336 | 0.5128594074897577 | 0.034 | 0.015 |
| Left.X | asd | 12306221.0 | 0.4927131802506728 | -0.013420706977127272 | 0.5067103534885636 | 0.021 | -0.023 |
| Right.Crus.I | asd | 12560139.0 | 0.07928845242720665 | -0.0343309245877339 | 0.517165462293867 | 0.028 | -0.021 |
| Right.Crus.II | asd | 12603662.0 | 0.052619174495313586 | -0.03791505568937481 | 0.5189575278446874 | 0.018 | -0.016 |
| Right.I.III | asd | 12225973.0 | 0.7276840091502537 | -0.006812261956230747 | 0.5034061309781154 | -0.02 | 0.018 |
| Right.IX | asd | 11863141.0 | 0.23836841803491726 | 0.023067053712968133 | 0.48846647314351593 | 0.04 | 0.085 |
| Right.V | asd | 13344561.0 | 4.2649201071891454e-07 | -0.09892829349638688 | 0.5494641467481934 | -0.022 | -0.182 |
| Right.VI | asd | 11842022.0 | 0.20480795022920828 | 0.024806209210878505 | 0.48759689539456075 | 0.013 | 0.069 |
| Right.VIIB | asd | 11904122.0 | 0.3141408034866936 | 0.019692257015214265 | 0.49015387149239287 | 0.007 | 0.094 |
| Right.VIIIA | asd | 11509239.0 | 0.0076129076460311465 | 0.05221098140942504 | 0.4738945092952875 | 0.018 | 0.135 |
| Right.VIIIB | asd | 12374389.0 | 0.3305817008277181 | -0.019034360653037652 | 0.5095171803265188 | 0.014 | -0.012 |
| Right.X | asd | 12348684.0 | 0.3871806760995682 | -0.016917546785251147 | 0.5084587733926256 | 0.007 | -0.018 |
| Rigt.IV | asd | 12460125.5 | 0.1822570184394603 | -0.026094785168715218 | 0.5130473925843576 | -0.017 | -0.055 |
| Vermis.IX | asd | 12533235.0 | 0.10067680910306306 | -0.03211537273794085 | 0.5160576863689704 | 0.012 | 0.004 |
| Vermis.VI | asd | 12318868.0 | 0.45976398562256404 | -0.014462190929116936 | 0.5072310954645585 | 0.003 | -0.024 |
| Vermis.VII | asd | 12712806.0 | 0.016509251648698493 | -0.046903094311654625 | 0.5234515471558273 | 0.007 | -0.076 |
| Vermis.VIII | asd | 12028138.0 | 0.6279994192837113 | 0.009479505074835859 | 0.49526024746258207 | 0.016 | 0.031 |
| Vermis.X | asd | 13048666.5 | 0.00013828659769703737 | -0.07456129948736945 | 0.5372806497436847 | 0.015 | -0.12 |
| Corpus.Medullare | ad | 2044196.0 | 0.43120694046863794 | -0.03771298615923113 | 0.5188564930796156 | -0.003 | -0.009 |
| Left.Crus.I | ad | 2316433.0 | 0.00024108120195350416 | -0.17591102108984957 | 0.5879555105449248 | 0.013 | -0.267 |
| Left.Crus.II | ad | 1974161.0 | 0.9640370253699901 | -0.0021605102784143515 | 0.5010802551392072 | 0.008 | 0.008 |
| Left.I.III | ad | 1888522.0 | 0.3885391428464511 | 0.041313159771664165 | 0.4793434201141679 | 0.008 | 0.071 |
| Left.IV | ad | 2312759.0 | 0.0002805516882258359 | -0.17404595653089872 | 0.5870229782654494 | -0.007 | -0.263 |
| Left.IX | ad | 1889482.0 | 0.39415958961112185 | 0.04082582662615708 | 0.47958708668692146 | 0.034 | 0.142 |
| Left.V | ad | 2120157.0 | 0.11139472645735497 | -0.07627372893616702 | 0.5381368644680835 | -0.014 | -0.114 |
| Left.VI | ad | 2052280.0 | 0.3827829091505178 | -0.04181673735535463 | 0.5209083686776773 | -0.001 | -0.041 |
| Left.VIIB | ad | 2216034.0 | 0.009112665363159954 | -0.12494460392760054 | 0.5624723019638003 | 0.018 | -0.24 |
| Left.VIIIA | ad | 1663783.0 | 0.001180955445552225 | 0.15539937205093646 | 0.42230031397453177 | -0.001 | 0.238 |
| Left.VIIIB | ad | 1943581.0 | 0.7803180207443725 | 0.013363080960757001 | 0.4933184595196215 | 0.034 | 0.046 |
| Left.X | ad | 1549488.0 | 8.41175988537094e-06 | 0.2134199364943995 | 0.39329003175280025 | 0.021 | 0.391 |
| Right.Crus.I | ad | 2355598.0 | 4.378980590871989e-05 | -0.1957926905104561 | 0.5978963452552281 | 0.028 | -0.447 |
| Right.Crus.II | ad | 1905711.0 | 0.496411042048718 | 0.03258735827362236 | 0.4837063208631888 | 0.018 | 0.078 |
| Right.I.III | ad | 2133507.0 | 0.08302419440673349 | -0.08305070549087401 | 0.541525352745437 | -0.02 | -0.177 |
| Right.IX | ad | 1930382.0 | 0.6753976744921834 | 0.020063404072785285 | 0.48996829796360736 | 0.04 | 0.059 |

| | | | | | | | |
|---|---|---|---|---|---|---|---|
| Right.V | ad | 2458081.0 | 2.3116577758022247e-07 | -0.24781702670940975 | 0.6239085133547049 | -0.022 | -0.412 |
| Right.VI | ad | 2048212.0 | 0.40672051913678353 | -0.03975166315126866 | 0.5198758315756343 | 0.013 | -0.008 |
| Right.VIIB | ad | 2072597.0 | 0.27657555214169904 | -0.05213043268584028 | 0.5260652163429201 | 0.007 | -0.127 |
| Right.VIIIA | ad | 1618884.0 | 0.00019988845736611488 | 0.17819184173856095 | 0.4109040791307195 | 0.018 | 0.281 |
| Right.VIIIB | ad | 1999401.0 | 0.7546508191771495 | -0.014973310895703085 | 0.5074866554478515 | 0.014 | -0.023 |
| Right.X | ad | 1527428.0 | 2.7567441897444275e-06 | 0.22461844606719616 | 0.3876907769664019 | 0.007 | 0.438 |
| Rigt.IV | ad | 2121593.0 | 0.10801703537841888 | -0.07700269809965454 | 0.5385013490498273 | -0.017 | -0.088 |
| Vermis.IX | ad | 2061578.0 | 0.3314013959409675 | -0.046536761925067394 | 0.5232683809625337 | 0.012 | -0.067 |
| Vermis.VI | ad | 2148768.0 | 0.05807959116196852 | -0.09079777958835589 | 0.5453988897941779 | 0.003 | -0.201 |
| Vermis.VII | ad | 1995436.0 | 0.786773637911772 | -0.01296052347702048 | 0.5064802617385102 | 0.007 | -0.031 |
| Vermis.VIII | ad | 1871118.0 | 0.2952519137075724 | 0.05014810358875177 | 0.4749259482056241 | 0.016 | 0.052 |
| Vermis.X | ad | 1709296.0 | 0.005758626809369336 | 0.13229521220566476 | 0.4338523938971676 | 0.015 | 0.229 |
| Corpus.Medullare | mci | 1777517.0 | 0.12749533863957652 | -0.0798452084795136 | 0.5399226042397568 | -0.003 | -0.145 |
| Left.Crus.I | mci | 1798934.0 | 0.07632804795644665 | -0.09285607972856802 | 0.546428039864284 | 0.013 | -0.121 |
| Left.Crus.II | mci | 1699668.0 | 0.5343811943816714 | -0.0325517819553669 | 0.5162758909776834 | 0.008 | -0.062 |
| Left.I.III | mci | 1503831.0 | 0.09903638125938848 | 0.08641959558589019 | 0.4567902022070549 | 0.008 | 0.187 |
| Left.IV | mci | 1739586.0 | 0.2782702793184071 | -0.05680204849688808 | 0.528401024248444 | -0.007 | -0.18 |
| Left.IX | mci | 1532973.0 | 0.1896489019929012 | 0.06871577105678017 | 0.4656421144716099 | 0.034 | 0.322 |
| Left.V | mci | 1739107.0 | 0.2807398524738238 | -0.05651105501842246 | 0.5282555275092112 | -0.014 | -0.061 |
| Left.VI | mci | 1742279.0 | 0.2646616189974429 | -0.05843805149794812 | 0.5292190257489741 | -0.001 | -0.065 |
| Left.VIIB | mci | 1883722.0 | 0.0058587173526226805 | -0.14436496292718792 | 0.572182481463594 | 0.018 | -0.156 |
| Left.VIIIA | mci | 1501034.0 | 0.0925733484840645 | 0.08811877879939367 | 0.45594061060030316 | -0.001 | -0.013 |
| Left.VIIIB | mci | 1685620.0 | 0.6466405262840675 | -0.02401759325915731 | 0.5120087966295787 | 0.034 | 0.178 |
| Left.X | mci | 1595231.0 | 0.5554004640171246 | 0.030893908880768595 | 0.4845530455596157 | 0.021 | 0.06 |
| Right.Crus.I | mci | 1883580.0 | 0.0058882743902148385 | -0.14427869763712087 | 0.5721393488185604 | 0.028 | -0.259 |
| Right.Crus.II | mci | 1669110.0 | 0.7894787678684858 | -0.013987734533757346 | 0.5069938672668787 | 0.018 | 0.024 |
| Right.I.III | mci | 1603240.0 | 0.6193185989758021 | 0.026028425020579116 | 0.48698578748971044 | -0.02 | 0.088 |
| Right.IX | mci | 1615283.0 | 0.7209659893883023 | 0.018712277920034492 | 0.49064386103998275 | 0.04 | 0.036 |
| Right.V | mci | 2118489.0 | 4.304104576467849e-08 | -0.28698639499175327 | 0.6434931974958766 | -0.022 | -0.528 |
| Right.VI | mci | 1599251.0 | 0.5870802635485538 | 0.028451750669011622 | 0.4857741246654942 | 0.013 | 0.006 |
| Right.VIIB | mci | 1578297.0 | 0.4318379794716708 | 0.041181348472284274 | 0.47940932576385786 | 0.007 | 0.067 |
| Right.VIIIA | mci | 1384644.0 | 0.0024325162527701148 | 0.1588259415522285 | 0.42058702922388574 | 0.018 | 0.287 |
| Right.VIIIB | mci | 1812620.0 | 0.05347057067810026 | -0.10117035268531094 | 0.5505851763426555 | 0.014 | -0.2 |
| Right.X | mci | 1588690.0 | 0.5057082326054114 | 0.034867579742236865 | 0.48256621012888157 | 0.007 | 0.038 |
| Rigt.IV | mci | 1713984.0 | 0.43108432182962797 | -0.04124878119902675 | 0.5206243905995134 | -0.017 | -0.186 |
| Vermis.IX | mci | 1675538.5 | 0.7327037735604642 | -0.01789306141541891 | 0.5089465307077095 | 0.012 | -0.08 |
| Vermis.VI | mci | 1629874.0 | 0.8508972413686849 | 0.009848215614625011 | 0.4950758921926875 | 0.003 | -0.021 |
| Vermis.VII | mci | 1629654.0 | 0.8488979034804802 | 0.009981866064024625 | 0.4950090669679877 | 0.007 | 0.026 |
| Vermis.VIII | mci | 1513675.0 | 0.12468702070780276 | 0.08043934547729914 | 0.45978032726135043 | 0.016 | 0.245 |
| Vermis.X | mci | 1672106.0 | 0.7628590076400781 | -0.015807810653763354 | 0.5079039053268817 | 0.015 | -0.007 |
| Corpus.Medullare | bd | 3549472.0 | 0.1492276569330777 | 0.05028880197515806 | 0.47485559901242097 | -0.003 | 0.099 |
| Left.Crus.I | bd | 4025534.0 | 0.0270450738670583 | -0.07708828744943874 | 0.5385441437247194 | 0.013 | -0.09 |
| Left.Crus.II | bd | 3984189.0 | 0.05827824980921621 | -0.06602585070325873 | 0.5330129253516294 | 0.008 | -0.103 |
| Left.I.III | bd | 3731960.5 | 0.9665706766768903 | 0.0014614349862772658 | 0.49926928250686137 | 0.008 | -0.008 |
| Left.IV | bd | 3899634.0 | 0.21322274129802443 | -0.0434019702080779 | 0.521700985104039 | -0.007 | -0.102 |
| Left.IX | bd | 4060855.0 | 0.013068098328197296 | -0.08653891819830384 | 0.5432694590991519 | 0.034 | -0.175 |
| Left.V | bd | 3691493.0 | 0.7245055683183441 | 0.012289084255258764 | 0.4938554578723706 | -0.014 | 0.003 |

| | | | | | | | |
|---|---|---|---|---|---|---|---|
| Left.VI | bd | 3839390.0 | 0.43394407016473924 | -0.027282839978621576 | 0.5136414199893108 | -0.001 | -0.05 |
| Left.VIIB | bd | 3849741.0 | 0.3887478064481775 | -0.030052395735296278 | 0.5150261978676481 | 0.018 | -0.115 |
| Left.VIIIA | bd | 3732438.0 | 0.9694919158450784 | 0.0013336731397105561 | 0.4993331634301447 | -0.001 | -0.025 |
| Left.VIIIB | bd | 3764364.0 | 0.836215926157071 | -0.007208577569167041 | 0.5036042887845835 | 0.034 | 0.011 |
| Left.X | bd | 3696963.0 | 0.7562045435572107 | 0.010825508756368896 | 0.49458724562181555 | 0.021 | 0.007 |
| Right.Crus.I | bd | 3853234.0 | 0.3741672605173968 | -0.030986997054788468 | 0.5154934985273942 | 0.028 | -0.079 |
| Right.Crus.II | bd | 3919142.0 | 0.16318166419071467 | -0.04862161021399114 | 0.5243108051069956 | 0.018 | -0.059 |
| Right.I.III | bd | 3495193.0 | 0.06305781207661428 | 0.06481191248781748 | 0.46759404375609126 | -0.02 | 0.076 |
| Right.IX | bd | 3927565.0 | 0.14454155660499557 | -0.05087530243102023 | 0.5254376512155101 | 0.04 | -0.031 |
| Right.V | bd | 3784231.5 | 0.7194509818888588 | -0.012524406860610515 | 0.5062622034303053 | -0.022 | 0.006 |
| Right.VI | bd | 3937334.0 | 0.12501745502132922 | -0.053489135895125495 | 0.5267445679475627 | 0.013 | -0.087 |
| Right.VIIB | bd | 3959129.0 | 0.08888721938703234 | -0.05932069494417602 | 0.529660347472088 | 0.007 | -0.117 |
| Right.VIIIA | bd | 3880580.0 | 0.27196940226047683 | -0.03830380429293179 | 0.5191519021464659 | 0.018 | -0.018 |
| Right.VIIIB | bd | 3929464.0 | 0.1405737347646773 | -0.05138340661244478 | 0.5256917033062224 | 0.014 | -0.046 |
| Right.X | bd | 3625916.5 | 0.3921883082775389 | 0.029834999923075367 | 0.4850825000384623 | 0.007 | 0.105 |
| Rigt.IV | bd | 3753528.5 | 0.9016413560171705 | -0.004309387017389588 | 0.5021546935086948 | -0.017 | -0.002 |
| Vermis.IX | bd | 3894091.0 | 0.22927931438283733 | -0.0419188625315976 | 0.5209594312657988 | 0.012 | -0.02 |
| Vermis.VI | bd | 4009564.0 | 0.036769137155047135 | -0.07281528914646396 | 0.536407644573232 | 0.003 | -0.109 |
| Vermis.VII | bd | 3751424.5 | 0.9144377108223032 | -0.0037464322002662698 | 0.5018732161001331 | 0.007 | -0.033 |
| Vermis.VIII | bd | 3806345.0 | 0.5968857966675553 | -0.01844118506805148 | 0.5092205925340257 | 0.016 | -0.092 |

**Supplementary Table 4**. Anatomical atlas based case-control analyses of extreme deviations

a) positive deviation

| ROI | Diagnosis | U-value | P-value | RBC | CLES | Median Clinical | Median Control |
|---|---|---|---|---|---|---|---|
| Left. Crus.I | asd | 10321891.0 | 1.7630727559821753e-14 | 0.149989417989418 | 0.425005291005291 | 0.011 | 0.007 |
| Left.Crus.II | asd | 8708620.5 | 2.0676489262978824e-47 | 0.28284269038354637 | 0.3585786548082268 | 0.012 | 0.004 |
| Left.VIIB | asd | 7151688.5 | 3.534346746725028e-98 | 0.41105647170238613 | 0.29447176414880694 | 0.018 | 0.004 |
| Left.VI | asd | 8917097.5 | 5.208304281043644e-42 | 0.2656745517056801 | 0.36716272414715995 | 0.01 | 0.005 |
| Left.VIIIA | asd | 7120217.5 | 1.8788972528552227e-99 | 0.4136481172667943 | 0.29317594136660285 | 0.019 | 0.003 |
| Left..VI | asd | 9566153.5 | 1.9289655444831937e-27 | 0.21222461038025242 | 0.3938876948098738 | 0.01 | 0.005 |
| Left.X | asd | 10075887.5 | 4.359607681859814e-19 | 0.17024787433347743 | 0.4148760628332613 | 0.011 | 0.004 |
| Left.IV | asd | 9603653.0 | 1.0801338394815358e-26 | 0.2091365161715356 | 0.3954317419142322 | 0.01 | 0.006 |
| Left.VIIIA | asd | 8062446.5 | 2.0743295355633886e-66 | 0.33605529821094027 | 0.33197235089452987 | 0.013 | 0.003 |
| Left.IX | asd | 9210968.5 | 4.2818947044810174e-35 | 0.24147419348197563 | 0.3792629032590122 | 0.009 | 0.004 |
| Left.I.III | asd | 10109958.0 | 6.059231126587038e-19 | 0.16744215922426042 | 0.4162789203878698 | 0.005 | 0.001 |
| Vermis.VIII | asd | 9684883.0 | 2.5489090557996483e-25 | 0.20244720317872067 | 0.39877639841063967 | 0.009 | 0.003 |
| Vermis.IX | asd | 10605202.5 | 2.340455371844139e-11 | 0.12665863751466866 | 0.43667068124266567 | 0.002 | 0.001 |
| Vermis.VII | asd | 11087320.0 | 3.725021536030678e-06 | 0.08695612788998008 | 0.45652193605500996 | 0.003 | 0.001 |
| Vermis.VI | asd | 11741958.5 | 0.08811899682271558 | 0.0330464661437424 | 0.4834767669281288 | 0.003 | 0.003 |
| Vermis.X | asd | 10994923.0 | 5.302599353754356e-07 | 0.09456504642496866 | 0.45271747678751567 | 0.011 | 0.004 |
| Right.V | asd | 9813615.5 | 1.0001724392683929e-22 | 0.19184604615733025 | 0.4040769769213349 | 0.009 | 0.005 |
| Right.IV | asd | 9760710.0 | 1.1061064974666677e-23 | 0.19620282873201156 | 0.4018985856339942 | 0.012 | 0.006 |
| Right.IX | asd | 9267514.5 | 7.381088077857814e-34 | 0.23681761472422946 | 0.38159119263788527 | 0.009 | 0.003 |
| Right.I.III | asd | 10251910.5 | 3.004285011214546e-16 | 0.1557523315422148 | 0.4221238342288926 | 0.005 | 0.002 |
| Right.Crus.II | asd | 8631230.5 | 1.540042996858127e-49 | 0.28921577831305456 | 0.3553921108434727 | 0.011 | 0.003 |

| | | | | | | | |
|---|---|---|---|---|---|---|---|
| Right.VIIB | asd | 7333145.5 | 1.8194871567126124e-91 | 0.3961134375064336 | 0.3019432812467832 | 0.017 | 0.003 |
| Right.VI | asd | 9436655.5 | 4.4720645303874086e-30 | 0.22288880653861198 | 0.388555596730694 | 0.009 | 0.004 |
| Right.Crus.I | asd | 10244024.0 | 1.2998549251336448e-15 | 0.15640178700100882 | 0.4217991064994956 | 0.01 | 0.006 |
| Right.Crus.II | asd | 7303350.0 | 1.252276413559156e-92 | 0.3985671051818912 | 0.3007164474090544 | 0.018 | 0.003 |
| Right.VIIIB | asd | 8052182.0 | 1.0863030365936386e-66 | 0.3369005826282091 | 0.33154970868589545 | 0.014 | 0.003 |
| Right.X | asd | 10099907.0 | 2.7106855496728963e-18 | 0.16826986185741044 | 0.4158650690712948 | 0.014 | 0.005 |
| Left. Crus.I | mci | 1213415.5 | 5.242892193380137e-07 | 0.2628476050750721 | 0.36857619746246395 | 0.016 | 0.007 |
| Left.Crus.II | mci | 1292396.0 | 4.077868078820629e-05 | 0.2148667899895813 | 0.39256660500520935 | 0.009 | 0.004 |
| Left.VIIB | mci | 1192241.5 | 1.3825359677099181e-07 | 0.27571085332774437 | 0.3621445733361278 | 0.011 | 0.004 |
| Left.VI | mci | 1319403.5 | 0.0001516494732453922 | 0.19845967857066915 | 0.4007701607146654 | 0.009 | 0.005 |
| Left.VIIIA | mci | 1380366.5 | 0.002035380895743104 | 0.16142453153998726 | 0.41928773423000637 | 0.006 | 0.003 |
| Left..VI | mci | 1494896.0 | 0.07942676596369258 | 0.09184762633764354 | 0.45407618683117823 | 0.005 | 0.005 |
| Left.X | mci | 1371330.0 | 0.0010731333411950284 | 0.16691422374907738 | 0.4165428881254613 | 0.009 | 0.004 |
| Left.IV | mci | 1514985.0 | 0.1282935936299526 | 0.07964351780132861 | 0.4601782410993357 | 0.007 | 0.006 |
| Left.VIIIA | mci | 1505202.0 | 0.10154961547729263 | 0.08558671028531328 | 0.45720664485734336 | 0.004 | 0.003 |
| Left.IX | mci | 1428872.5 | 0.01165320658420932 | 0.13195703745553844 | 0.4340214812722308 | 0.006 | 0.004 |
| Left.I.III | mci | 1690503.0 | 0.5921970101147542 | -0.026984025733786554 | 0.5134920128668933 | 0.001 | 0.001 |
| Vermis.VIII | mci | 1228702.0 | 1.1512890832801015e-06 | 0.25356102509894685 | 0.3732194874505266 | 0.01 | 0.003 |
| Vermis.IX | mci | 1367542.0 | 0.0008498475789407344 | 0.16921544148692202 | 0.415392279256539 | 0.004 | 0.001 |
| Vermis.VII | mci | 1313465.0 | 5.917767680466657e-05 | 0.20206732945139527 | 0.39896633527430236 | 0.006 | 0.001 |
| Vermis.VI | mci | 1389097.5 | 0.0026242090022740084 | 0.1561204312049499 | 0.42193978439752505 | 0.007 | 0.003 |
| Vermis.X | mci | 1318949.0 | 8.231486429515832e-05 | 0.198735788249088 | 0.400632105875456 | 0.014 | 0.004 |
| Right.V | mci | 1415264.5 | 0.007402402359824727 | 0.1402239252529487 | 0.42988803737352566 | 0.007 | 0.005 |
| Right.IV | mci | 1494278.5 | 0.0782693389849359 | 0.09222275884902664 | 0.4538886205754867 | 0.008 | 0.006 |
| Right.IX | mci | 1476492.5 | 0.04875899712638207 | 0.10302779018094443 | 0.4484861049095278 | 0.005 | 0.003 |
| Right.I.III | mci | 1543447.5 | 0.22147732738792947 | 0.062352490910250635 | 0.4688237545448747 | 0.003 | 0.002 |
| Right.Crus.II | mci | 1288274.5 | 3.2805096978085496e-05 | 0.21737060965867494 | 0.39131469517066253 | 0.007 | 0.003 |
| Right.VIIB | mci | 1325571.0 | 0.00019638129312931043 | 0.19471290972215893 | 0.40264354513892053 | 0.007 | 0.003 |
| Right.VI | mci | 1364640.0 | 0.0010988147476025346 | 0.17097841241491174 | 0.41451079379254413 | 0.008 | 0.004 |
| Right.Crus.I | mci | 1095997.0 | 1.7849042592737674e-10 | 0.33417958367885015 | 0.3329102081605749 | 0.02 | 0.006 |
| Right.Crus.II | mci | 1347947.5 | 0.0005309680828844517 | 0.18111914026310916 | 0.4094404298684454 | 0.005 | 0.003 |
| Right.VIIIB | mci | 1567709.5 | 0.362520957802113 | 0.04761327634964174 | 0.47619336182517913 | 0.004 | 0.003 |
| Right.X | mci | 1306392.0 | 6.433563109461754e-05 | 0.2063641913995936 | 0.3968179043002032 | 0.014 | 0.005 |
| Left. Crus.I | ad | 1767894.0 | 0.032323920783197624 | 0.1025486000593937 | 0.44872569997030315 | 0.009 | 0.007 |
| Left.Crus.II | ad | 1757472.0 | 0.024339448644129253 | 0.10783921052030432 | 0.44608039473984784 | 0.007 | 0.004 |
| Left.VIIB | ad | 1696749.5 | 0.0037693249542063807 | 0.1386643010703562 | 0.4306678494648219 | 0.007 | 0.004 |
| Left.VI | ad | 1963661.0 | 0.9472540110533848 | 0.0031696960005686003 | 0.4984151519997157 | 0.004 | 0.005 |
| Left.VIIIA | ad | 1779872.0 | 0.043806156233989105 | 0.09646810379180726 | 0.45176594810409637 | 0.006 | 0.003 |
| Left..VI | ad | 1977987.0 | 0.9317302369747326 | -0.00410273591873711 | 0.5020513679593686 | 0.004 | 0.005 |
| Left.X | ad | 1660079.5 | 0.0007525304862329802 | 0.15727941195133777 | 0.4213602940243311 | 0.01 | 0.004 |
| Left.IV | ad | 2021532.5 | 0.5842131569889503 | -0.02620811663506606 | 0.513104058317533 | 0.006 | 0.006 |
| Left.VIIIA | ad | 1853486.0 | 0.21634568005488874 | 0.059098789027897336 | 0.47045060548605133 | 0.004 | 0.003 |
| Left.IX | ad | 1885044.0 | 0.3678794522748615 | 0.04307872714674055 | 0.4784606364266297 | 0.005 | 0.004 |
| Left.I.III | ad | 2092681.5 | 0.17612375582125317 | -0.06232610202014821 | 0.5311630510100741 | 0.001 | 0.001 |
| Vermis.VIII | ad | 1774686.0 | 0.03765303778750758 | 0.09910071805493159 | 0.4504496409725342 | 0.005 | 0.003 |
| Vermis.IX | ad | 1988549.0 | 0.8383231631102619 | -0.009464415796700765 | 0.5047322078983504 | 0.001 | 0.001 |
| Vermis.VII | ad | 1785129.5 | 0.04151407280757378 | 0.0937991933621164 | 0.4531004033189418 | 0.003 | 0.001 |

| | | | | | | | |
|---|---|---|---|---|---|---|---|
| Vermis.VI | ad | 1902505.0 | 0.470914957290538 | 0.034214847924138425 | 0.4828925760379308 | 0.003 | 0.003 |
| Vermis.X | ad | 1862871.0 | 0.23907437231643103 | 0.05433459989187295 | 0.4728327000540635 | 0.007 | 0.004 |
| Right.V | ad | 2049982.0 | 0.3959068806258438 | -0.04065018363829731 | 0.5203250918191487 | 0.004 | 0.005 |
| Right.IV | ad | 2050084.0 | 0.39546040615434697 | -0.04070196278500737 | 0.5203509813925037 | 0.005 | 0.006 |
| Right.IX | ad | 1890378.0 | 0.3984562336366594 | 0.04037098235701719 | 0.4798145088214914 | 0.004 | 0.003 |
| Right.I.III | ad | 1984159.5 | 0.8767092173689334 | -0.007236135752739292 | 0.5036180678763696 | 0.002 | 0.002 |
| Right.Crus.II | ad | 1727984.5 | 0.010297601617401893 | 0.12280820648711488 | 0.43859589675644256 | 0.005 | 0.003 |
| Right.VIIB | ad | 1709038.0 | 0.005619804913542164 | 0.13242618298851971 | 0.43378690850574014 | 0.006 | 0.003 |
| Right.VI | ad | 2046516.5 | 0.4169055172484093 | -0.03889096174688622 | 0.5194454808734431 | 0.004 | 0.004 |
| Right.Crus.I | ad | 1663493.0 | 0.0011680993266796837 | 0.155546587271975 | 0.4222267063640125 | 0.012 | 0.006 |
| Right.Crus.II | ad | 1734778.5 | 0.012537543606393454 | 0.11935930920526627 | 0.44032034539736686 | 0.005 | 0.003 |
| Right.VIIIB | ad | 1906404.5 | 0.500251810764552 | 0.03223531083986286 | 0.48388234458006857 | 0.004 | 0.003 |
| Right.X | ad | 1657315.5 | 0.0007788749278398287 | 0.15868252529944338 | 0.4206587373502783 | 0.015 | 0.005 |
| Left. Crus.I | bd | 3637239.5 | 0.44202631129825243 | 0.02680537188396548 | 0.48659731405801726 | 0.007 | 0.007 |
| Left.Crus.II | bd | 3420358.5 | 0.014930509722763939 | 0.0848349363766071 | 0.45758253181169645 | 0.006 | 0.004 |
| Left.VIIB | bd | 3167289.0 | 1.1920585928527027e-05 | 0.1525472434545465 | 0.42372637827272674 | 0.007 | 0.004 |
| Left.VI | bd | 3526832.0 | 0.10607443070734011 | 0.056346452668918245 | 0.4718267736655409 | 0.005 | 0.005 |
| Left.VIIIA | bd | 3389405.0 | 0.007498528795948501 | 0.09311698102101118 | 0.4534415094894944 | 0.004 | 0.003 |
| Left..VI | bd | 3580547.5 | 0.22843545632949958 | 0.041974114513411265 | 0.47901294274329437 | 0.005 | 0.005 |
| Left.X | bd | 3151621.0 | 3.953579024737085e-06 | 0.1567394374063944 | 0.4216302812968028 | 0.011 | 0.004 |
| Left.IV | bd | 3389053.5 | 0.007485864783108183 | 0.09321102979392881 | 0.4533944851030356 | 0.007 | 0.006 |
| Left.VIIIA | bd | 3390407.0 | 0.007608939620089663 | 0.09284888181627848 | 0.45357555909186076 | 0.005 | 0.003 |
| Left.IX | bd | 3401266.5 | 0.00978488127454865 | 0.08994326972666322 | 0.4550283651366684 | 0.005 | 0.004 |
| Left.I.III | bd | 3172398.0 | 6.5737193320269285e-06 | 0.15118025858730177 | 0.4244098707063491 | 0.004 | 0.001 |
| Vermis.VIII | bd | 3447518.5 | 0.02537765319886053 | 0.07756789605670755 | 0.4612160519716462 | 0.004 | 0.003 |
| Vermis.IX | bd | 3448963.0 | 0.02223959631855868 | 0.07718139974808846 | 0.46140930012595577 | 0.002 | 0.001 |
| Vermis.VII | bd | 3609145.5 | 0.30533901937602337 | 0.03432231705139033 | 0.48283884147430484 | 0.002 | 0.001 |
| Vermis.VI | bd | 3767772.5 | 0.8140953060130333 | -0.008120569724188176 | 0.5040602848620941 | 0.003 | 0.003 |
| Vermis.X | bd | 3472059.0 | 0.034529060186095334 | 0.07100173983540792 | 0.46449913008229604 | 0.007 | 0.004 |
| Right.V | bd | 3526522.5 | 0.10537097444110065 | 0.05642926375061963 | 0.4717853681246902 | 0.005 | 0.005 |
| Right.IV | bd | 3388808.0 | 0.0074527423548630565 | 0.09327671677472915 | 0.4533616416126354 | 0.008 | 0.006 |
| Right.IX | bd | 3292047.0 | 0.0006152952608546222 | 0.11916648438863953 | 0.44041675780568024 | 0.005 | 0.003 |
| Right.I.III | bd | 3314296.0 | 0.0008526308406704996 | 0.11321345124882187 | 0.44339327437558906 | 0.004 | 0.002 |
| Right.Crus.II | bd | 3541915.0 | 0.13317805997547713 | 0.05231078370186937 | 0.4738446081490653 | 0.004 | 0.003 |
| Right.VIIB | bd | 3271111.0 | 0.0003370696103019153 | 0.1247682058959082 | 0.4376158970520459 | 0.004 | 0.003 |
| Right.VI | bd | 3558407.0 | 0.16948955591636894 | 0.047898116950920056 | 0.47605094152454 | 0.005 | 0.004 |
| Right.Crus.I | bd | 3488583.0 | 0.056193423602370625 | 0.06658051103400808 | 0.46670974448299596 | 0.009 | 0.006 |
| Right.Crus.II | bd | 3236216.5 | 0.00011605772374266403 | 0.1341047205661121 | 0.43294763971694394 | 0.004 | 0.003 |
| Right.VIIIB | bd | 3483093.0 | 0.050541529755931965 | 0.06804943781442963 | 0.4659752810927852 | 0.004 | 0.003 |
| Right.X | bd | 3079462.5 | 3.0257409172816755e-07 | 0.17604645982625722 | 0.4119767700868714 | 0.012 | 0.005 |
| Left. Crus.I | scz | 4415820.0 | 0.1363062736952736 | -0.048973076210430344 | 0.5244865381052152 | 0.005 | 0.007 |
| Left.Crus.II | scz | 4105685.0 | 0.4522824979267108 | 0.02469914434894982 | 0.4876504278255251 | 0.004 | 0.004 |
| Left.VIIB | scz | 3683842.0 | 0.00014298939433215997 | 0.12490747471292218 | 0.4375462626435389 | 0.006 | 0.004 |
| Left.VI | scz | 4000439.5 | 0.1305659142126117 | 0.04970009454445257 | 0.4751499527277737 | 0.005 | 0.005 |
| Left.VIIIA | scz | 3809508.0 | 0.0037921043639762737 | 0.0950556577015721 | 0.45247217114921395 | 0.004 | 0.003 |
| Left..VI | scz | 4059124.0 | 0.2764708599129304 | 0.0357596575495408 | 0.4821201712252296 | 0.006 | 0.005 |
| Left.X | scz | 3918513.5 | 0.030810488666193337 | 0.06916152373350815 | 0.4654192381332459 | 0.007 | 0.004 |

| | | | | | | | |
|---|---|---|---|---|---|---|---|
| Left.IV | scz | 3978229.0 | 0.0943273793063513 | 0.054976173847769205 | 0.4725119130761154 | 0.007 | 0.006 |
| Left.VIIIA | scz | 3984406.0 | 0.10281631845773938 | 0.053508834442686615 | 0.4732455827786567 | 0.004 | 0.003 |
| Left.IX | scz | 3984401.5 | 0.10308655673470188 | 0.05350990341262718 | 0.4732450482936864 | 0.005 | 0.004 |
| Left.I.III | scz | 3928305.5 | 0.034542599073410646 | 0.06683544514283812 | 0.46658227742858094 | 0.003 | 0.001 |
| Vermis.VIII | scz | 4134324.5 | 0.5843396717345731 | 0.017895863323878936 | 0.49105206833806053 | 0.005 | 0.003 |
| Vermis.IX | scz | 4178286.5 | 0.8148604916374413 | 0.007452739651183204 5 | 0.4962736301744084 | 0.001 | 0.001 |
| Vermis.VII | scz | 4372141.0 | 0.221401446586735 | -0.038597178869552495 | 0.5192985894347762 | 0.001 | 0.001 |
| Vermis.VI | scz | 4393148.0 | 0.18067349988444792 | -0.043587368100986845 | 0.5217936840504934 | 0.003 | 0.003 |
| Vermis.X | scz | 4196542.0 | 0.9216118137544531 | 0.0031161661511855954 | 0.4984419169244072 | 0.004 | 0.004 |
| Right.V | scz | 4006040.5 | 0.1409641344563097 | 0.04836958329176233 | 0.47581520835411883 | 0.006 | 0.005 |
| Right.IV | scz | 3975325.5 | 0.0903154084693645 | 0.055665897008309484 | 0.47216705149584526 | 0.007 | 0.006 |
| Right.IX | scz | 3907395.5 | 0.02862125912765443 5 | 0.0718025921333314 | 0.4640987039333343 | 0.005 | 0.003 |
| Right.I.III | scz | 3895378.0 | 0.019673294090097725 | 0.07465733574682987 | 0.46267133212658507 | 0.003 | 0.002 |
| Right.Crus.II | scz | 4153804.0 | 0.6862213514138658 | 0.013268530000047463 | 0.49336573499997627 | 0.003 | 0.003 |
| Right.VIIB | scz | 3884288.0 | 0.018499338542693047 | 0.07729175277813405 | 0.461354123610933 | 0.004 | 0.003 |
| Right.VI | scz | 3959065.5 | 0.07015065445332765 | 0.05952844172688532 | 0.47023577913655734 | 0.005 | 0.004 |
| Right.Crus.I | scz | 4164957.0 | 0.7466829008230806 | 0.010619147389575367 | 0.4946904263052123 | 0.007 | 0.006 |
| Right.Crus.II | scz | 3717070.5 | 0.00036109727790411135 | 0.11701408189735041 | 0.4414929590513248 | 0.005 | 0.003 |
| Right.VIIIB | scz | 3954323.5 | 0.06452525553789634 | 0.06065489849536543 | 0.4696725507523173 | 0.005 | 0.003 |
| Right.X | scz | 3811810.5 | 0.0035412774576973737 | 0.0945087014153162 | 0.4527456492923419 | 0.008 | 0.005 |

b) negative deviation

| ROI | Diagnosis | U-value | P-value | RBC | CLES | Median Clinical | Median Control |
|---|---|---|---|---|---|---|---|
| Left. Crus.I | asd | 9454683.0 | 1.0784925530337901e-29 | 0.2214042369217466 | 0.3892978815391267 | 0.009 | 0.006 |
| Left.Crus.II | asd | 8025198.0 | 2.512213325035142e-67 | 0.3391227225001544 | 0.3304386387499228 | 0.009 | 0.004 |
| Left.VIIB | asd | 5122726.5 | 2.323003650499576e-192 | 0.5781420542276574 | 0.21092897288617132 | 0.013 | 0.002 |
| Left.VI | asd | 8029542.5 | 3.457811513049505e-67 | 0.33876495172215015 | 0.3306175241389249 | 0.006 | 0.003 |
| Left.VIIIA | asd | 4819689.5 | 4.8354726781321815e-209 | 0.603097235089453 | 0.1984513824552735 | 0.013 | 0.002 |
| Left..VI | asd | 9269129.5 | 9.980474726404285e-34 | 0.23668461902703153 | 0.38165769048648424 | 0.006 | 0.003 |
| Left.X | asd | 8727373.5 | 2.946418685858163e-53 | 0.2812983756407881 | 0.35935081217960596 | 0.004 | 0.0 |
| Left.IV | asd | 8976224.0 | 1.4527347053714118e-40 | 0.26080546805838634 | 0.36959726597080683 | 0.008 | 0.005 |
| Left.VIIIA | asd | 6801497.0 | 2.8829818166137747e-112 | 0.4398948386963951 | 0.28005258065180244 | 0.011 | 0.003 |
| Left.IX | asd | 7526578.5 | 2.7276242173393633e-84 | 0.38018417639429314 | 0.30990791180285343 | 0.01 | 0.003 |
| Left.I.III | asd | 10013713.0 | 3.4739064902406426e-20 | 0.17536796162477097 | 0.4123160191876145 | 0.003 | 0.002 |
| Vermis.VIII | asd | 7972568.0 | 2.5997484942991593e-69 | 0.34345681757354907 | 0.32827159121322547 | 0.009 | 0.003 |
| Vermis.IX | asd | 9209867.5 | 1.094602753333717e-36 | 0.2415648611368456 | 0.3792175694315772 | 0.004 | 0.001 |
| Vermis.VII | asd | 9736861.0 | 3.0971393862111247e-25 | 0.19816680048586666 | 0.40091659975706667 | 0.005 | 0.002 |
| Vermis.VI | asd | 11554857.0 | 0.012935235684121003 | 0.04845432647767278 | 0.4757728367611636 | 0.005 | 0.004 |
| Vermis.X | asd | 10521525.0 | 2.3353531668496838e-14 | 0.13354950281020317 | 0.4332252485948984 | 0.004 | 0.0 |
| Right.V | asd | 9559158.5 | 1.3772598872148813e-27 | 0.21280065056718755 | 0.3935996747164062 | 0.005 | 0.003 |
| Right.IV | asd | 9065529.5 | 2.1372701759389112e-38 | 0.25345113540444275 | 0.3732744322977786 | 0.009 | 0.005 |
| Right.IX | asd | 8026762.0 | 2.1621682003350696e-67 | 0.3389939266670784 | 0.3305030366664608 | 0.009 | 0.004 |
| Right.I.III | asd | 10148872.5 | 1.3858676332145637e-17 | 0.1642375393737261 | 0.41788123031313695 | 0.004 | 0.002 |
| Right.Crus.II | asd | 7766158.5 | 7.883755159966362e-76 | 0.36045469705391886 | 0.31977265147304057 | 0.009 | 0.003 |
| Right.VIIB | asd | 5493137.5 | 2.987033143786519e-173 | 0.547638605809812 | 0.22618069709509397 | 0.009 | 0.001 |
| Right.VI | asd | 8126135.0 | 3.66781053185991e-64 | 0.3308105326004158 | 0.3345947336997921 | 0.006 | 0.003 |

| | | | | | | | |
|---|---|---|---|---|---|---|---|
| Right.Crus.I | asd | 9884565.0 | 1.9495632546794576e-21 | 0.1860033351862146 | 0.4069983324068927 | 0.008 | 0.006 |
| Right.Crus.II | asd | 5472732.0 | 2.736820997434607e-174 | 0.5493190043851522 | 0.22534049780742388 | 0.007 | 0.002 |
| Right.VIIIB | asd | 6910303.0 | 7.71938090848068e-108 | 0.4309346344677084 | 0.2845326827661458 | 0.008 | 0.003 |
| Right.X | asd | 8770636.0 | 5.662471441839031e-51 | 0.2777356967862804 | 0.3611321516068598 | 0.005 | 0.002 |
| Left. Crus.I | mci | 728714.0 | 1.9890518136775664e-26 | 0.5573047564372435 | 0.22134762178137823 | 0.025 | 0.006 |
| Left.Crus.II | mci | 852278.0 | 3.3909562576477886e-20 | 0.48223937402989514 | 0.25888031298505243 | 0.013 | 0.004 |
| Left.VIIB | mci | 882757.0 | 7.849023178496977e-19 | 0.46372331926966104 | 0.2681383403651695 | 0.007 | 0.002 |
| Left.VI | mci | 1029009.5 | 8.293548828237584e-13 | 0.37487462676593253 | 0.31256268661703374 | 0.007 | 0.003 |
| Left.VIIIA | mci | 1028418.0 | 7.54321661291927e-13 | 0.37523396422420474 | 0.31238301788789763 | 0.006 | 0.002 |
| Left..VI | mci | 1068648.5 | 2.0903847504338166e-11 | 0.35079385329433166 | 0.32460307335283417 | 0.008 | 0.003 |
| Left.X | mci | 1003432.0 | 1.4311728436449705e-15 | 0.390413010263747 | 0.3047934948681265 | 0.013 | 0.0 |
| Left.IV | mci | 1215389.5 | 5.862934887740937e-07 | 0.2616483960427317 | 0.36917580197863414 | 0.009 | 0.005 |
| Left.VIIIA | mci | 906124.5 | 8.469362932465188e-18 | 0.4495275152862701 | 0.27523624235686495 | 0.012 | 0.003 |
| Left.IX | mci | 1020263.5 | 3.731956599370989e-13 | 0.38018783963161074 | 0.30990608018419463 | 0.01 | 0.003 |
| Left.I.III | mci | 1310337.0 | 6.334569566927225e-05 | 0.20396759584104096 | 0.3980162020794795 | 0.004 | 0.002 |
| Vermis.VIII | mci | 1171110.5 | 3.3633945610809005e-08 | 0.28854797899257933 | 0.35572601050371033 | 0.009 | 0.003 |
| Vermis.IX | mci | 1162867.0 | 9.127360594494129e-09 | 0.29355592208178805 | 0.353222038959106 | 0.005 | 0.001 |
| Vermis.VII | mci | 1018907.0 | 8.816620522525714e-14 | 0.381011916152568 | 0.309494041923716 | 0.014 | 0.002 |
| Vermis.VI | mci | 1142715.0 | 4.690663270608267e-09 | 0.3057983032467947 | 0.34710084837660266 | 0.013 | 0.004 |
| Vermis.X | mci | 1182965.5 | 1.8333596393044782e-09 | 0.2813460422760672 | 0.3593269788619664 | 0.007 | 0.0 |
| Right.V | mci | 1191440.0 | 1.3259614758161682e-07 | 0.2761977662149889 | 0.36190111689250554 | 0.007 | 0.003 |
| Right.IV | mci | 1138129.5 | 3.835972348420178e-09 | 0.308584003863713 | 0.3457079980681435 | 0.01 | 0.005 |
| Right.IX | mci | 1080530.5 | 5.209104056166783e-11 | 0.34357551402266595 | 0.328212242988667 | 0.009 | 0.004 |
| Right.I.III | mci | 1103303.5 | 1.5366473827689796e-10 | 0.32974087000367536 | 0.3351295649981623 | 0.006 | 0.002 |
| Right.Crus.II | mci | 871272.5 | 2.562049971963591e-19 | 0.47070017647934337 | 0.2646499117603283 | 0.011 | 0.003 |
| Right.VIIB | mci | 873734.0 | 2.7252221973748676e-19 | 0.4692048102011743 | 0.26539759489941284 | 0.006 | 0.001 |
| Right.VI | mci | 1074928.5 | 3.494151600305733e-11 | 0.34697874046601485 | 0.3265106297669926 | 0.006 | 0.003 |
| Right.Crus.I | mci | 724351.0 | 1.1541721494805018e-26 | 0.5599552878496554 | 0.22002235607517232 | 0.021 | 0.006 |
| Right.Crus.II | mci | 1016061.5 | 2.4001801775757536e-13 | 0.3827405632151438 | 0.3086297183924281 | 0.005 | 0.002 |
| Right.VIIIB | mci | 889368.0 | 1.5160530446088786e-18 | 0.459707123265202 | 0.270146438367399 | 0.008 | 0.003 |
| Right.X | mci | 855099.0 | 2.4195198008196117e-22 | 0.4805256107673662 | 0.2597371946163169 | 0.013 | 0.002 |
| Left. Crus.I | ad | 1226526.0 | 3.370478174177334e-15 | 0.3773679441394382 | 0.3113160279302809 | 0.017 | 0.006 |
| Left.Crus.II | ad | 1463585.5 | 8.08748770032565e-08 | 0.2570273693401458 | 0.3714863153299271 | 0.007 | 0.004 |
| Left.VIIB | ad | 1396259.5 | 1.1626397850652296e-09 | 0.2912046520009848 | 0.3543976739995076 | 0.004 | 0.002 |
| Left.VI | ad | 1604888.5 | 0.00010980261058991617 | 0.1852964990697521 | 0.40735175046512395 | 0.004 | 0.003 |
| Left.VIIIA | ad | 1581673.0 | 3.832212247738555e-05 | 0.19708158515258345 | 0.4014592074237083 | 0.004 | 0.002 |
| Left..VI | ad | 1646506.5 | 0.0006071287635868816 | 0.16416959193463643 | 0.4179152040326818 | 0.006 | 0.003 |
| Left.X | ad | 1326662.0 | 2.8708130403260713e-13 | 0.3265350359535104 | 0.3367324820232448 | 0.009 | 0.0 |
| Left.IV | ad | 1772037.5 | 0.035988780115389564 | 0.10044519913396843 | 0.4497774004330158 | 0.005 | 0.005 |
| Left.VIIIA | ad | 1459700.5 | 6.173288547906649e-08 | 0.2589995456633696 | 0.3705002271683152 | 0.007 | 0.003 |
| Left.IX | ad | 1660754.0 | 0.0010410736052588001 | 0.1569370096527497 | 0.42153149517362515 | 0.005 | 0.003 |
| Left.I.III | ad | 1698630.5 | 0.003147170962561148 | 0.13770943268837832 | 0.43114528365581084 | 0.003 | 0.002 |
| Vermis.VIII | ad | 1765430.0 | 0.02986646561601622 | 0.10379942179952839 | 0.4481002891002358 | 0.005 | 0.003 |
| Vermis.IX | ad | 1571821.0 | 1.5222931474728608e-05 | 0.2020828415583492 | 0.3989585792208254 | 0.003 | 0.001 |
| Vermis.VII | ad | 1504206.0 | 4.1973047555922893e-07 | 0.23640683180153355 | 0.3817965840992332 | 0.005 | 0.002 |
| Vermis.VI | ad | 1538269.0 | 4.442064015190405e-06 | 0.21911513499381952 | 0.39044243250309024 | 0.009 | 0.004 |
| Vermis.X | ad | 1706943.5 | 0.0018096192136550428 | 0.13348943223150356 | 0.4332552838842482 | 0.004 | 0.0 |

| | | | | | | | |
|---|---|---|---|---|---|---|---|
| Right.V | ad | 1598550.5 | 8.249686627187521e-05 | 0.1885139131074849 | 0.40574304344625756 | 0.005 | 0.003 |
| Right.IV | ad | 1681883.0 | 0.002271933819992312 | 0.1462111117033562 | 0.4268944441483219 | 0.007 | 0.005 |
| Right.IX | ad | 1627766.0 | 0.0002847268436178708 | 0.17368299486523464 | 0.4131585025673827 | 0.006 | 0.004 |
| Right.I.III | ad | 1591968.5 | 4.641093420350435e-05 | 0.19185519098636739 | 0.4040724045068163 | 0.004 | 0.002 |
| Right.Crus.II | ad | 1399165.0 | 1.4650804495960575e-09 | 0.2897297077777863 | 0.35513514611110686 | 0.007 | 0.003 |
| Right.VIIB | ad | 1374698.0 | 2.568663289789874e-10 | 0.3021501036852031 | 0.34892494815739844 | 0.005 | 0.001 |
| Right.VI | ad | 1429275.0 | 1.0099878449722022e-08 | 0.274444706724436 | 0.362777646637782 | 0.005 | 0.003 |
| Right.Crus.I | ad | 1131454.0 | 6.467504602462141e-19 | 0.42563016998281644 | 0.2871849150085918 | 0.017 | 0.006 |
| Right.Crus.II | ad | 1517082.0 | 1.5084083977265932e-06 | 0.22987047598742072 | 0.38506476200628964 | 0.003 | 0.002 |
| Right.VIIIB | ad | 1464310.5 | 8.080017341764651e-08 | 0.25665933128754936 | 0.3716703343562253 | 0.005 | 0.003 |
| Right.X | ad | 1412686.5 | 3.8689871450910076e-10 | 0.2828656711871892 | 0.3585671644064054 | 0.007 | 0.002 |
| Left. Crus.I | bd | 2810758.0 | 1.1523305104721719e-12 | 0.24794213124151743 | 0.3760289343792413 | 0.01 | 0.006 |
| Left.Crus.II | bd | 2820328.0 | 1.947084935781223e-12 | 0.24538154302865145 | 0.3773092284856743 | 0.006 | 0.004 |
| Left.VIIB | bd | 2667712.5 | 2.061291867768903e-16 | 0.28621596835787233 | 0.35689201582106383 | 0.004 | 0.002 |
| Left.VI | bd | 3120926.0 | 2.231231333003012e-06 | 0.1649523167370026 | 0.4175238416314987 | 0.004 | 0.003 |
| Left.VIIIA | bd | 2480820.5 | 4.827213403967101e-22 | 0.3362215537579709 | 0.33188922312101454 | 0.005 | 0.002 |
| Left..VI | bd | 3411347.5 | 0.012294250880725903 | 0.08724595627066511 | 0.45637702186466744 | 0.005 | 0.003 |
| Left.X | bd | 2645302.5 | 2.9062297393560366e-19 | 0.29221207931401927 | 0.35389396034299037 | 0.007 | 0.0 |
| Left.IV | bd | 3074785.5 | 3.651323711237656e-07 | 0.17729785701241962 | 0.4113510714937902 | 0.007 | 0.005 |
| Left.VIIIA | bd | 2363717.0 | 4.7329434684190855e-26 | 0.3675542435997 | 0.31622287820015 | 0.009 | 0.003 |
| Left.IX | bd | 2550186.5 | 7.481601998826521e-20 | 0.31766170402195626 | 0.34116914798902187 | 0.008 | 0.003 |
| Left.I.III | bd | 2960628.0 | 9.207540376218282e-10 | 0.20784230308454554 | 0.39607884845772723 | 0.004 | 0.002 |
| Vermis.VIII | bd | 3004947.0 | 1.75412790864861e-08 | 0.19598413077461807 | 0.40200793461269096 | 0.006 | 0.003 |
| Vermis.IX | bd | 3347253.5 | 0.0021377535994224995 | 0.10439520819495252 | 0.44780239590252374 | 0.001 | 0.001 |
| Vermis.VII | bd | 3052032.5 | 6.938598169501866e-08 | 0.18338574244683337 | 0.4083071287765833 | 0.005 | 0.002 |
| Vermis.VI | bd | 3260953.5 | 0.00024327398647466493 | 0.12748598800376465 | 0.4362570059981177 | 0.006 | 0.004 |
| Vermis.X | bd | 3565692.0 | 0.13997989841872296 | 0.04594891265303824 | 0.4770255436734809 | 0.0 | 0.0 |
| Right.V | bd | 3497135.5 | 0.0650349646006968 | 0.06429216926906178 | 0.4678539153654691 | 0.004 | 0.003 |
| Right.IV | bd | 2980987.5 | 6.414580513699847e-09 | 0.20239483226742494 | 0.39880258386628753 | 0.007 | 0.005 |
| Right.IX | bd | 2520258.5 | 8.794059643139763e-21 | 0.32566936170582805 | 0.337165319147086 | 0.009 | 0.004 |
| Right.I.III | bd | 2960744.0 | 1.3482873678197827e-09 | 0.2078112656516623 | 0.39609436717416885 | 0.005 | 0.002 |
| Right.Crus.II | bd | 2748748.0 | 3.248330720753739e-14 | 0.2645337796302131 | 0.36773311018489346 | 0.007 | 0.003 |
| Right.VIIB | bd | 2373446.5 | 9.245964358303408e-26 | 0.36495097891661965 | 0.3175245105416902 | 0.004 | 0.001 |
| Right.VI | bd | 2994667.0 | 1.1940907409695857e-08 | 0.19873468948185546 | 0.40063265525907227 | 0.004 | 0.003 |
| Right.Crus.I | bd | 2782917.5 | 2.3958453248521145e-13 | 0.25539124891552933 | 0.37230437554223533 | 0.01 | 0.006 |
| Right.Crus.II | bd | 2483659.0 | 5.141448030283586e-22 | 0.3354620731265999 | 0.33226896343670004 | 0.004 | 0.002 |
| Right.VIIIB | bd | 2338159.0 | 5.668606791873275e-27 | 0.3743926462689193 | 0.31280367686554034 | 0.007 | 0.003 |
| Right.X | bd | 2620160.5 | 1.042297857373536e-19 | 0.29893917532738135 | 0.3505304123363093 | 0.005 | 0.002 |
| Left. Crus.I | scz | 2521824.5 | 3.26146719426983e-34 | 0.4009434253597678 | 0.2995282873201161 | 0.017 | 0.006 |
| Left.Crus.II | scz | 2284297.0 | 5.224264025023573e-44 | 0.45736781592812725 | 0.2713160920359364 | 0.011 | 0.004 |
| Left.VIIB | scz | 2379154.5 | 4.972513338779174e-40 | 0.4348345234532005 | 0.28258273827339975 | 0.006 | 0.002 |
| Left.VI | scz | 2670170.5 | 9.46065310791141e-29 | 0.36570399984796875 | 0.3171480000760156 | 0.006 | 0.003 |
| Left.VIIIA | scz | 2168004.5 | 2.4094467557497722e-49 | 0.4849929685532799 | 0.25750351572336005 | 0.009 | 0.002 |
| Left..VI | scz | 2962926.5 | 1.9934567638829173e-19 | 0.29616014119905165 | 0.3519199294004742 | 0.007 | 0.003 |
| Left.X | scz | 2363807.0 | 3.3058827280163724e-46 | 0.43848030482271727 | 0.28075984758864136 | 0.011 | 0.0 |
| Left.IV | scz | 2701309.0 | 1.1259942538807981e-27 | 0.3583070841825705 | 0.32084645790871474 | 0.009 | 0.005 |
| Left.VIIIA | scz | 2099793.5 | 1.259780750302447e-52 | 0.5011964149123682 | 0.2494017925438159 | 0.012 | 0.003 |

| | | | | | | | |
|---|---|---|---|---|---|---|---|
| Left.IX | scz | 2254898.0 | 2.150154009694198e-45 | 0.464351515324278 | 0.267824242337861 | 0.012 | 0.003 |
| Left.I.III | scz | 2625555.5 | 6.720380531556424e-32 | 0.3763022429364842 | 0.3118488785317579 | 0.008 | 0.002 |
| Vermis.VIII | scz | 2597859.0 | 1.710201756626925e-31 | 0.3828815153717877 | 0.30855924231410614 | 0.01 | 0.003 |
| Vermis.IX | scz | 3150349.5 | 4.2412834123947375e-15 | 0.2516380182722595 | 0.37418099086387024 | 0.004 | 0.001 |
| Vermis.VII | scz | 2838051.0 | 2.9898088349790823e-24 | 0.3258241758241758 | 0.3370879120879121 | 0.009 | 0.002 |
| Vermis.VI | scz | 3016406.0 | 5.036426412902908e-18 | 0.2834561461020605 | 0.35827192694896975 | 0.011 | 0.004 |
| Vermis.X | scz | 3353684.5 | 4.5068554482370154e-12 | 0.20333601763562859 | 0.3983319911821857 | 0.004 | 0.0 |
| Right.V | scz | 3248554.5 | 3.678469489631541e-12 | 0.22830953093599005 | 0.385845234532005 | 0.005 | 0.003 |
| Right.IV | scz | 2776382.0 | 3.837942954886038e-25 | 0.340473577438558 | 0.329763211280721 | 0.01 | 0.005 |
| Right.IX | scz | 2407327.0 | 7.580654347040218e-39 | 0.42814217775307273 | 0.28592891112346364 | 0.012 | 0.004 |
| Right.I.III | scz | 2774066.5 | 5.167957036332498e-26 | 0.34102362186019775 | 0.3294881890699011 | 0.006 | 0.002 |
| Right.Crus.II | scz | 2483983.5 | 1.0682151472749574e-35 | 0.40993251236441897 | 0.2950337438177905 | 0.01 | 0.003 |
| Right.VIIB | scz | 2187123.5 | 1.3118386022734672e-48 | 0.4804512715991315 | 0.25977436420043426 | 0.007 | 0.001 |
| Right.VI | scz | 2757578.5 | 9.191179612022028e-26 | 0.3449403277224289 | 0.32752983613878556 | 0.006 | 0.003 |
| Right.Crus.I | scz | 2541855.5 | 1.907904596254188e-33 | 0.39618508383099826 | 0.30190745808450087 | 0.015 | 0.006 |
| Right.Crus.II | scz | 2186667.5 | 1.2577882261050228e-48 | 0.4805595938864421 | 0.25972020305677895 | 0.007 | 0.002 |
| Right.VIIIB | scz | 2056391.0 | 9.492218916856934e-55 | 0.5115066299891202 | 0.24424668500543986 | 0.012 | 0.003 |
| Right.X | scz | 2351921.5 | 7.157748288410067e-46 | 0.4413036919846258 | 0.2793481540076871 | 0.011 | 0.002 |

**Supplementary Table 5**. Task atlas based case-control analyses of extreme deviations

a) positive deviation

| ROI | Diagnosis | U-value | P-value | RBC | CLES | Median Clinical | Median Control |
|---|---|---|---|---|---|---|---|
| 10: Autobiographical recall/visual letter recognition/interference resolution | asd | 8774154.0 | 1.183051587029169e-45 | 0.2774459885121364 | 0.3612770057439318 | 0.021 | 0.007 |
| 3: Saccades/visual working memory/visual letter recognition | asd | 10351921.0 | 4.684747523023595e-14 | 0.14751643917402668 | 0.42624178041298666 | 0.013 | 0.009 |
| 4: Action Observation/divided attention/motor planning | asd | 7865053.5 | 1.6714361997285894e-72 | 0.35231066641961584 | 0.3238446667901921 | 0.022 | 0.007 |
| 5: Divided attention/active maintenance/mental arithmetic | asd | 9297144.5 | 4.492357084238655e-33 | 0.2343775760195994 | 0.3828112119902003 | 0.011 | 0.005 |
| 6: Divided attention/verbal fluency/active maintenance | asd | 8920009.0 | 6.21219342413808e-42 | 0.2654347888744776 | 0.3672826055627612 | 0.012 | 0.005 |
| 9: Verbal Fluency/word comprehension/mental arithmetic | asd | 8760836.5 | 5.171647457032251e-46 | 0.27854268832478946 | 0.36072865583760527 | 0.017 | 0.006 |
| 1: Left-hand presses/ motor planning/ interference resolution | asd | 9372312.5 | 1.9467530561817727e-31 | 0.22818747040536924 | 0.3859062647973154 | 0.011 | 0.006 |
| 2: Right-hand presses/ motor planning/ divided attention | asd | 9389059.0 | 4.443403412045818e-31 | 0.2268083914932164 | 0.3865958042533918 | 0.012 | 0.006 |
| 7: Narrative/ emotion processing/ language processing | asd | 9118822.0 | 3.91133018027069e-37 | 0.24906248327260005 | 0.3754687583637 | 0.01 | 0.005 |
| 8: Word comprehension/ language processing/ narrative | asd | 9893202.0 | 2.7486034640756396e-21 | 0.18529207584460505 | 0.4073539620776975 | 0.009 | 0.005 |
| 10: Autobiographical recall/visual letter recognition/interference resolution | mci | 1215770.5 | 6.038819834196191e-07 | 0.2614169377644532 | 0.3692915311177734 | 0.016 | 0.007 |

| | | | | | | | |
|---|---|---|---|---|---|---|---|
| 3: Saccades/visual working memory/visual letter recognition | mci | 1195081.0 | 1.6968521653498722e-07 | 0.2739858512774249 | 0.36300707436128754 | 0.019 | 0.009 |
| 4: Action Observation/divided attention/motor planning | mci | 1187521.0 | 1.052433288690333e-07 | 0.2785785667204306 | 0.3607107166397847 | 0.015 | 0.007 |
| 5: Divided attention/active maintenance/mental arithmetic | mci | 1377047.5 | 0.0018098224958085505 | 0.16344083081979366 | 0.41827958459010317 | 0.01 | 0.005 |
| 6: Divided attention/verbal fluency/active maintenance | mci | 1272656.5 | 1.4895408399087007e-05 | 0.22685857656196373 | 0.38657071171901813 | 0.012 | 0.005 |
| 9: Verbal Fluency/word comprehension/mental arithmetic | mci | 1281025.5 | 2.2971203577251235e-05 | 0.22177439196639304 | 0.3891128040168035 | 0.016 | 0.006 |
| 1: Left-hand presses/ motor planning/ interference resolution | mci | 1430228.0 | 0.012312317077851779 | 0.13113356843662383 | 0.4344332157816881 | 0.009 | 0.006 |
| 2: Right-hand presses/ motor planning/ divided attention | mci | 1388499.0 | 0.0028176387396316254 | 0.15648402117752125 | 0.4217579894112394 | 0.009 | 0.006 |
| 7: Narrative/ emotion processing/ language processing | mci | 1386607.5 | 0.0026193414862112326 | 0.15763311129133672 | 0.42118344435433164 | 0.011 | 0.005 |
| 8: Word comprehension/ language processing/ narrative | mci | 1170579.0 | 3.502581313137822e-08 | 0.28887086632828807 | 0.35556456683585597 | 0.019 | 0.005 |
| 10: Autobiographical recall/visual letter recognition/interference resolution | ad | 1650376.0 | 0.0007102711911156342 | 0.16220528401115786 | 0.41889735799442107 | 0.013 | 0.007 |
| 3: Saccades/visual working memory/visual letter recognition | ad | 1819781.0 | 0.11169553338330361 | 0.07620875118343273 | 0.46189562440828363 | 0.011 | 0.009 |
| 4: Action Observation/divided attention/motor planning | ad | 1698366.0 | 0.00401432177797008 | 0.1378437031227394 | 0.4310781484386303 | 0.012 | 0.007 |
| 5: Divided attention/active maintenance/mental arithmetic | ad | 1842823.5 | 0.178142835801824 | 0.06451148659453121 | 0.4677442567027344 | 0.006 | 0.005 |
| 6: Divided attention/verbal fluency/active maintenance | ad | 1871925.0 | 0.29921082687491574 | 0.04973843916330989 | 0.47513078041834506 | 0.008 | 0.005 |
| 9: Verbal Fluency/word comprehension/mental arithmetic | ad | 1760476.0 | 0.02646457149534109 | 0.10631426388582188 | 0.44684286805708906 | 0.011 | 0.006 |
| 1: Left-hand presses/ motor planning/ interference resolution | ad | 1992368.0 | 0.8118813734736043 | -0.011403087966171066 | 0.5057015439830855 | 0.006 | 0.006 |
| 2: Right-hand presses/ motor planning/ divided attention | ad | 2027636.0 | 0.5407487418643313 | -0.02930648939923497 | 0.5146532446996175 | 0.006 | 0.006 |
| 7: Narrative/ emotion processing/ language processing | ad | 1815370.0 | 0.1015197854851913 | 0.07844794545929878 | 0.4607760272703506 | 0.007 | 0.005 |
| 8: Word comprehension/ language processing/ narrative | ad | 1647206.0 | 0.0006277807843286952 | 0.16381449866871756 | 0.4180927506656412 | 0.008 | 0.005 |
| 10: Autobiographical recall/visual letter recognition/interference resolution | bd | 3488443.0 | 0.056051771683232804 | 0.06661797000472924 | 0.4666910149976354 | 0.008 | 0.007 |
| 3: Saccades/visual working memory/visual letter recognition | bd | 3381893.5 | 0.006367360886388024 | 0.09512678858223811 | 0.45243660570888095 | 0.012 | 0.009 |
| 4: Action Observation/divided attention/motor planning | bd | 3217496.5 | 6.614386996655579e-05 | 0.13911352007968059 | 0.4304432399601597 | 0.01 | 0.007 |
| 5: Divided attention/active maintenance/mental arithmetic | bd | 3562778.0 | 0.1801814932093765 | 0.04672859437219101 | 0.4766357028139045 | 0.007 | 0.005 |
| 6: Divided attention/verbal fluency/active maintenance | bd | 3529743.5 | 0.1110106928816267 | 0.055567439859956935 | 0.47221628007002153 | 0.007 | 0.005 |

| | | | | | | | |
|---|---|---|---|---|---|---|---|
| 9: Verbal Fluency/word comprehension/mental arithmetic | bd | 3490403.0 | 0.05797891918940178 | 0.06609354441463333 | 0.46695322779268333 | 0.007 | 0.006 |
| 1: Left-hand presses/ motor planning/ interference resolution | bd | 3533092.5 | 0.11688633627487093 | 0.05467136776749215 | 0.4726643161162539 | 0.006 | 0.006 |
| 2: Right-hand presses/ motor planning/ divided attention | bd | 3567551.0 | 0.19238510342644644 | 0.04545151103467693 | 0.47727424448266154 | 0.007 | 0.006 |
| 7: Narrative/ emotion processing/ language processing | bd | 3422059.0 | 0.015508764181131543 | 0.08437994366438373 | 0.45781002816780814 | 0.006 | 0.005 |
| 8: Word comprehension/ language processing/ narrative | bd | 3631762.5 | 0.4174502912574205 | 0.0282708203313915 | 0.48586458983430425 | 0.005 | 0.005 |
| 10: Autobiographical recall/visual letter recognition/interference resolution | scz | 4138793.5 | 0.6085961228432353 | 0.016834257398459784 | 0.4915828713007701 | 0.008 | 0.007 |
| 3: Saccades/visual working memory/visual letter recognition | scz | 4051583.5 | 0.25335378479596526 | 0.03755089484661467 | 0.48122455257669267 | 0.01 | 0.009 |
| 4: Action Observation/divided attention/motor planning | scz | 3722535.5 | 0.0004317436613592534 | 0.11571587729175281 | 0.4421420613541236 | 0.01 | 0.007 |
| 5: Divided attention/active maintenance/mental arithmetic | scz | 4282101.0 | 0.6006568784633439 | -0.017208278103219676 | 0.5086041390516098 | 0.005 | 0.005 |
| 6: Divided attention/verbal fluency/active maintenance | scz | 4161767.0 | 0.7292939784877461 | 0.011376928302998301 | 0.49431153584850085 | 0.005 | 0.005 |
| 9: Verbal Fluency/word comprehension/mental arithmetic | scz | 4238085.0 | 0.8372402451809475 | -0.006752326791237273 | 0.5033761633956186 | 0.006 | 0.006 |
| 1: Left-hand presses/ motor planning/ interference resolution | scz | 4131648.0 | 0.5729552985195527 | 0.01853166288963959 | 0.4907341685551802 | 0.006 | 0.006 |
| 2: Right-hand presses/ motor planning/ divided attention | scz | 4114469.5 | 0.49155526402397576 | 0.022612396250528577 | 0.4886938018747357 | 0.007 | 0.006 |
| 7: Narrative/ emotion processing/ language processing | scz | 4162661.0 | 0.7341254260126095 | 0.011164559608139335 | 0.49441772019593033 | 0.005 | 0.005 |
| 8: Word comprehension/ language processing/ narrative | scz | 4278823.0 | 0.6172224956448824 | -0.0164295928887368 | 0.5082147964443684 | 0.004 | 0.005 |

b) negative deviation

| ROI | Diagnosis | U-value | P-value | RBC | CLES | Median Clinical | Median Control |
|---|---|---|---|---|---|---|---|
| 10: Autobiographical recall/visual letter recognition/interference resolution | asd | 7516470.5 | 1.753242034220553e-84 | 0.38101657299322667 | 0.30949171350338667 | 0.016 | 0.007 |
| 3: Saccades/visual working memory/visual letter recognition | asd | 9521135.0 | 2.515778816983356e-28 | 0.2159318963210014 | 0.3920340518394993 | 0.01 | 0.006 |
| 4: Action Observation/divided attention/motor planning | asd | 5691240.5 | 1.9815994898879124e-162 | 0.5313247689045355 | 0.23433761554773228 | 0.014 | 0.004 |
| 5: Divided attention/active maintenance/mental arithmetic | asd | 7794537.5 | 7.458562923815715e-75 | 0.3581176785456941 | 0.32094116072715295 | 0.009 | 0.004 |
| 6: Divided attention/verbal fluency/active maintenance | asd | 7323184.5 | 1.5920648740067808e-91 | 0.3969337286146625 | 0.30153313569266876 | 0.01 | 0.004 |
| 9: Verbal Fluency/word comprehension/mental arithmetic | asd | 7549608.5 | 2.5544689837810175e-83 | 0.3782876495172215 | 0.31085617524138925 | 0.011 | 0.004 |
| 1: Left-hand presses/ motor planning/ interference resolution | asd | 7818916.0 | 4.88900732910926e-74 | 0.356110102320219 | 0.3219449488398905 | 0.009 | 0.004 |

| | | | | | | | |
|---|---|---|---|---|---|---|---|
| 2: Right-hand presses/ motor planning/ divided attention | asd | 8285822.0 | 2.7307362565218676e-59 | 0.31766026393263747 | 0.34116986803368127 | 0.008 | 0.004 |
| 7: Narrative/ emotion processing/ language processing | asd | 7611569.0 | 3.921699906451809e-81 | 0.3731851851851852 | 0.3134074074074074 | 0.009 | 0.004 |
| 8: Word comprehension/ language processing/ narrative | asd | 9024822.0 | 2.3090305663867055e-39 | 0.2568034093014637 | 0.37159829534926814 | 0.008 | 0.004 |
| 10: Autobiographical recall/visual letter recognition/interference resolution | mci | 780671.5 | 1.0660998699766431e-23 | 0.5257404690523272 | 0.2371297654738364 | 0.023 | 0.007 |
| 3: Saccades/visual working memory/visual letter recognition | mci | 734041.5 | 3.8465208070271004e-26 | 0.5540682893046227 | 0.22296585534768862 | 0.019 | 0.006 |
| 4: Action Observation/divided attention/motor planning | mci | 719186.0 | 6.0405886344967384e-27 | 0.5630930359003332 | 0.2184534820498334 | 0.014 | 0.004 |
| 5: Divided attention/active maintenance/mental arithmetic | mci | 814726.0 | 5.392904825592127e-22 | 0.5050522907383276 | 0.2474738546308362 | 0.018 | 0.004 |
| 6: Divided attention/verbal fluency/active maintenance | mci | 827672.5 | 2.3054599008004153e-21 | 0.4971872655421804 | 0.2514063672289098 | 0.012 | 0.004 |
| 9: Verbal Fluency/word comprehension/mental arithmetic | mci | 826435.5 | 1.9893457090440293e-21 | 0.49793874556903195 | 0.251030627215484 | 0.014 | 0.004 |
| 1: Left-hand presses/ motor planning/ interference resolution | mci | 992793.0 | 3.57523831623855e-14 | 0.3968762244963049 | 0.30156188775184756 | 0.01 | 0.004 |
| 2: Right-hand presses/ motor planning/ divided attention | mci | 1016550.5 | 2.874618677201684e-13 | 0.38244349471625094 | 0.30877825264187453 | 0.009 | 0.004 |
| 7: Narrative/ emotion processing/ language processing | mci | 879228.5 | 5.943964055353554e-19 | 0.46586689022741834 | 0.26706655488629083 | 0.014 | 0.004 |
| 8: Word comprehension/ language processing/ narrative | mci | 760639.0 | 9.849015339406619e-25 | 0.5379102537232281 | 0.23104487313838593 | 0.015 | 0.004 |
| 10: Autobiographical recall/visual letter recognition/interference resolution | ad | 1290106.5 | 5.900427145858069e-13 | 0.3450920222041164 | 0.3274539888979418 | 0.014 | 0.007 |
| 3: Saccades/visual working memory/visual letter recognition | ad | 1178422.5 | 5.02336068888448e-17 | 0.401787142019539 | 0.2991064289902305 | 0.014 | 0.006 |
| 4: Action Observation/divided attention/motor planning | ad | 1222386.5 | 2.3706471408980342e-15 | 0.3794693145100906 | 0.3102653427449547 | 0.01 | 0.004 |
| 5: Divided attention/active maintenance/mental arithmetic | ad | 1320844.0 | 6.106955101742073e-12 | 0.3294884778707603 | 0.33525576106461985 | 0.009 | 0.004 |
| 6: Divided attention/verbal fluency/active maintenance | ad | 1269526.0 | 1.1644921087315306e-13 | 0.3555394803302697 | 0.32223025983486514 | 0.01 | 0.004 |
| 9: Verbal Fluency/word comprehension/mental arithmetic | ad | 1294765.5 | 8.421993320873094e-13 | 0.34272693353232775 | 0.3286365332338361 | 0.01 | 0.004 |
| 1: Left-hand presses/ motor planning/ interference resolution | ad | 1613749.0 | 0.00016088580957366095 | 0.18079856642833025 | 0.4096007167858349 | 0.006 | 0.004 |
| 2: Right-hand presses/ motor planning/ divided attention | ad | 1559612.0 | 1.3783937240944986e-05 | 0.20828060236407342 | 0.3958596988179633 | 0.007 | 0.004 |
| 7: Narrative/ emotion processing/ language processing | ad | 1426242.0 | 8.37412421603683e-09 | 0.27598437488102223 | 0.3620078125594889 | 0.008 | 0.004 |
| 8: Word comprehension/ language processing/ narrative | ad | 1219173.5 | 1.7997321462882988e-15 | 0.38110035763145933 | 0.30944982118427034 | 0.013 | 0.004 |
| 10: Autobiographical recall/visual letter recognition/interference resolution | bd | 2588958.0 | 1.217642047088717e-18 | 0.30728784342685367 | 0.34635607828657317 | 0.012 | 0.007 |

| | | | | | | | |
|---|---|---|---|---|---|---|---|
| 3: Saccades/visual working memory/visual letter recognition | bd | 2835764.0 | 4.543044013830166e-12 | 0.24125142394257004 | 0.379374288028715 | 0.01 | 0.006 |
| 4: Action Observation/divided attention/motor planning | bd | 2376773.0 | 1.60658966512996e-25 | 0.3640609270158779 | 0.31796953649206106 | 0.008 | 0.004 |
| 5: Divided attention/active maintenance/mental arithmetic | bd | 2790473.0 | 3.6827969471137436e-13 | 0.2533696685349328 | 0.3733151657325336 | 0.007 | 0.004 |
| 6: Divided attention/verbal fluency/active maintenance | bd | 2580517.0 | 6.821343379701216e-19 | 0.3095463517972613 | 0.34522682410136935 | 0.008 | 0.004 |
| 9: Verbal Fluency/word comprehension/mental arithmetic | bd | 2824682.0 | 2.472295675063121e-12 | 0.24421656903922417 | 0.3778917154803879 | 0.008 | 0.004 |
| 1: Left-hand presses/ motor planning/ interference resolution | bd | 2997104.5 | 1.3386749769744493e-08 | 0.19808250204519295 | 0.4009587489774035 | 0.006 | 0.004 |
| 2: Right-hand presses/ motor planning/ divided attention | bd | 2963118.5 | 2.81737634959627e-09 | 0.20717593475182428 | 0.39641203262408786 | 0.006 | 0.004 |
| 7: Narrative/ emotion processing/ language processing | bd | 2691797.5 | 1.019756331131639e-15 | 0.2797716875734547 | 0.36011415621327264 | 0.008 | 0.004 |
| 8: Word comprehension/ language processing/ narrative | bd | 2825148.5 | 2.5473672065285604e-12 | 0.24409175039749986 | 0.37795412480125007 | 0.008 | 0.004 |
| 10: Autobiographical recall/visual letter recognition/interference resolution | scz | 2189434.0 | 2.891662564251064e-48 | 0.47990241492187014 | 0.26004879253906493 | 0.021 | 0.007 |
| 3: Saccades/visual working memory/visual letter recognition | scz | 2255506.0 | 2.8337719558669815e-45 | 0.46420708560786383 | 0.2678964571960681 | 0.015 | 0.006 |
| 4: Action Observation/divided attention/motor planning | scz | 2069826.0 | 6.244358522739012e-54 | 0.5083151608443437 | 0.24584241957782813 | 0.013 | 0.004 |
| 5: Divided attention/active maintenance/mental arithmetic | scz | 2392876.0 | 2.274734942372842e-39 | 0.4315749965555413 | 0.28421250172222934 | 0.012 | 0.004 |
| 6: Divided attention/verbal fluency/active maintenance | scz | 2281326.5 | 3.943056354713615e-44 | 0.45807345486333817 | 0.2709632725683309 | 0.012 | 0.004 |
| 9: Verbal Fluency/word comprehension/mental arithmetic | scz | 2358728.0 | 8.351649859986073e-41 | 0.43968681556230194 | 0.28015659221884903 | 0.014 | 0.004 |
| 1: Left-hand presses/ motor planning/ interference resolution | scz | 2513241.0 | 1.517777983649289e-34 | 0.4029824261341771 | 0.29850878693291144 | 0.01 | 0.004 |
| 2: Right-hand presses/ motor planning/ divided attention | scz | 2484549.5 | 1.1493731357266558e-35 | 0.4097980597007834 | 0.2951009701496083 | 0.01 | 0.004 |
| 7: Narrative/ emotion processing/ language processing | scz | 2230346.0 | 2.084892716522352e-46 | 0.4701838153200021 | 0.26490809233999896 | 0.013 | 0.004 |
| 8: Word comprehension/ language processing/ narrative | scz | 2575730.0 | 3.60033194128596e-32 | 0.38813823444173634 | 0.30593088277913183 | 0.013 | 0.004 |

**Supplementary Table 6**. Resting-state atlas based case-control analyses of extreme deviations

a) positive deviation

| ROI | Diagnosis | U-value | P-value | RBC | CLES | Median Clinical | Median Control |
|---|---|---|---|---|---|---|---|
| 13: Control C | asd | 9264127.0 | 8.34343388396497e-34 | 0.2370965762872378 | 0.3814517118563811 | 0.013 | 0.006 |
| 16: Default C | asd | 9374309.5 | 2.1407337190945768e-31 | 0.22802301690239435 | 0.3859884915488028 | 0.012 | 0.005 |
| 8: Salience/ Ventral Attention B | asd | 9529450.0 | 3.7168932256150376e-28 | 0.21524715376855452 | 0.39237642311572274 | 0.014 | 0.008 |
| 11: Control A | asd | 10065574.5 | 1.902248491709886e-18 | 0.17109715273917614 | 0.41445142363041193 | 0.013 | 0.007 |
| 7: Salience/Ventral Attention A | asd | 8703662.5 | 1.6572386398135442e-47 | 0.2832509830564305 | 0.35837450847178476 | 0.016 | 0.007 |
| 12 Control B | asd | 8136053.5 | 7.701888576556493e-64 | 0.32999374137895543 | 0.3350031293105223 | 0.016 | 0.005 |
| 17: Temporal Parietal | asd | 9885278.0 | 1.9955148281183934e-21 | 0.18594461943878282 | 0.4070276902806086 | 0.009 | 0.005 |
| 6: Dorsal Attention B | asd | 7455519.0 | 1.1163878622706744e-86 | 0.3860359458958681 | 0.30698202705206595 | 0.02 | 0.006 |
| 9: Limbic B | asd | 8671059.5 | 2.1972103662321266e-48 | 0.2859358491342927 | 0.35703207543285365 | 0.024 | 0.01 |
| 3: Somatomotor A | asd | 8759245.5 | 4.810279970367128e-46 | 0.2786737076153418 | 0.3606631461923291 | 0.01 | 0.004 |
| 15: Default B | asd | 8965386.5 | 7.332386058338036e-41 | 0.2616979391843205 | 0.36915103040783975 | 0.02 | 0.008 |
| 4: Somatomotor B | asd | 9404182.0 | 8.923923246862173e-31 | 0.22556300825561526 | 0.38721849587219237 | 0.008 | 0.004 |
| 14: Default A | asd | 10239898.5 | 2.4028798266563578e-21 | 0.1567415230683713 | 0.42162923846581435 | 0.0 | 0.0 |
| 5: Dorsal Attention A | asd | 10026592.5 | 4.414259764817851e-22 | 0.17430733123340125 | 0.4128463343832994 | 0.005 | 0.0 |
| 2: Visual B | asd | 9946461.0 | 1.4021157713377723e-21 | 0.1809061824470385 | 0.40954690877648076 | 0.006 | 0.002 |
| 10: Limbic A | asd | 9000941.5 | 3.8007822588637894e-45 | 0.2587699750890412 | 0.3706150124554794 | 0.007 | 0.002 |
| 1: Visual A | asd | 10130989.5 | 5.390962532416814e-26 | 0.165710209375579 | 0.4171448953122105 | 0.0 | 0.0 |
| 13: Control C | mci | 1255436.0 | 5.900563713070268e-06 | 0.2373200654887202 | 0.3813399672556399 | 0.015 | 0.006 |
| 16: Default C | mci | 1220702.5 | 8.103554694136512e-07 | 0.2584207376897305 | 0.37078963115513475 | 0.015 | 0.005 |
| 8: Salience/ Ventral Attention B | mci | 1144196.0 | 5.88986994063866e-09 | 0.304898592721518 | 0.347550703639241 | 0.019 | 0.008 |
| 11: Control A | mci | 1199993.0 | 2.1792270884247182e-07 | 0.27100180124355666 | 0.36449909937822167 | 0.02 | 0.007 |
| 7: Salience/Ventral Attention A | mci | 1330858.5 | 0.0002568692228036618 | 0.19150074267124728 | 0.40424962866437636 | 0.012 | 0.007 |
| 12 Control B | mci | 1335532.5 | 0.0003167142784202677 | 0.18866127812354772 | 0.40566936093822614 | 0.011 | 0.005 |
| 17: Temporal Parietal | mci | 1158901.0 | 1.6072892864737352e-08 | 0.2959652751832378 | 0.3520173624083811 | 0.018 | 0.005 |
| 6: Dorsal Attention B | mci | 1221909.5 | 8.702505822513567e-07 | 0.25768748272416064 | 0.3711562586379197 | 0.014 | 0.006 |
| 9: Limbic B | mci | 1496121.5 | 0.0820215731233079 | 0.09110313258428326 | 0.45444843370785837 | 0.011 | 0.01 |
| 3: Somatomotor A | mci | 1481654.5 | 0.05654842609126799 | 0.09989186463639488 | 0.45005406768180256 | 0.005 | 0.004 |
| 15: Default B | mci | 1370572.0 | 0.0013884301152478102 | 0.16737471029746342 | 0.4163126448512683 | 0.017 | 0.008 |
| 4: Somatomotor B | mci | 1478590.5 | 0.05202375370552947 | 0.10175325089530618 | 0.4491233745523469 | 0.006 | 0.004 |
| 14: Default A | mci | 1623364.0 | 0.754126119457112 | 0.013803053912768815 | 0.4930984730436156 | 0.0 | 0.0 |
| 5: Dorsal Attention A | mci | 1444581.0 | 0.011110227894827887 | 0.12241409161738304 | 0.4387929541913085 | 0.003 | 0.0 |
| 2: Visual B | mci | 1634994.5 | 0.8943229380760686 | 0.006737501404848523 | 0.49663124929757574 | 0.001 | 0.002 |
| 10: Limbic A | mci | 1383012.0 | 0.0011165743845108574 | 0.15981738488595665 | 0.4200913075570217 | 0.003 | 0.002 |
| 1: Visual A | mci | 1568250.5 | 0.25859436491303467 | 0.04728461774452719 | 0.4763576911277364 | 0.0 | 0.0 |
| 13: Control C | ad | 1691946.0 | 0.003228804040651361 | 0.14110274353331764 | 0.4294486282333412 | 0.01 | 0.006 |
| 16: Default C | ad | 1638995.0 | 0.00045450837263940263 | 0.16798271997888226 | 0.41600864001055887 | 0.011 | 0.005 |
| 8: Salience/ Ventral Attention B | ad | 1792170.5 | 0.059678946055910946 | 0.0902249093230384 | 0.4548875453384808 | 0.011 | 0.008 |
| 11: Control A | ad | 1617651.0 | 0.0001841313205008928 | 0.17881776024732154 | 0.41059111987633923 | 0.014 | 0.007 |
| 7: Salience/Ventral Attention A | ad | 1853144.5 | 0.21604673926033902 | 0.059272147641637574 | 0.4703639261791812 | 0.009 | 0.007 |
| 12 Control B | ad | 1803933.5 | 0.07865021661642635 | 0.08425355537449775 | 0.4578732223127511 | 0.008 | 0.005 |
| 17: Temporal Parietal | ad | 1724550.5 | 0.009328199247728233 | 0.12455143775968891 | 0.43772428112015555 | 0.008 | 0.005 |
| 6: Dorsal Attention B | ad | 1720349.0 | 0.008187179879772166 | 0.126684281729322 | 0.436657859135339 | 0.01 | 0.006 |

| | | | | | | | |
|---|---|---|---|---|---|---|---|
| 9: Limbic B | ad | 2030923.5 | 0.5179217801927075 | -0.030975351603249823 | 0.5154876758016249 | 0.009 | 0.01 |
| 3: Somatomotor A | ad | 2030212.0 | 0.522827448016372 | -0.0306141666730122 | 0.5153070833365061 | 0.003 | 0.004 |
| 15: Default B | ad | 1832797.0 | 0.14602539356278868 | 0.06960132595226676 | 0.4651993370238666 | 0.011 | 0.008 |
| 4: Somatomotor B | ad | 2035142.0 | 0.48928170114643743 | -0.03311682543066796 | 0.516558412715334 | 0.003 | 0.004 |
| 14: Default A | ad | 2141785.0 | 0.030307593783662212 | -0.08725293859348549 | 0.5436264692967427 | 0.0 | 0.0 |
| 5: Dorsal Attention A | ad | 1881883.0 | 0.3108171926194897 | 0.04468337305606107 | 0.47765831347196946 | 0.003 | 0.0 |
| 2: Visual B | ad | 2143313.5 | 0.05770130865978478 | -0.08802886433609736 | 0.5440144321680487 | 0.001 | 0.002 |
| 10: Limbic A | ad | 1809145.5 | 0.06874615787943493 | 0.08160774250534919 | 0.4591961287473254 | 0.002 | 0.002 |
| 1: Visual A | ad | 2009598.5 | 0.5984888833031767 | -0.020149956469982078 | 0.510074978234991 | 0.0 | 0.0 |
| 13: Control C | bd | 3616187.5 | 0.35220506677379804 | 0.032438130824117395 | 0.4837809345879413 | 0.007 | 0.006 |
| 16: Default C | bd | 3332671.5 | 0.001896036546343656 | 0.10829682755963499 | 0.4458515862201825 | 0.008 | 0.005 |
| 8: Salience/ Ventral Attention B | bd | 3366237.0 | 0.004394300488973525 | 0.09931590554720537 | 0.4503420472263973 | 0.01 | 0.008 |
| 11: Control A | bd | 3130660.0 | 3.0777665850209406e-06 | 0.16234784801557756 | 0.4188260759922112 | 0.014 | 0.007 |
| 7: Salience/Ventral Attention A | bd | 3560056.5 | 0.1734984264307211 | 0.04745677000660209 | 0.47627161499669896 | 0.009 | 0.007 |
| 12 Control B | bd | 3449858.0 | 0.027331564839230486 | 0.07694192989954973 | 0.46152903505022513 | 0.007 | 0.005 |
| 17: Temporal Parietal | bd | 3663972.0 | 0.5729771911677017 | 0.019652715206803673 | 0.49017364239659816 | 0.005 | 0.005 |
| 6: Dorsal Attention B | bd | 3249238.5 | 0.00017942114015609 | 0.13062050116089363 | 0.4346897494195532 | 0.008 | 0.006 |
| 9: Limbic B | bd | 3355455.0 | 0.0033753598153017663 | 0.10220078142088562 | 0.4488996092895572 | 0.013 | 0.01 |
| 3: Somatomotor A | bd | 3452536.5 | 0.0287998817123705 | 0.07622525952043158 | 0.4618873702397842 | 0.005 | 0.004 |
| 15: Default B | bd | 3013234.0 | 2.6813569064336896e-08 | 0.19376682727200367 | 0.40311658636399816 | 0.015 | 0.008 |
| 4: Somatomotor B | bd | 3753094.5 | 0.9042441038802812 | -0.004193264208154135 | 0.5020966321040771 | 0.003 | 0.004 |
| 14: Default A | bd | 3452133.0 | 0.009298701237762632 | 0.0763332216253314 | 0.4618333891873343 | 0.0 | 0.0 |
| 5: Dorsal Attention A | bd | 3367309.0 | 0.0020330947311136835 | 0.09902907685711204 | 0.450485461571444 | 0.003 | 0.0 |
| 2: Visual B | bd | 3539215.0 | 0.11619024213316202 | 0.053033206708634095 | 0.47348339664568295 | 0.004 | 0.002 |
| 10: Limbic A | bd | 3599512.0 | 0.2581209860481757 | 0.036899895583119124 | 0.48155005220844044 | 0.002 | 0.002 |
| 1: Visual A | bd | 3854454.5 | 0.2606248644696191 | -0.03131355901025379 | 0.5156567795051269 | 0.0 | 0.0 |
| 13: Control C | scz | 4321895.0 | 0.4173699455850388 | -0.026661298062076177 | 0.5133306490310381 | 0.006 | 0.006 |
| 16: Default C | scz | 4063076.0 | 0.2894948745825491 | 0.034820864392848816 | 0.4825895678035756 | 0.006 | 0.005 |
| 8: Salience/ Ventral Attention B | scz | 3909198.5 | 0.02992393623211227 | 0.07137429151047825 | 0.4643128542447609 | 0.01 | 0.008 |
| 11: Control A | scz | 3770878.0 | 0.0014879963922808162 | 0.10423217076913571 | 0.44788391461543214 | 0.012 | 0.007 |
| 7: Salience/Ventral Attention A | scz | 3989824.5 | 0.11217497251069168 | 0.05222167585980819 | 0.4738891620700959 | 0.007 | 0.007 |
| 12 Control B | scz | 4070524.5 | 0.31470417700450204 | 0.03305148159233762 | 0.4834742592038312 | 0.006 | 0.005 |
| 17: Temporal Parietal | scz | 4380772.0 | 0.2162682804791214 | -0.04064746321555668 | 0.5203237316077783 | 0.004 | 0.005 |
| 6: Dorsal Attention B | scz | 3685913.0 | 0.00015389560054773494 | 0.12441551099138648 | 0.43779224450430676 | 0.008 | 0.006 |
| 9: Limbic B | scz | 4065457.0 | 0.2973836792249662 | 0.03425526051985195 | 0.482872369740074 | 0.011 | 0.01 |
| 3: Somatomotor A | scz | 4060140.0 | 0.27994212164258125 | 0.03551830789184873 | 0.48224084605407563 | 0.004 | 0.004 |
| 15: Default B | scz | 3753401.5 | 0.0009698100982157236 | 0.10838369369497769 | 0.44580815315251116 | 0.011 | 0.008 |
| 4: Somatomotor B | scz | 4261868.0 | 0.7058913980925721 | -0.012401951701562552 | 0.5062009758507813 | 0.004 | 0.004 |
| 14: Default A | scz | 4131638.0 | 0.5027831822612195 | 0.0185340383783964 | 0.4907329808108018 | 0.0 | 0.0 |
| 5: Dorsal Attention A | scz | 3922449.5 | 0.024144853647616083 | 0.06822653135882706 | 0.46588673432058647 | 0.003 | 0.0 |
| 2: Visual B | scz | 4266192.0 | 0.6730392612605045 | -0.013429113040007934 | 0.506714556520004 | 0.001 | 0.002 |
| 10: Limbic A | scz | 4212637.5 | 0.9816594903923682 | -0.0007073017773406853 | 0.5003536508886703 | 0.002 | 0.002 |
| 1: Visual A | scz | 4280329.5 | 0.5225540025806521 | -0.016787460269950483 | 0.5083937301349752 | 0.0 | 0.0 |

b) negative deviation

| ROI | Diagnosis | U-value | P-value | RBC | CLES | Median Clinical | Median Control |
|---|---|---|---|---|---|---|---|
| 13: Control C | asd | 8193069.0 | 4.384018930744187e-62 | 0.32529849916620346 | 0.33735075041689827 | 0.011 | 0.005 |
| 16: Default C | asd | 8491453.5 | 2.5261571516411616e-53 | 0.30072645296769807 | 0.34963677351615097 | 0.011 | 0.006 |
| 8: Salience/ Ventral Attention B | asd | 8574614.5 | 5.277764796904111e-51 | 0.2938781215901839 | 0.35306093920490805 | 0.01 | 0.006 |
| 11: Control A | asd | 8036010.5 | 1.70022021774991e-67 | 0.33823231013114285 | 0.3308838449344286 | 0.012 | 0.004 |
| 7: Salience/Ventral Attention A | asd | 6999843.5 | 6.003129250113669e-104 | 0.4235609494986927 | 0.28821952525065364 | 0.01 | 0.004 |
| 12 Control B | asd | 6033000.5 | 6.764873877209608e-146 | 0.5031807382702325 | 0.2484096308648838 | 0.012 | 0.003 |
| 17: Temporal Parietal | asd | 9240240.5 | 2.428364316021974e-34 | 0.23906363617647663 | 0.3804681819117617 | 0.007 | 0.004 |
| 6: Dorsal Attention B | asd | 5320617.5 | 1.9814556352706707e-181 | 0.5618456755810841 | 0.21907716220945794 | 0.012 | 0.003 |
| 9: Limbic B | asd | 6684652.0 | 7.45598844516599e-117 | 0.44951705680110354 | 0.27524147159944823 | 0.015 | 0.005 |
| 3: Somatomotor A | asd | 7223156.5 | 2.7377671769432178e-95 | 0.4051710621126964 | 0.2974144689436518 | 0.008 | 0.003 |
| 15: Default B | asd | 7823433.5 | 4.8494141546097234e-74 | 0.3557380849443106 | 0.3221309575278447 | 0.012 | 0.004 |
| 4: Somatomotor B | asd | 8550633.0 | 1.1099570156175814e-51 | 0.2958530047557285 | 0.35207349762213574 | 0.007 | 0.004 |
| 14: Default A | asd | 8826137.5 | 1.9717237925067652e-90 | 0.27316513289275934 | 0.36341743355362033 | 0.0 | 0.0 |
| 5: Dorsal Attention A | asd | 9532239.5 | 1.4962789297579523e-34 | 0.21501743767113413 | 0.39249128116443294 | 0.003 | 0.0 |
| 2: Visual B | asd | 9797934.0 | 5.389716644689699e-24 | 0.1931374220245815 | 0.40343128898770925 | 0.005 | 0.002 |
| 10: Limbic A | asd | 8885717.5 | 3.496170716623814e-49 | 0.2682587033948902 | 0.3658706483025549 | 0.005 | 0.002 |
| 1: Visual A | asd | 9123086.0 | 8.461998251016572e-95 | 0.24871134169188647 | 0.37564432915405677 | 0.0 | 0.0 |
| 13: Control C | mci | 735435.0 | 4.57730788852882e-26 | 0.5532217352080846 | 0.22338913239595767 | 0.02 | 0.005 |
| 16: Default C | mci | 814729.0 | 5.398951485418523e-22 | 0.5050504682321995 | 0.24747476588390027 | 0.018 | 0.006 |
| 8: Salience/ Ventral Attention B | mci | 731371.5 | 2.7642426624366704e-26 | 0.5556903197587002 | 0.2221548401206499 | 0.018 | 0.006 |
| 11: Control A | mci | 756879.5 | 3.98161456196372e-25 | 0.5401941576528551 | 0.22990292117357244 | 0.025 | 0.004 |
| 7: Salience/Ventral Attention A | mci | 853156.5 | 3.759964417589222e-20 | 0.4817056834853607 | 0.25914715825731965 | 0.012 | 0.004 |
| 12 Control B | mci | 926783.0 | 7.351357444893818e-17 | 0.43697743433662295 | 0.2815112828316885 | 0.009 | 0.003 |
| 17: Temporal Parietal | mci | 766249.0 | 1.9292055102206018e-24 | 0.5345021672635375 | 0.23274891636823128 | 0.017 | 0.004 |
| 6: Dorsal Attention B | mci | 808757.0 | 2.7088989544969407e-22 | 0.5086784704313568 | 0.24566076478432158 | 0.011 | 0.003 |
| 9: Limbic B | mci | 1034957.5 | 1.3674625244076881e-12 | 0.3712612046158005 | 0.31436939769209976 | 0.012 | 0.005 |
| 3: Somatomotor A | mci | 1152451.0 | 1.0383098499902822e-08 | 0.299883663358818 | 0.350058168320591 | 0.006 | 0.003 |
| 15: Default B | mci | 835666.0 | 5.046217825591192e-21 | 0.4923311979636531 | 0.25383440101817345 | 0.019 | 0.004 |
| 4: Somatomotor B | mci | 1048244.5 | 4.104610542963405e-12 | 0.3631893249741053 | 0.31840533751294736 | 0.008 | 0.004 |
| 14: Default A | mci | 1564702.0 | 0.1659089082252082 | 0.04944033874313902 | 0.4752798306284305 | 0.0 | 0.0 |
| 5: Dorsal Attention A | mci | 1074726.0 | 1.248045178549605e-13 | 0.3471017596296667 | 0.32644912018516664 | 0.008 | 0.0 |
| 2: Visual B | mci | 1297331.5 | 3.4507954167505966e-05 | 0.21186846365770906 | 0.39406576817114547 | 0.005 | 0.002 |
| 10: Limbic A | mci | 1218436.5 | 9.005257636618471e-08 | 0.2597973373185467 | 0.37010133134072665 | 0.007 | 0.002 |
| 1: Visual A | mci | 1344243.5 | 6.612354292855608e-09 | 0.18336932782936488 | 0.40831533608531756 | 0.0 | 0.0 |
| 13: Control C | ad | 1201574.5 | 3.929381549633656e-16 | 0.3900342909937281 | 0.30498285450313595 | 0.013 | 0.005 |
| 16: Default C | ad | 1307144.0 | 2.1835140563864577e-12 | 0.33644312796809994 | 0.33177843601595003 | 0.013 | 0.006 |
| 8: Salience/ Ventral Attention B | ad | 1135649.0 | 9.630123197226541e-19 | 0.4235006256646894 | 0.2882496871676553 | 0.015 | 0.006 |
| 11: Control A | ad | 1363918.0 | 1.134592452799643e-10 | 0.30762244879829226 | 0.34618877560085387 | 0.01 | 0.004 |
| 7: Salience/Ventral Attention A | ad | 1434769.5 | 1.4284038810700953e-08 | 0.2716554859244481 | 0.36417225703777595 | 0.008 | 0.004 |
| 12 Control B | ad | 1312767.5 | 3.3345630422335237e-12 | 0.3335884217766847 | 0.33320578911165766 | 0.007 | 0.003 |
| 17: Temporal Parietal | ad | 1216667.5 | 1.4520676113468315e-15 | 0.38237250019671 | 0.308813749901645 | 0.011 | 0.004 |
| 6: Dorsal Attention B | ad | 1273699.0 | 1.611973113810795e-13 | 0.3534211040633939 | 0.32328944796830306 | 0.007 | 0.003 |
| 9: Limbic B | ad | 1526389.5 | 2.6055166268892802e-06 | 0.22514562885012224 | 0.3874271855749389 | 0.01 | 0.005 |
| 3: Somatomotor A | ad | 1702969.0 | 0.004678284893455761 | 0.13550704221777188 | 0.43224647889111406 | 0.004 | 0.003 |
| 15: Default B | ad | 1428554.0 | 9.337548731456814e-09 | 0.27481071422225944 | 0.3625946428888703 | 0.009 | 0.004 |
| 4: Somatomotor B | ad | 1532359.5 | 3.541109135555944e-06 | 0.22211502585150045 | 0.3889424870742498 | 0.006 | 0.004 |

| | | | | | | | |
|---|---|---|---|---|---|---|---|
| 14: Default A | ad | 1947938.0 | 0.732509812863366 | 0.011151299174325668 | 0.49442435041283717 | 0.0 | 0.0 |
| 5: Dorsal Attention A | ad | 1409230.0 | 3.031784705340573e-11 | 0.2846203243303611 | 0.35768983783481945 | 0.005 | 0.0 |
| 2: Visual B | ad | 1806460.0 | 0.07613583813069041 | 0.08297100621603581 | 0.4585144968919821 | 0.004 | 0.002 |
| 10: Limbic A | ad | 1485803.0 | 3.2293334499163064e-08 | 0.2457489066731644 | 0.3771255466634178 | 0.005 | 0.002 |
| 1: Visual A | ad | 1839401.5 | 0.021751932376714696 | 0.06624862620278638 | 0.4668756868986068 | 0.0 | 0.0 |
| 13: Control C | bd | 2918731.0 | 3.3338526635661066e-10 | 0.2190524352009975 | 0.39047378239950126 | 0.008 | 0.005 |
| 16: Default C | bd | 2675946.0 | 3.7762586931506303e-16 | 0.2840129795333549 | 0.35799351023332254 | 0.009 | 0.006 |
| 8: Salience/ Ventral Attention B | bd | 2466953.0 | 1.856997809740919e-22 | 0.3399319985899373 | 0.33003400070503136 | 0.011 | 0.006 |
| 11: Control A | bd | 1915702.5 | 9.161359437401114e-45 | 0.4874268295864329 | 0.25628658520678355 | 0.019 | 0.004 |
| 7: Salience/Ventral Attention A | bd | 2711059.5 | 3.3795788959355407e-15 | 0.27461786833038004 | 0.36269106583481 | 0.008 | 0.004 |
| 12 Control B | bd | 2723037.0 | 7.005163533948906e-15 | 0.27141311960314896 | 0.3642934401984255 | 0.006 | 0.003 |
| 17: Temporal Parietal | bd | 3072625.5 | 3.3687593607470075e-07 | 0.17787579541783138 | 0.4110621022910843 | 0.006 | 0.004 |
| 6: Dorsal Attention B | bd | 2424073.5 | 6.795488135491511e-24 | 0.3514050124116286 | 0.3242974937941857 | 0.007 | 0.003 |
| 9: Limbic B | bd | 2719015.0 | 5.461939016959264e-15 | 0.2724892623191517 | 0.36375536884042414 | 0.008 | 0.005 |
| 3: Somatomotor A | bd | 3055972.5 | 1.7000360164711917e-07 | 0.1823315399851101 | 0.40883423000744495 | 0.004 | 0.003 |
| 15: Default B | bd | 2248251.5 | 2.6293943647870012e-30 | 0.398448663483992 | 0.300775668258004 | 0.012 | 0.004 |
| 4: Somatomotor B | bd | 3035028.5 | 7.015208985278428e-08 | 0.18793540200499137 | 0.4060322989975043 | 0.005 | 0.004 |
| 14: Default A | bd | 3470671.0 | 0.002682479529699735 | 0.07137311877370034 | 0.46431344061314983 | 0.0 | 0.0 |
| 5: Dorsal Attention A | bd | 2983812.5 | 1.0014407864922442e-10 | 0.2016389637510878 | 0.3991805181244561 | 0.003 | 0.0 |
| 2: Visual B | bd | 3431633.5 | 0.016264344966841978 | 0.08181815141317317 | 0.4590909242934134 | 0.002 | 0.002 |
| 10: Limbic A | bd | 3207280.0 | 1.1579455518708957e-05 | 0.14184708846805516 | 0.4290764557659724 | 0.003 | 0.002 |
| 1: Visual A | bd | 3774643.5 | 0.6350378855491989 | -0.009959002494366098 | 0.504979501247183 | 0.0 | 0.0 |
| 13: Control C | scz | 2488382.5 | 1.6311972010254434e-35 | 0.4088875348602975 | 0.29555623256985125 | 0.014 | 0.005 |
| 16: Default C | scz | 2318878.0 | 1.6973141335514957e-42 | 0.4491531382581966 | 0.2754234308709017 | 0.016 | 0.006 |
| 8: Salience/ Ventral Attention B | scz | 2263861.0 | 6.667433683510564e-45 | 0.46222236475154765 | 0.2688888176242262 | 0.016 | 0.006 |
| 11: Control A | scz | 1711452.0 | 1.959667811171306e-73 | 0.5934465016177078 | 0.20327674919114608 | 0.024 | 0.004 |
| 7: Salience/Ventral Attention A | scz | 2269135.5 | 1.1437824889078767e-44 | 0.4609694132067673 | 0.26951529339661634 | 0.011 | 0.004 |
| 12 Control B | scz | 2248700.5 | 1.4023148217429138e-45 | 0.46582372448131204 | 0.267088137759344 | 0.01 | 0.003 |
| 17: Temporal Parietal | scz | 2739820.5 | 2.3787196215175824e-26 | 0.3491587206567751 | 0.32542063967161244 | 0.011 | 0.004 |
| 6: Dorsal Attention B | scz | 2086141.0 | 3.7453066713232556e-53 | 0.5044395509376054 | 0.2477802245311973 | 0.011 | 0.003 |
| 9: Limbic B | scz | 2312637.5 | 8.962117258254687e-43 | 0.4506355620168849 | 0.27468221899155754 | 0.015 | 0.005 |
| 3: Somatomotor A | scz | 2609438.0 | 6.312394831481374e-31 | 0.38013093694027544 | 0.3099345315298623 | 0.007 | 0.003 |
| 15: Default B | scz | 1960316.0 | 1.5698390925875416e-59 | 0.5343291382201888 | 0.2328354308899056 | 0.019 | 0.004 |
| 4: Somatomotor B | scz | 2631751.0 | 4.018537477349028e-30 | 0.3748305088772015 | 0.31258474556139926 | 0.008 | 0.004 |
| 14: Default A | scz | 3332227.0 | 1.9143468656536846e-20 | 0.2084332226355573 | 0.39578338868222135 | 0.0 | 0.0 |
| 5: Dorsal Attention A | scz | 2853731.5 | 6.837452146992438e-28 | 0.3220992906790572 | 0.3389503546604714 | 0.005 | 0.0 |
| 2: Visual B | scz | 3079867.5 | 6.380120662122315e-17 | 0.2683809381280199 | 0.36580953093599006 | 0.006 | 0.002 |
| 10: Limbic A | scz | 2945616.0 | 7.634497975433094e-23 | 0.30027223101153067 | 0.34986388449423467 | 0.007 | 0.002 |
| 1: Visual A | scz | 4005498.0 | 0.01441986734511377 | 0.048498453556819365 | 0.4757507732215903 | 0.0 | 0.0 |

**Supplementary Table 7**. Association of extreme deviations and intelligence scores in ASD using anatomical atlas

| Group | ROI | Test | Correlation of % positive deviation | P value of % positive deviation | Correlation of % negative deviation | P value of % negative deviation |
|---|---|---|---|---|---|---|
| asd | Left. Crus.I | FIQ | 0.18892914915578754 | 0.00357681066399968 | -0.06347602031330316 | 0.3315786079882972 |
| asd | Left.Crus.II | FIQ | 0.21673224835868066 | 0.0008030146111996874 | -0.12777034403405046 | 0.04994135009930475 |
| asd | Left.VIIB | FIQ | 0.08625907584031904 | 0.1866468352747465 | -0.10874633254675482 | 0.09558023958839204 |
| asd | Left.VI | FIQ | 0.18419051580263737 | 0.004527018856046755 | -0.13366118727284312 | 0.040203240775133486 |
| asd | Left.VIIIA | FIQ | 0.07186171876678966 | 0.271540759299809 | -0.041454660761523376 | 0.526257212377321 |
| asd | Left..VI | FIQ | 0.21395997417276616 | 0.0009401422863682814 | -0.152551899575718 | 0.01903501429944865 |
| asd | Left.X | FIQ | 0.11338725554170505 | 0.08216685810212024 | -0.11132340613901083 | 0.08793172322196424 |
| asd | Left.IV | FIQ | 0.16657270301802354 | 0.010368703588194108 | -0.1269259602721657 | 0.05148633586752263 |
| asd | Left.VIIIA | FIQ | 0.04479632146609672 | 0.49342965447344533 | -0.04741467047878972 | 0.4684898442565003 |
| asd | Left.IX | FIQ | 0.06765567654048281 | 0.3006623080162698 | -0.06118272646895776 | 0.3493795747703583 |
| asd | Left.I.III | FIQ | 0.10831652817208043 | 0.09690587980893292 | -0.02038845623212543 | 0.7553576268673303 |
| asd | Vermis.VIII | FIQ | 0.08335656136435629 | 0.2019659842724884 | -0.21240040023481896 | 0.0010264531741989028 |
| asd | Vermis.IX | FIQ | 0.07698164422519095 | 0.2387629943014541 | -0.21953045723109935 | 0.0006835154691522269 |
| asd | Vermis.VII | FIQ | 0.08316797744618476 | 0.20299192105714808 | -0.17410664348935212 | 0.007340629459848169 |
| asd | Vermis.VI | FIQ | 0.15812133204656056 | 0.01503542130605377 | -0.15916674454289578 | 0.014372819477392029 |
| asd | Vermis.X | FIQ | 0.1251842564273615 | 0.054798881347280955 | -0.02814438452006452 | 0.6670852124609004 |
| asd | Right.V | FIQ | 0.23548921523501143 | 0.00026233285548516393 | -0.23291438933549047 | 0.00030754501699405894 |
| asd | Right.IV | FIQ | 0.22414638637493858 | 0.0005216845920673173 | -0.12900180683518397 | 0.04775750605202508 |
| asd | Right.IX | FIQ | 0.028247653309175502 | 0.6659370966346126 | -0.13038536457789537 | 0.04539965939147092 |
| asd | Right.I.III | FIQ | 0.16203118972704306 | 0.012686657886670984 | -0.0728320186441599 | 0.26510476659491533 |
| asd | Right.Crus.II | FIQ | 0.22000385296074074 | 0.0006650034478442549 | -0.17530935706725967 | 0.006938243833717989 |
| asd | Right.VIIB | FIQ | 0.12304419521880301 | 0.05910890114936562 | -0.10328831659349932 | 0.11352272752007891 |
| asd | Right.VI | FIQ | 0.2417951108232532 | 0.0001764054140721864 | -0.16064883475361869 | 0.013477340607311103 |
| asd | Right.Crus.I | FIQ | 0.1933470738703676 | 0.0028573751411937083 | -0.13714346779501596 | 0.035238182892213026 |
| asd | Right.Crus.II | FIQ | 0.12722080422011786 | 0.05094239824492121 | -0.06042990920545032 | 0.35535211919829357 |
| asd | Right.VIIIB | FIQ | 0.08222806616532494 | 0.20816149978776088 | -0.12159151198388655 | 0.06219096123203937 |
| asd | Right.X | FIQ | 0.11203576027498571 | 0.0859063746210338 | -0.19971113657184889 | 0.002050306629481491 |
| asd | Left. Crus.I | PIQ | 0.1922013562287126 | 0.0030301123054460215 | -0.08074205702007513 | 0.21652702800834742 |
| asd | Left.Crus.II | PIQ | 0.18019390444333408 | 0.005498908783632934 | -0.11151636618308275 | 0.08737936221197985 |
| asd | Left.VIIB | PIQ | 0.0838704726564726 | 0.19918927056603322 | -0.09159411802673063 | 0.16074377016721458 |
| asd | Left.VI | PIQ | 0.22124128867626483 | 0.0006187792889412692 | -0.10470178302323721 | 0.10864096763644633 |
| asd | Left.VIIIA | PIQ | 0.08507190958834301 | 0.19280615157143913 | -0.02769916794319071 | 0.6720438598179244 |
| asd | Left..VI | PIQ | 0.23310760517662193 | 0.00030391561682042913 | -0.11395161930432764 | 0.08064462589275365 |
| asd | Left.X | PIQ | 0.12775564518123222 | 0.04996791081732876 | -0.12200677019670521 | 0.061296743080877646 |
| asd | Left.IV | PIQ | 0.20584592556502085 | 0.0014748109286242888 | -0.08828819547433282 | 0.1764557249461928 |
| asd | Left.VIIIA | PIQ | 0.06940815029719494 | 0.28828654386946495 | 0.010416248457744363 | 0.873533252947559 |
| asd | Left.IX | PIQ | 0.07573721412050435 | 0.24646234309042492 | -0.058935771993209436 | 0.3673942503728389 |
| asd | Left.I.III | PIQ | 0.10478564906340751 | 0.1083565839547207 | 0.007427307437832225 | 0.9096371730953259 |
| asd | Vermis.VIII | PIQ | 0.08177338958519548 | 0.21069601141263947 | -0.19311432313385848 | 0.0028917258500517156 |
| asd | Vermis.IX | PIQ | 0.12034312251965991 | 0.06494401707936226 | -0.2061161608676716 | 0.0014532480919195675 |
| asd | Vermis.VII | PIQ | 0.09002222289834592 | 0.16807826609582524 | -0.1539254035945195 | 0.01797122485857095 |
| asd | Vermis.VI | PIQ | 0.1741169701870691 | 0.007337087789308535 | -0.15495673874477767 | 0.017206825626778156 |
| asd | Vermis.X | PIQ | 0.11849267345405018 | 0.06920783583226794 | -0.07517296309145617 | 0.25000974662061876 |
| asd | Right.V | PIQ | 0.25526965164714527 | 7.289896856905132e-05 | -0.16374380129292082 | 0.011763859166727293 |
| asd | Right.IV | PIQ | 0.26421628915422224 | 3.944568857740468e-05 | -0.08058596061689202 | 0.21741953410434126 |

| | | | | | | |
|---|---|---|---|---|---|---|
| asd | Right.IX | PIQ | 0.04918777944004154 | 0.4520027800371702 | -0.09503003902193338 | 0.1455520335125134 |
| asd | Right.I.III | PIQ | 0.11624220102795128 | 0.07469839220401382 | -0.03410980634955444 | 0.6021060478507574 |
| asd | Right.Crus.II | PIQ | 0.19598125327716165 | 0.0024936065201133905 | -0.16963724531116844 | 0.009024308643072299 |
| asd | Right.VIIB | PIQ | 0.11204532363600411 | 0.08587944126313281 | -0.10978109322261775 | 0.09244788898015874 |
| asd | Right.VI | PIQ | 0.25316913156590487 | 8.393769654927195e-05 | -0.09394681975863854 | 0.15021869655378053 |
| asd | Right.Crus.I | PIQ | 0.18100323109048508 | 0.0052881878663578 | -0.12823958018770248 | 0.04909960445570013 |
| asd | Right.Crus.II | PIQ | 0.13370605556380402 | 0.04013570130078502 | -0.04355738118914074 | 0.5054721649608267 |
| asd | Right.VIIIB | PIQ | 0.11693972660572631 | 0.07296003753571274 | -0.054958018031843005 | 0.4006652512551213 |
| asd | Right.X | PIQ | 0.09536894060233711 | 0.14411488315146576 | -0.13797155049337176 | 0.034137215171336055 |
| asd | 73 | VIQ | 0.1531868535492718 | 0.018536623179652198 | -0.0030463825290251525 | 0.9628710544308737 |
| asd | Left. Crus.I | VIQ | 0.21368537529148202 | 0.0009548374435683365 | -0.06337734116721694 | 0.3323323663021268 |
| asd | Left.Crus.II | VIQ | 0.08163175911667353 | 0.21149001676948112 | -0.07376850337410096 | 0.25899284776813153 |
| asd | Left.VIIB | VIQ | 0.12149770356111958 | 0.06239444634056777 | -0.09414604415928024 | 0.14935200772948667 |
| asd | Left.VI | VIQ | 0.06481070660814015 | 0.3214916361193831 | -0.02121947902228732 | 0.7457218733657587 |
| asd | Left.VIIIA | VIQ | 0.15462461877202985 | 0.017449831474603973 | -0.1149514564351424 | 0.07800373516264589 |
| asd | Left..VI | VIQ | 0.09381762589916963 | 0.1507827638691891 | -0.027548736510603952 | 0.6737225290704618 |
| asd | Left.X | VIQ | 0.10982902645784345 | 0.09230479804913436 | -0.0817636573145678 | 0.21075050358270134 |
| asd | Left.IV | VIQ | 0.03565161900868111 | 0.585783881488283 | -0.04291033699402509 | 0.51182201775244 |
| asd | Left.VIIIA | VIQ | 0.06560250890206001 | 0.3156026430034776 | -0.02261407608765259 | 0.7296405877718992 |
| asd | Left.IX | VIQ | 0.08449876000040954 | 0.1958323605345737 | 0.0002471009403837571 | 0.996987289683569 |
| asd | Left.I.III | VIQ | 0.07448066536780185 | 0.2544103774483471 | -0.1331682612809915 | 0.040951560330861614 |
| asd | Vermis.VIII | VIQ | 0.03002148332386802 | 0.64633939866381 | -0.11846697577333995 | 0.06926861932064678 |
| asd | Vermis.IX | VIQ | 0.05476240052695399 | 0.40234654188745833 | -0.13530916456960443 | 0.03778455312070406 |
| asd | Vermis.VII | VIQ | 0.1325491235691472 | 0.04190806674420479 | -0.10562121842941967 | 0.10555509337881193 |
| asd | Vermis.VI | VIQ | 0.10385800161032849 | 0.11153491859791911 | 0.028914323084427724 | 0.6585440425852283 |
| asd | Vermis.X | VIQ | 0.17915894440951555 | 0.0057793342244139585 | -0.1825333795273725 | 0.004909493284757845 |
| asd | Right.V | VIQ | 0.13928996236084407 | 0.0324446528597524 | -0.1097395976079161 | 0.09257190511692703 |
| asd | Right.IV | VIQ | 0.035984906453599474 | 0.5822825875299282 | -0.09575846340167571 | 0.14247644762630537 |
| asd | Right.IX | VIQ | 0.17513392859064086 | 0.006995682086926318 | -0.04382125816175565 | 0.5028944565235074 |
| asd | Right.I.III | VIQ | 0.18724443394582865 | 0.0038917335045905634 | -0.0970066874995384 | 0.13732174147345852 |
| asd | Right.Crus.II | VIQ | 0.1066461835340375 | 0.10219692776411227 | -0.04303201258785403 | 0.5106247891739157 |
| asd | Right.VIIB | VIQ | 0.1799739676446216 | 0.0055574591198081095 | -0.12778966609030407 | 0.049906453179882404 |
| asd | Right.VI | VIQ | 0.1655038987004329 | 0.010877617488979054 | -0.07930179016910126 | 0.22486186142482936 |
| asd | Right.Crus.I | VIQ | 0.10935215485487852 | 0.09373625368653948 | -0.04000669682195477 | 0.5408181762857277 |
| asd | Right.Crus.II | VIQ | 0.05870144300296012 | 0.3693055028795993 | -0.11779316844334589 | 0.07087800448630158 |
| asd | Right.VIIIB | VIQ | 0.11679561892272948 | 0.07331645723235454 | -0.17333891098178453 | 0.007608239270008206 |

**Supplementary Table 8.** Association of extreme deviations and intelligence scores in ASD using task-based atlas

| Diagnosis | ROI | Test | Correlation of % positive dev | P value of % positive dev | Correlation of % negative dev | P value of % negative dev |
|---|---|---|---|---|---|---|
| asd | 10: Autobiographical recall/visual letter recognition/interference resolution | FIQ | 0.11895224578437975 | 0.06812812661958419 | -0.06057874112652337 | 0.3541662995081153 |
| asd | 3: Saccades/visual working memory/visual letter recognition | FIQ | 0.18403747204819168 | 0.004561182807096344 | -0.21508215869815914 | 0.0008822264769072235 |
| asd | 4: Action Observation/divided attention/motor planning | FIQ | 0.1499781062910333 | 0.021176338441102546 | -0.08331503611835965 | 0.20219156741247152 |

| | | | | | | |
|---|---|---|---|---|---|---|
| asd | 5: Divided attention/active maintenance/mental arithmetic | FIQ | 0.15949715808756934 | 0.014168788359368438 | -0.11594392072844818 | 0.07545194856170635 |
| asd | 6: Divided attention/verbal fluency/active maintenance | FIQ | 0.1795048517197964 | 0.005684215801323208 | -0.13027520962916042 | 0.04558374276179747 |
| asd | 9: Verbal Fluency/word comprehension/mental arithmetic | FIQ | 0.1498139985234233 | 0.02131967767682188 | -0.12316510085257527 | 0.058858164482092024 |
| asd | 1: Left-hand presses/ motor planning/ interference resolution | FIQ | 0.17399878245994438 | 0.00737771293757722 | -0.12468154802667523 | 0.055787179521880406 |
| asd | 2: Right-hand presses/ motor planning/ divided attention | FIQ | 0.22055170154311726 | 0.0006441586335142442 | -0.19407844127536392 | 0.002751830905785027 |
| asd | 7: Narrative/ emotion processing/ language processing | FIQ | 0.17574932464650056 | 0.006796038251038484 | -0.09400979168467934 | 0.14994433796884443 |
| asd | 8: Word comprehension/ language processing/ narrative | FIQ | 0.1842795252466251 | 0.004507255377922146 | -0.17744101434744416 | 0.0062731862074951305 |
| asd | 10: Autobiographical recall/visual letter recognition/interference resolution | PIQ | 0.11345746268245953 | 0.08197624006101308 | -0.058789122744481094 | 0.3685896444723047 |
| asd | 3: Saccades/visual working memory/visual letter recognition | PIQ | 0.19740473040354253 | 0.00231505368137928 | -0.19290616732699933 | 0.002922763481803223 |
| asd | 4: Action Observation/divided attention/motor planning | PIQ | 0.16017804500899288 | 0.013756331672693098 | -0.030150431212192127 | 0.6449240058308587 |
| asd | 5: Divided attention/active maintenance/mental arithmetic | PIQ | 0.14764497131189008 | 0.02329453445967468 | -0.10299273635512993 | 0.11456496753809084 |
| asd | 6: Divided attention/verbal fluency/active maintenance | PIQ | 0.18798491712280327 | 0.003750355668343521 | -0.12332391586996361 | 0.05853014434952116 |
| asd | 9: Verbal Fluency/word comprehension/mental arithmetic | PIQ | 0.14837390446135476 | 0.02261393862800952 | -0.1371003277592686 | 0.03529635504401125 |
| asd | 1: Left-hand presses/ motor planning/ interference resolution | PIQ | 0.18257704396851435 | 0.004899053892231517 | -0.07118616847070348 | 0.2760839641095939 |
| asd | 2: Right-hand presses/ motor planning/ divided attention | PIQ | 0.2338494726683431 | 0.00029034728021905064 | -0.14110035655689748 | 0.0302372532389639 |
| asd | 7: Narrative/ emotion processing/ language processing | PIQ | 0.17973943820636862 | 0.0056205094583171595 | -0.09972120729591533 | 0.12660629191463893 |
| asd | 8: Word comprehension/ language processing/ narrative | PIQ | 0.1741838207505504 | 0.007314197373441939 | -0.17285059312553983 | 0.007782941464632897 |
| asd | 10: Autobiographical recall/visual letter recognition/interference resolution | VIQ | 0.12794650240721 | 0.04962394732436631 | -0.03165195489922811 | 0.6285377198283666 |
| asd | 3: Saccades/visual working memory/visual letter recognition | VIQ | 0.14412961227801005 | 0.02683033540868835 | -0.13479243607719646 | 0.03852934442150161 |
| asd | 4: Action Observation/divided attention/motor planning | VIQ | 0.12561737920381125 | 0.05395905945786877 | -0.07142348785567905 | 0.274482112627284 |
| asd | 5: Divided attention/active maintenance/mental arithmetic | VIQ | 0.1394623206369305 | 0.03222875173452967 | -0.06220595704638704 | 0.341363861161196 |
| asd | 6: Divided attention/verbal fluency/active maintenance | VIQ | 0.13198828483123592 | 0.04279065648167175 | -0.05176890586324462 | 0.42859548861066676 |
| asd | 9: Verbal Fluency/word comprehension/mental arithmetic | VIQ | 0.13225824528225266 | 0.042363890827744236 | -0.06950569609096802 | 0.28760786102668234 |
| asd | 1: Left-hand presses/ motor planning/ interference resolution | VIQ | 0.13537628529364967 | 0.03768870700183165 | -0.09796911555074342 | 0.13344578600411422 |

| | | | | | | |
|---|---|---|---|---|---|---|
| asd | 2: Right-hand presses/ motor planning/ divided attention | VIQ | 0.1654183156803436 | 0.010919306183696789 | -0.14390574863926606 | 0.02707031922693994 |
| asd | 7: Narrative/ emotion processing/ language processing | VIQ | 0.1409922101605215 | 0.030365417521855013 | -0.030272414487217025 | 0.6435862354291426 |
| asd | 8: Word comprehension/ language processing/ narrative | VIQ | 0.16241536960545258 | 0.012474287713275037 | -0.1029309818154008 | 0.11478366097109313 |

**Supplementary Table 9**. Association of extreme deviations and intelligence scores in ASD using resting-state atlas

| Diagnosis | ROI | Test | Correlation of % positive deviation | P value of % positive deviation | Correlation of % negative deviation | P value of % negative deviation |
|---|---|---|---|---|---|---|
| asd | 13: Control C | FIQ | 0.21243785789785558 | 0.0010242974872942067 | -0.14382193543625732 | 0.027160640164402592 |
| asd | 16: Default C | FIQ | 0.17418977402390798 | 0.007312161993314162 | -0.11025587187549492 | 0.09103835397871875 |
| asd | 8: Salience/ Ventral Attention B | FIQ | 0.17386973464186087 | 0.007422299926288363 | -0.13298894589362395 | 0.04122667529517777 |
| asd | 11: Control A | FIQ | 0.14146511373156131 | 0.02980837971841762 | -0.07667824373001043 | 0.2406244171977096 |
| asd | 7: Salience/Ventral Attention A | FIQ | 0.17089206022405534 | 0.008520053417030164 | -0.1105938589834288 | 0.09004543144560917 |
| asd | 12 Control B | FIQ | 0.13998946479937632 | 0.03157601121668286 | -0.1185608485054588 | 0.06904679025423076 |
| asd | 17: Temporal Parietal | FIQ | 0.20933836759060553 | 0.0012174612053769277 | -0.13423439313950045 | 0.03934757024509968 |
| asd | 6: Dorsal Attention B | FIQ | 0.12375007855646397 | 0.057657387012651204 | -0.09735501233728554 | 0.13590910342897114 |
| asd | 9: Limbic B | FIQ | 0.12338677641705618 | 0.05840072815156439 | -0.1904596894105129 | 0.003310873129308823 |
| asd | 3: Somatomotor A | FIQ | 0.13963116225024627 | 0.03201844291513889 | -0.15803158386560628 | 0.015093532082261489 |
| asd | 15: Default B | FIQ | 0.1557672779460081 | 0.01662611489745426 | -0.1682371284713098 | 0.009618053557383885 |
| asd | 4: Somatomotor B | FIQ | 0.22425068172167892 | 0.0005184765962852928 | -0.1367225688296502 | 0.035809230972442804 |
| asd | 14: Default A | FIQ | 0.08692061048188147 | 0.18327792557483685 | -0.13206280719944624 | 0.0426724888475242 |
| asd | 5: Dorsal Attention A | FIQ | 0.14409452678902923 | 0.02686782631888584 | -0.1412602741313887 | 0.030048581444744205 |
| asd | 2: Visual B | FIQ | 0.14476937181754584 | 0.0261545441241114 | -0.1531559202411191 | 0.01856063796243722 |
| asd | 10: Limbic A | FIQ | 0.004858884246307866 | 0.9408132843007142 | -0.04101097485341399 | 0.5306977550921939 |
| asd | 1: Visual A | FIQ | 0.05596070752246759 | 0.39211339712574944 | -0.025201384661402107 | 0.700121396858381 |
| asd | 13: Control C | PIQ | 0.20936601843514024 | 0.001215599319642519 | -0.12672205208684587 | 0.05186533043856839 |
| asd | 16: Default C | PIQ | 0.15996970653778264 | 0.013881402439878333 | -0.11572518096631945 | 0.0760084677596895 |
| asd | 8: Salience/ Ventral Attention B | PIQ | 0.15683010917513926 | 0.015890607230708658 | -0.10054026666066067 | 0.12350348831569456 |
| asd | 11: Control A | PIQ | 0.14107647476460233 | 0.030265515746945887 | -0.07056318543229743 | 0.2803190496857079 |
| asd | 7: Salience/Ventral Attention A | PIQ | 0.17865631521375364 | 0.005920090306697551 | -0.0659754362284742 | 0.31285355414515226 |
| asd | 12 Control B | PIQ | 0.13924911755619454 | 0.032495996329943884 | -0.10663630307169447 | 0.10222889170627955 |
| asd | 17: Temporal Parietal | PIQ | 0.20714123801033568 | 0.0013740528484109784 | -0.124932022316219 | 0.05529293174696321 |
| asd | 6: Dorsal Attention B | PIQ | 0.13341774937091141 | 0.04057135535164941 | -0.06213372970780528 | 0.34192580413918705 |
| asd | 9: Limbic B | PIQ | 0.13544848776065266 | 0.03758583385803283 | -0.13994206447359522 | 0.031634240436887 |
| asd | 3: Somatomotor A | PIQ | 0.18017247905938727 | 0.005504588110272361 | -0.05595401378794155 | 0.3921701201772242 |
| asd | 15: Default B | PIQ | 0.17578301831093335 | 0.006785255882064922 | -0.15127826296754843 | 0.020069912830392733 |
| asd | 4: Somatomotor B | PIQ | 0.250461381244642 | 0.00010049027333792826 | -0.07885265886525492 | 0.22750697022608052 |
| asd | 14: Default A | PIQ | 0.12218798491616435 | 0.060909844183766125 | -0.11542945222819027 | 0.07676614781534039 |
| asd | 5: Dorsal Attention A | PIQ | 0.1602175201154994 | 0.013732745529710145 | -0.13774118714553973 | 0.034440519196913696 |
| asd | 2: Visual B | PIQ | 0.14721514963623866 | 0.02370407729679787 | -0.12260349899672042 | 0.06003029421870536 |
| asd | 10: Limbic A | PIQ | 0.03626000241011993 | 0.57939994996658 | -0.0011008146611909766 | 0.9865792168890208 |
| asd | 1: Visual A | PIQ | 0.05130548057820378 | 0.4327457630975632 | -0.034640520075717 | 0.5964647527927149 |
| asd | 13: Control C | VIQ | 0.18309463283867825 | 0.004776818713101596 | -0.095784341444159 | 0.14236810318404558 |
| asd | 16: Default C | VIQ | 0.16326130470505762 | 0.012017658802246735 | -0.05243794610824057 | 0.422644561993489 |

| | | | | | | |
|---|---|---|---|---|---|---|
| asd | 8: Salience/ Ventral Attention B | VIQ | 0.1446477821407997 | 0.026281844941067646 | -0.08747986395019626 | 0.18046498659043697 |
| asd | 11: Control A | VIQ | 0.12660949503456936 | 0.052075526404004535 | -0.014837065219198284 | 0.8206309088463328 |
| asd | 7: Salience/Ventral Attention A | VIQ | 0.1319729439962302 | 0.0428150160179383 | -0.08895040190249816 | 0.17322069470298498 |
| asd | 12 Control B | VIQ | 0.12252519422190584 | 0.06019524290655343 | -0.061873053270905747 | 0.3439588055104126 |
| asd | 17: Temporal Parietal | VIQ | 0.1634981325004354 | 0.011892487812675642 | -0.07611178510038712 | 0.244126887186978 |
| asd | 6: Dorsal Attention B | VIQ | 0.10778647833483616 | 0.09856077645166204 | -0.0662160669082198 | 0.3110880514090448 |
| asd | 9: Limbic B | VIQ | 0.10401189624244662 | 0.1110026410580695 | -0.13800884245229605 | 0.03408832929473468 |
| asd | 3: Somatomotor A | VIQ | 0.08961098434694531 | 0.17003769094509338 | -0.1515236788281825 | 0.019866786649224845 |
| asd | 15: Default B | VIQ | 0.13604941680992882 | 0.036738812359438426 | -0.1030946669584479 | 0.11420471006079343 |
| asd | 4: Somatomotor B | VIQ | 0.15946576101863208 | 0.014188066336133343 | -0.13628104654513357 | 0.03641666737083378 |
| asd | 14: Default A | VIQ | 0.05422024448351307 | 0.40702817423322535 | -0.08409906532672712 | 0.1979631083318005 |
| asd | 5: Dorsal Attention A | VIQ | 0.0861645790707036 | 0.1871317528712264 | -0.08959776009703616 | 0.1701009815313175 |
| asd | 2: Visual B | VIQ | 0.1340739219519136 | 0.03958555578558694 | -0.11900503862276264 | 0.06800498235515248 |
| asd | 10: Limbic A | VIQ | 0.01742253940205677 | 0.7900472669569881 | -0.021862477561853316 | 0.7382932070083068 |
| asd | 1: Visual A | VIQ | 0.05829046693897756 | 0.3726723388793839 | 0.01276311533370527 | 0.8453635179945831 |

**Supplementary Table 10**. Association of extreme deviations and ADOS scores in ASD using anatomical atlas

| Diagnosis | ROI | Test | Correlation of % positive deviation | P value of % positive deviation | Correlation of % negative deviation | P value of % negative deviation |
|---|---|---|---|---|---|---|
| asd | Left. Crus.I | ADOS_COMM | 0.13671858903872278 | 0.03581466769414873 | -0.06099275637065445 | 0.3508807133501377 |
| asd | Left.Crus.II | ADOS_COMM | 0.11927948294230671 | 0.06736774220680193 | -0.12754885303679245 | 0.05034283162523417 |
| asd | Left.VIIB | ADOS_COMM | 0.013483706283126989 | 0.8367517238486035 | -0.11155819357851299 | 0.08725999611791797 |
| asd | Left.VI | ADOS_COMM | 0.1539857919961727 | 0.017925665338598046 | -0.10045028344946356 | 0.12384145567446778 |
| asd | Left.VIIIA | ADOS_COMM | 0.032704738340586514 | 0.6171554569639535 | -0.10811260537469196 | 0.0975399255577135 |
| asd | Left..VI | ADOS_COMM | 0.1491018882771471 | 0.02195143523001151 | -0.1281123078859442 | 0.04932673594917621 |
| asd | Left.X | ADOS_COMM | 0.1745491061338851 | 0.0071902401255103075 | -0.027543689337823588 | 0.6737788786948864 |
| asd | Left.IV | ADOS_COMM | 0.2110901534967471 | 0.0011045349007155853 | -0.03680058840318714 | 0.5737548026637136 |
| asd | Left.VIIIA | ADOS_COMM | 0.10645868399957883 | 0.10280485039572564 | -0.017253217246706812 | 0.792040929384941 |
| asd | Left.IX | ADOS_COMM | 0.15856459022534655 | 0.014751279932590825 | -0.036929454405710764 | 0.5724129330320019 |
| asd | Left.I.III | ADOS_COMM | 0.15274664954875092 | 0.01888092693816362 | -0.18552347315634235 | 0.0042390632315967075 |
| asd | Vermis.VIII | ADOS_COMM | 0.15365496210307814 | 0.018176495587101743 | -0.0583279206619337 | 0.37236472684646704 |
| asd | Vermis.IX | ADOS_COMM | 0.15880790077060725 | 0.014597318578311593 | -0.09095328474289731 | 0.16370435738956818 |
| asd | Vermis.VII | ADOS_COMM | 0.11591101689798428 | 0.07553545093083794 | -0.05991035118273989 | 0.35951118284985584 |
| asd | Vermis.VI | ADOS_COMM | 0.13409616545117994 | 0.03955249544389646 | -0.15890214913451922 | 0.014538059658728232 |
| asd | Vermis.X | ADOS_COMM | 0.1535176704451786 | 0.01828148126064723 | -0.10417221039661512 | 0.11045028122896186 |
| asd | Right.V | ADOS_COMM | 0.10065763388439515 | 0.12306374514927652 | -0.1036878136991332 | 0.11212587778355605 |
| asd | Right.IV | ADOS_COMM | 0.18765462069246747 | 0.0038128348225448566 | -0.00821594969471543 | 0.900089899946831 |
| asd | Right.IX | ADOS_COMM | 0.16863851670993135 | 0.009444396905083093 | -0.026058650087637938 | 0.6904365554386382 |
| asd | Right.I.III | ADOS_COMM | 0.15917476179989556 | 0.014367838485344701 | 0.007211590482294881 | 0.9122509531677182 |
| asd | Right.Crus.II | ADOS_COMM | 0.09073578561550905 | 0.16471840725028786 | -0.1743664440157841 | 0.007251989200435428 |
| asd | Right.VIIB | ADOS_COMM | 0.08825245730514232 | 0.17663157806545893 | -0.18736245659071843 | 0.0038688824206862247 |
| asd | Right.VI | ADOS_COMM | 0.11127730852827128 | 0.08806409388416389 | -0.11204896098760353 | 0.08586919914513935 |
| asd | Right.Crus.I | ADOS_COMM | 0.15348190995276004 | 0.018308913417243892 | -0.10402332982308975 | 0.11096317522258442 |
| asd | Right.Crus.II | ADOS_COMM | 0.07659207988951654 | 0.241154896593278 | -0.15295326825911784 | 0.01871863500411129 |
| asd | Right.VIIIB | ADOS_COMM | 0.16240859321356432 | 0.012478006356078838 | -0.05936853024196028 | 0.36388068469860735 |
| asd | Right.X | ADOS_COMM | 0.2219208922177269 | 0.0005946746544330556 | 0.059796326595620224 | 0.3604279997683364 |

| | | | | | | |
|---|---|---|---|---|---|---|
| asd | Left. Crus.I | ADOS_SOCIAL | 0.10797790470940909 | 0.09796054642868327 | -0.12503137474723766 | 0.055097891829016206 |
| asd | Left.Crus.II | ADOS_SOCIAL | 0.11128680286974507 | 0.0880368175531844 | -0.19087279233597185 | 0.0032422294178624804 |
| asd | Left.VIIB | ADOS_SOCIAL | 0.04832715951945612 | 0.459964151130582 | -0.15918036125112933 | 0.01436436053418198 |
| asd | Left.VI | ADOS_SOCIAL | 0.07840032256815821 | 0.23019314984343708 | -0.10413676945486032 | 0.11057220632544391 |
| asd | Left.VIIIA | ADOS_SOCIAL | 0.052332611927083404 | 0.42357827765265954 | -0.16948765865442797 | 0.00908615344300526 |
| asd | Left..VI | ADOS_SOCIAL | 0.13233763286623162 | 0.04223907410230619 | -0.12426185525867843 | 0.056623530873204084 |
| asd | Left.X | ADOS_SOCIAL | 0.16674592339037006 | 0.010288237339195079 | -0.1379429191485206 | 0.03417478827455196 |
| asd | Left.IV | ADOS_SOCIAL | 0.17645244447852912 | 0.006574174625886857 | 0.018698378207958465 | 0.7750697972788744 |
| asd | Left.VIIIA | ADOS_SOCIAL | 0.061797720378931574 | 0.3445477478012221 | -0.05997529668429278 | 0.3589896385593203 |
| asd | Left.IX | ADOS_SOCIAL | 0.14567891178959372 | 0.02521897059521929 | -0.033077286991889415 | 0.6131492164158068 |
| asd | Left.I.III | ADOS_SOCIAL | 0.05751260497973809 | 0.37909633092847683 | -0.09268732954934261 | 0.15578629008779962 |
| asd | Vermis.VIII | ADOS_SOCIAL | 0.10114278030066269 | 0.12125891259952891 | -0.10793902167867316 | 0.09808223132295746 |
| asd | Vermis.IX | ADOS_SOCIAL | 0.15270899351053246 | 0.01891063581929309 | -0.039706860235585596 | 0.54385819238514 |
| asd | Vermis.VII | ADOS_SOCIAL | 0.12691659512313602 | 0.05150369183179467 | -0.1039123024720435 | 0.1113468794383402 |
| asd | Vermis.VI | ADOS_SOCIAL | 0.0997905309813453 | 0.1263413627502769 | -0.11722274000592862 | 0.072264164207331 |
| asd | Vermis.X | ADOS_SOCIAL | 0.07772204501804107 | 0.23426293388084732 | -0.06104424619383072 | 0.3504734414165467 |
| asd | Right.V | ADOS_SOCIAL | 0.08319137561176071 | 0.20286442550285116 | -0.10901736833396637 | 0.09475173168790776 |
| asd | Right.IV | ADOS_SOCIAL | 0.10085915852242198 | 0.12231152218346623 | 0.027675699524850675 | 0.6723056379812313 |
| asd | Right.IX | ADOS_SOCIAL | 0.1616303522784774 | 0.012911619297547338 | -0.010128393883201388 | 0.8770001471176873 |
| asd | Right.I.III | ADOS_SOCIAL | 0.022389119987884978 | 0.7322268202519548 | -0.041901626633305926 | 0.5218029941030482 |
| asd | Right.Crus.II | ADOS_SOCIAL | 0.11315088170496092 | 0.08281125873384526 | -0.1404484439847099 | 0.031016884174026083 |
| asd | Right.VIIB | ADOS_SOCIAL | 0.12849239945280466 | 0.04865100685819769 | -0.21627777053771377 | 0.0008241505069863097 |
| asd | Right.VI | ADOS_SOCIAL | 0.06308629061185327 | 0.33456194879369505 | -0.12383490125245002 | 0.057484963943317466 |
| asd | Right.Crus.I | ADOS_SOCIAL | 0.07471857669475485 | 0.25289208127977264 | -0.13895675340563474 | 0.032865532015721476 |
| asd | Right.Crus.II | ADOS_SOCIAL | 0.07378377027057223 | 0.2588940194063455 | -0.1747484877142458 | 0.007123374635927767 |
| asd | Right.VIIIB | ADOS_SOCIAL | 0.13370186546326865 | 0.040142004537266296 | -0.06862993116448969 | 0.2937394740233279 |
| asd | Right.X | ADOS_SOCIAL | 0.16303744083568283 | 0.012137043268011653 | -0.021871935659955825 | 0.7381841151118311 |
| asd | Left. Crus.I | ADOS_STEREO_BEHAV | 0.034542565591863854 | 0.5975041725327037 | 0.15988441316292273 | 0.013932893266563613 |
| asd | Left.Crus.II | ADOS_STEREO_BEHAV | 0.09854225653371997 | 0.13117788310067172 | 0.16680145780732827 | 0.010262557035890824 |
| asd | Left.VIIB | ADOS_STEREO_BEHAV | 0.16608442266019088 | 0.010598527736687277 | 0.192201165243548 | 0.003030141872113564 |
| asd | Left.VI | ADOS_STEREO_BEHAV | 0.10508070186560989 | 0.1073607308486509 | 0.146019108693468 | 0.024876507904874078 |
| asd | Left.VIIIA | ADOS_STEREO_BEHAV | 0.13101233446657318 | 0.04436372688904364 | 0.11153316195397317 | 0.08733141509392488 |
| asd | Left..VI | ADOS_STEREO_BEHAV | 0.1368484044031503 | 0.03563768950639349 | 0.026238562815785394 | 0.6884103291435513 |
| asd | Left.X | ADOS_STEREO_BEHAV | 0.08282876193138423 | 0.2048468088089934 | -0.000522726102685093 | 0.9936268579173754 |
| asd | Left.IV | ADOS_STEREO_BEHAV | 0.1501863736298626 | 0.02099562979550516 | 0.004301719393309805 | 0.9475898478355557 |
| asd | Left.VIIIA | ADOS_STEREO_BEHAV | 0.19559024798017982 | 0.002544796867045744 | 0.050305246201819184 | 0.44178185446380613 |
| asd | Left.IX | ADOS_STEREO_BEHAV | 0.183430476554547 | 0.004698979055718794 | 0.013961837430003203 | 0.8310482127689148 |
| asd | Left.I.III | ADOS_STEREO_BEHAV | 0.05159880536410336 | 0.4301161702152746 | -0.10025004772416427 | 0.1245960928999507 |
| asd | Vermis.VIII | ADOS_STEREO_BEHAV | 0.17319175829083058 | 0.007660513749361426 | 0.013532997133617112 | 0.8361633455017644 |
| asd | Vermis.IX | ADOS_STEREO_BEHAV | 0.16565899649037205 | 0.0108024252171873 | 0.03542799503683998 | 0.5881385536550869 |
| asd | Vermis.VII | ADOS_STEREO_BEHAV | 0.08969834063450821 | 0.1696200486914342 | 0.04502078860053428 | 0.49126428266461797 |
| asd | Vermis.VI | ADOS_STEREO_BEHAV | 0.12294423483004774 | 0.05931686574013487 | -0.0553516689617602 | 0.3972946429182538 |
| asd | Vermis.X | ADOS_STEREO_BEHAV | 0.018270629595277803 | 0.7800823148658582 | 0.057643191007346335 | 0.37801318033949827 |
| asd | Right.V | ADOS_STEREO_BEHAV | 0.08420521414530902 | 0.19739559914330715 | -0.025979779893933845 | 0.6913255095555119 |
| asd | Right.IV | ADOS_STEREO_BEHAV | 0.08968620161997509 | 0.1696780384246022 | -0.025427756774266234 | 0.6975591988794383 |
| asd | Right.IX | ADOS_STEREO_BEHAV | 0.15943265565811743 | 0.014208418115022025 | -0.004622817791179208 | 0.9436839707020178 |
| asd | Right.I.III | ADOS_STEREO_BEHAV | 0.06708672519789956 | 0.30475471042083363 | 0.013841699104668039 | 0.832480494774952 |

| | | | | | | |
|---|---|---|---|---|---|---|
| asd | Right.Crus.II | ADOS_STEREO_BEHAV | 0.13960925217065673 | 0.03204566790783759 | 0.1653320703043571 | 0.010961460176935006 |
| asd | Right.VIIB | ADOS_STEREO_BEHAV | 0.22743738981511383 | 0.000428832091461221 | 0.13999625007901284 | 0.031567683276137515 |
| asd | Right.VI | ADOS_STEREO_BEHAV | 0.14203951169633663 | 0.02914355218874837 | 0.049730176117710134 | 0.4470252380345747 |
| asd | Right.Crus.I | ADOS_STEREO_BEHAV | 0.0835922265084639 | 0.20068920136409493 | 0.14037734845649102 | 0.031102935912529477 |
| asd | Right.Crus.II | ADOS_STEREO_BEHAV | 0.17192278432158872 | 0.008124742317733002 | 0.07319885798837923 | 0.26269894854694115 |
| asd | Right.VIIIB | ADOS_STEREO_BEHAV | 0.1693790049268404 | 0.009131310438205807 | 0.09768247351160485 | 0.13459127086181377 |
| asd | Right.X | ADOS_STEREO_BEHAV | 0.10259110349205927 | 0.11599312934114983 | 0.06038349292009287 | 0.35572244997066926 |

**Supplementary Table 11**. Association of extreme deviations and ADOS scores in ASD using task-based atlas

| Diagnosis | ROI | Test | Correlation of % positive deviation | P value of % positive deviation | Correlation of % negative deviation | P value of % negative deviation |
|---|---|---|---|---|---|---|
| asd | 10: Autobiographical recall/visual letter recognition/interference resolution | ADOS_COMM | 0.07396830628788242 | 0.2577015017090951 | -0.10335986363134692 | 0.11327156569790214 |
| asd | 3: Saccades/visual working memory/visual letter recognition | ADOS_COMM | 0.19520622229661086 | 0.002596001125426236 | 0.010651052650034612 | 0.8707070752604495 |
| asd | 4: Action Observation/divided attention/motor planning | ADOS_COMM | 0.11104670639244191 | 0.08872867347927878 | -0.06388018904816008 | 0.32850284452500134 |
| asd | 5: Divided attention/active maintenance/mental arithmetic | ADOS_COMM | 0.10367942563362394 | 0.11215506775916222 | -0.14340933973867412 | 0.027609048715940755 |
| asd | 6: Divided attention/verbal fluency/active maintenance | ADOS_COMM | 0.12853668138927357 | 0.048572786830664044 | -0.12527344763934747 | 0.05462506040102423 |
| asd | 9: Verbal Fluency/word comprehension/mental arithmetic | ADOS_COMM | 0.10569214479465472 | 0.10531994560414645 | -0.16193183799123226 | 0.012742093205764787 |
| asd | 1: Left-hand presses/ motor planning/ interference resolution | ADOS_COMM | 0.15851663412241093 | 0.014781792653635767 | -0.10244624814524865 | 0.11651161666700742 |
| asd | 2: Right-hand presses/ motor planning/ divided attention | ADOS_COMM | 0.15729608212366 | 0.015577210854554031 | -0.07033055670478434 | 0.2819116622540185 |
| asd | 7: Narrative/ emotion processing/ language processing | ADOS_COMM | 0.1482905198718958 | 0.022690913880560496 | -0.14770724480307879 | 0.023235708141747642 |
| asd | 8: Word comprehension/ language processing/ narrative | ADOS_COMM | 0.13430832182228264 | 0.03923833764974743 | -0.14975335521195618 | 0.021372858366067737 |
| asd | 10: Autobiographical recall/visual letter recognition/interference resolution | ADOS_SOCIAL | 0.10161314925241682 | 0.11952876236881746 | -0.14484228760533352 | 0.026078457714653357 |
| asd | 3: Saccades/visual working memory/visual letter recognition | ADOS_SOCIAL | 0.1811257951121952 | 0.005256916160254557 | -0.016381319232245735 | 0.8023281182601807 |
| asd | 4: Action Observation/divided attention/motor planning | ADOS_SOCIAL | 0.10992173694021844 | 0.09202853901240325 | -0.10671867895267455 | 0.10196264198264034 |
| asd | 5: Divided attention/active maintenance/mental arithmetic | ADOS_SOCIAL | 0.09090591090832568 | 0.16392483003386546 | -0.13732938687588536 | 0.034988409760012454 |
| asd | 6: Divided attention/verbal fluency/active maintenance | ADOS_SOCIAL | 0.12311950686006928 | 0.05895261502754199 | -0.1316216710087062 | 0.043375993472408565 |
| asd | 9: Verbal Fluency/word comprehension/mental arithmetic | ADOS_SOCIAL | 0.10736168029495462 | 0.09990317997517457 | -0.144772583471747 | 0.026151188807337414 |
| asd | 1: Left-hand presses/ motor planning/ interference resolution | ADOS_SOCIAL | 0.1370321134807552 | 0.03538850470969909 | -0.10237377097498825 | 0.11677171556554605 |

| | | | | | | |
|---|---|---|---|---|---|---|
| asd | 2: Right-hand presses/ motor planning/ divided attention | ADOS_SOCIAL | 0.13335206706167474 | 0.040671161793231936 | -0.0828176638103397 | 0.2049077013987111 |
| asd | 7: Narrative/ emotion processing/ language processing | ADOS_SOCIAL | 0.11282476410177512 | 0.08370699163605205 | -0.12163050947041014 | 0.06210653027410852 |
| asd | 8: Word comprehension/ language processing/ narrative | ADOS_SOCIAL | 0.07420133213707207 | 0.25620104663614574 | -0.11459450728351346 | 0.078938405589798 |
| asd | 10: Autobiographical recall/visual letter recognition/interference resolution | ADOS_STEREO_BEHAV | 0.15041849266513727 | 0.0207958002632068 | 0.1637634041670396 | 0.011753648399986468 |
| asd | 3: Saccades/visual working memory/visual letter recognition | ADOS_STEREO_BEHAV | 0.15693869271118022 | 0.01581708974589929 | 0.10544444400050405 | 0.10614296884124733 |
| asd | 4: Action Observation/divided attention/motor planning | ADOS_STEREO_BEHAV | 0.1911138435531016 | 0.003202772643372687 | 0.1805716563793896 | 0.005399635625316748 |
| asd | 5: Divided attention/active maintenance/mental arithmetic | ADOS_STEREO_BEHAV | 0.058164316095170324 | 0.373709581518873 | 0.14907093578309155 | 0.021979258286119224 |
| asd | 6: Divided attention/verbal fluency/active maintenance | ADOS_STEREO_BEHAV | 0.15300716350524848 | 0.018676502037611453 | 0.14252614514791836 | 0.028590266906020765 |
| asd | 9: Verbal Fluency/word comprehension/mental arithmetic | ADOS_STEREO_BEHAV | 0.14982958483638784 | 0.021306027889402782 | 0.16939844620652278 | 0.00912321595998434 |
| asd | 1: Left-hand presses/ motor planning/ interference resolution | ADOS_STEREO_BEHAV | 0.13241707726416874 | 0.04211447764876523 | 0.07420437921900484 | 0.256181466472992 |
| asd | 2: Right-hand presses/ motor planning/ divided attention | ADOS_STEREO_BEHAV | 0.1032147473889335 | 0.11378144232214525 | 0.04034582606839429 | 0.5373899755420666 |
| asd | 7: Narrative/ emotion processing/ language processing | ADOS_STEREO_BEHAV | 0.0830543439705495 | 0.20361192940246509 | 0.12676229932269278 | 0.05179034165428215 |
| asd | 8: Word comprehension/ language processing/ narrative | ADOS_STEREO_BEHAV | 0.13942412821545036 | 0.03227648691603674 | 0.11525931630870395 | 0.07720481406367782 |

**Supplementary Table 12**. Association of extreme deviations and ADOS scores in ASD using resting-state atlas

| Diagnosis | ROI | Test | Correlation of % positive deviation | P value of % positive deviation | Correlation of % negative deviation | P value of % negative deviation |
|---|---|---|---|---|---|---|
| asd | 13: Control C | ADOS_COMM | 0.12035142294194066 | 0.06492538770339208 | -0.17076379256857982 | 0.008570424273780375 |
| asd | 16: Default C | ADOS_COMM | 0.14330201178008795 | 0.027726727111322926 | -0.09696333958629205 | 0.13749832406627746 |
| asd | 8: Salience/ Ventral Attention B | ADOS_COMM | 0.12989463292128756 | 0.04622455228159063 | -0.02399618334092265 | 0.7138190223766288 |
| asd | 11: Control A | ADOS_COMM | 0.21875559156835836 | 0.0007148463186583321 | 0.0025715238384741534 | 0.9686553430011122 |
| asd | 7: Salience/Ventral Attention A | ADOS_COMM | 0.1289875348027983 | 0.04778235130985892 | -0.08658629485684817 | 0.1849748123742711 |
| asd | 12 Control B | ADOS_COMM | 0.07723353172578952 | 0.2372253022973874 | -0.14628929010304267 | 0.024607385194284383 |
| asd | 17: Temporal Parietal | ADOS_COMM | 0.11948647488836264 | 0.06689035838462506 | -0.1251182262677007 | 0.05492785972048443 |
| asd | 6: Dorsal Attention B | ADOS_COMM | 0.08489444355389487 | 0.1937394939642686 | -0.0833790884128121 | 0.20184368387478988 |
| asd | 9: Limbic B | ADOS_COMM | 0.16247000343443818 | 0.012444342215843972 | -0.06509045039410395 | 0.31940296182127476 |
| asd | 3: Somatomotor A | ADOS_COMM | 0.1307541373670846 | 0.04478791629908339 | -0.05375961233496663 | 0.4110310820744062 |
| asd | 15: Default B | ADOS_COMM | 0.22552260170254002 | 0.000480796310411915 | -0.07657081507303476 | 0.24128594210669826 |
| asd | 4: Somatomotor B | ADOS_COMM | 0.15309481645306441 | 0.018608154811634938 | -0.08853101907763383 | 0.17526431996969707 |
| asd | 14: Default A | ADOS_COMM | 0.2277386055649829 | 0.0004211486215482899 | -0.17396799162022483 | 0.007388329663864708 |
| asd | 5: Dorsal Attention A | ADOS_COMM | 0.17252254657277202 | 0.007902301053270491 | -0.0712250074579977 | 0.275821377236029 |
| asd | 2: Visual B | ADOS_COMM | 0.10107858294433913 | 0.12149655056626658 | -0.10318099681277065 | 0.11390028446761444 |
| asd | 10: Limbic A | ADOS_COMM | 0.0728678605546579 | 0.2648690443603762 | 0.02649714611196395 | 0.6855019840114555 |

| | | | | | | |
|---|---|---|---|---|---|---|
| asd | 1: Visual A | ADOS_COMM | -0.05012440958623565 | 0.4434269074915428 | -0.07266528682615918 | 0.26620320271488546 |
| asd | 13: Control C | ADOS_SOCIAL | 0.09919933578358209 | 0.1286145040102676 | -0.15321350449497637 | 0.01851595463746262 |
| asd | 16: Default C | ADOS_SOCIAL | 0.14833273013258194 | 0.02265191987474737 | -0.07913674481927263 | 0.2258313290022622 |
| asd | 8: Salience/ Ventral Attention B | ADOS_SOCIAL | 0.110035740897415 | 0.09168973476842239 | -0.12457563226597086 | 0.055997274743319614 |
| asd | 11: Control A | ADOS_SOCIAL | 0.16985036113082258 | 0.008936843545318533 | -0.04013826211985027 | 0.5394869163689382 |
| asd | 7: Salience/Ventral Attention A | ADOS_SOCIAL | 0.12534594716764783 | 0.0544841060380204 | -0.08582159466202402 | 0.18889957072679311 |
| asd | 12 Control B | ADOS_SOCIAL | 0.09643372907543059 | 0.1396698220660768 | -0.14781517449494994 | 0.023134056439705987 |
| asd | 17: Temporal Parietal | ADOS_SOCIAL | 0.06172667761570535 | 0.3451037357861474 | -0.1174908161150731 | 0.07161000786565354 |
| asd | 6: Dorsal Attention B | ADOS_SOCIAL | 0.09944041436852548 | 0.1276837746301653 | -0.15321027481214902 | 0.018518458277949165 |
| asd | 9: Limbic B | ADOS_SOCIAL | 0.10801190824867353 | 0.09785423020916573 | -0.03683969493417247 | 0.5733474335206318 |
| asd | 3: Somatomotor A | ADOS_SOCIAL | 0.12004958739037179 | 0.06560564642575407 | -0.024783785614145962 | 0.7048568916840413 |
| asd | 15: Default B | ADOS_SOCIAL | 0.155305611726408 | 0.016954743393062084 | -0.038653874748288866 | 0.5546007670427495 |
| asd | 4: Somatomotor B | ADOS_SOCIAL | 0.1120089455998839 | 0.08598192913581833 | -0.10378955681040239 | 0.1117722915639283 |
| asd | 14: Default A | ADOS_SOCIAL | 0.1330230117260903 | 0.04117429037207131 | -0.1044920832443352 | 0.10935461209547061 |
| asd | 5: Dorsal Attention A | ADOS_SOCIAL | 0.14454629009188538 | 0.026388511146936543 | -0.145648519798874 | 0.02524976135203736 |
| asd | 2: Visual B | ADOS_SOCIAL | 0.06570800653002533 | 0.3148233575762252 | -0.10494278218546237 | 0.10782533287619252 |
| asd | 10: Limbic A | ADOS_SOCIAL | 0.07464162378634161 | 0.25338248948093484 | 0.003738541545326384 | 0.9544433735797472 |
| asd | 1: Visual A | ADOS_SOCIAL | 0.06030984027399827 | 0.3563105813037385 | -0.07526924234017365 | 0.24940195208813065 |
| asd | 13: Control C | ADOS_STEREO_BEHAV | 0.10980828936239653 | 0.09236668099656464 | 0.18060092286161933 | 0.005392011920004123 |
| asd | 16: Default C | ADOS_STEREO_BEHAV | 0.12721008065916123 | 0.05096209728655514 | 0.11724583784386002 | 0.07220761039241591 |
| asd | 8: Salience/ Ventral Attention B | ADOS_STEREO_BEHAV | 0.07172336486023843 | 0.2724670481209962 | 0.13220552504417715 | 0.04244695112067783 |
| asd | 11: Control A | ADOS_STEREO_BEHAV | 0.07235749991719233 | 0.26823908179492845 | 0.04188284167150374 | 0.5219898062138368 |
| asd | 7: Salience/Ventral Attention A | ADOS_STEREO_BEHAV | 0.14915278355594702 | 0.021905751883363682 | 0.10481216792520756 | 0.10826678231339991 |
| asd | 12 Control B | ADOS_STEREO_BEHAV | 0.19323251724366125 | 0.0028742355477280866 | 0.19779095605026673 | 0.0022686567469772045 |
| asd | 17: Temporal Parietal | ADOS_STEREO_BEHAV | 0.1247279063142011 | 0.055695428393024646 | 0.16145180823544744 | 0.013012947006524386 |
| asd | 6: Dorsal Attention B | ADOS_STEREO_BEHAV | 0.2000163008337471 | 0.00201742572110186 | 0.14518967962104443 | 0.025718564969065043 |
| asd | 9: Limbic B | ADOS_STEREO_BEHAV | 0.20245169776482844 | 0.00177175503839758 | 0.10958965964304003 | 0.09302112475405087 |
| asd | 3: Somatomotor A | ADOS_STEREO_BEHAV | 0.16562934927581385 | 0.010816762761058378 | 0.09659240610553774 | 0.13901648408272146 |
| asd | 15: Default B | ADOS_STEREO_BEHAV | 0.10235392901407134 | 0.11684300137368786 | -0.02261083996614585 | 0.7296777706248105 |
| asd | 4: Somatomotor B | ADOS_STEREO_BEHAV | 0.12905989998732362 | 0.04765648674763101 | 0.04184051088093337 | 0.5224109013726583 |
| asd | 14: Default A | ADOS_STEREO_BEHAV | 0.08231279036457544 | 0.2076916580514447 | 0.08094979460647017 | 0.2153433229027423 |
| asd | 5: Dorsal Attention A | ADOS_STEREO_BEHAV | 0.09416695159201488 | 0.14926127409503545 | 0.08286382076907824 | 0.20465453569619294 |
| asd | 2: Visual B | ADOS_STEREO_BEHAV | 0.07629770171288798 | 0.2429734463862597 | 0.044715479737341315 | 0.49421075396559333 |
| asd | 10: Limbic A | ADOS_STEREO_BEHAV | 0.24254451168853455 | 0.000168160906365476 | 0.028558524471160134 | 0.6624856126917755 |
| asd | 1: Visual A | ADOS_STEREO_BEHAV | 0.23524456400186794 | 0.00026634637491696936 | 0.11504964607791031 | 0.07774820197554103 |

**Supplementary Table 13**. Association of extreme deviations and IQ in SZ using anatomical atlas

| Diagnosis | ROI | Test | Correlation of % positive deviation | P value of % positive deviation | Correlation of % negative deviation | P value of % negative deviation |
|---|---|---|---|---|---|---|
| scz | Left. Crus.I | IQ | 0.13032056857038055 | 0.032631961247396514 | -0.1648012390048176 | 0.006750649691131236 |
| scz | Left.Crus.II | IQ | 0.20123853844219436 | 0.0009027309677741256 | -0.20451915176318544 | 0.0007395490107213605 |
| scz | Left.VIIB | IQ | 0.21982064976684715 | 0.0002800636386395373 | -0.2576397138137502 | 1.8826386038125192e-05 |
| scz | Left.VI | IQ | 0.20029563291206492 | 0.0009554308752920696 | -0.21395318117348502 | 0.0004096951238498148 |
| scz | Left.VIIIA | IQ | 0.13958230286980963 | 0.02202826421242531 | -0.2650197853942772 | 1.0557309896141252e-05 |
| scz | Left..VI | IQ | 0.20012399660139077 | 0.0009653233408955842 | -0.18519817290919646 | 0.002290272725104287 |
| scz | Left.X | IQ | 0.0890322487207192 | 0.14530114107873002 | -0.18694314410793 | 0.0020769226903939913 |
| scz | Left.IV | IQ | 0.19037168647181144 | 0.0017096518553360721 | -0.18864732655364302 | 0.0018862134296452654 |

| | | | | | | |
|---|---|---|---|---|---|---|
| scz | Left.VIIIA | IQ | 0.2010760431773143 | 0.0009116175440411488 | -0.23396562789809605 | 0.00010735061020158035 |
| scz | Left.IX | IQ | 0.1355756835380044 | 0.026180238222910747 | -0.14376182868087564 | 0.018316517971243777 |
| scz | Left.I.III | IQ | 0.16515173347401538 | 0.006632732762329408 | -0.17754253406010476 | 0.0034823891139989043 |
| scz | Vermis.VIII | IQ | 0.17206083881893516 | 0.004654772628948412 | -0.17301133737701532 | 0.00442896078802772 |
| scz | Vermis.IX | IQ | 0.13594235788839798 | 0.02577415850330444 | -0.1699003700989384 | 0.005207045413527454 |
| scz | Vermis.VII | IQ | 0.10962245784914541 | 0.07265646554025769 | -0.16249232756965357 | 0.0075753668522895575 |
| scz | Vermis.VI | IQ | 0.08846403747813622 | 0.14789697064720989 | -0.24595608544772332 | 4.542222832923222e-05 |
| scz | Vermis.X | IQ | 0.08961127644307122 | 0.14269173869574397 | -0.1701486613732318 | 0.005140711969527629 |
| scz | Right.V | IQ | 0.19919434359167754 | 0.001020563579859658 | -0.17676747393049305 | 0.003630043089523112 |
| scz | Right.IV | IQ | 0.19506831187205984 | 0.001302612977991931 | -0.1681535061700016 | 0.005695887463452922 |
| scz | Right.IX | IQ | 0.16618189530095934 | 0.006296742599323807 | -0.11339770755642661 | 0.0632834837834972 |
| scz | Right.I.III | IQ | 0.16661695285262715 | 0.00615948246923299 | -0.1571081823394693 | 0.009856795521379795 |
| scz | Right.Crus.II | IQ | 0.19824694926556377 | 0.0010798384473714668 | -0.21633413692872863 | 0.0003515204936565396 |
| scz | Right.VIIB | IQ | 0.23044223475950154 | 0.0001370777189962888 | -0.23354472237025417 | 0.00011055339842084345 |
| scz | Right.VI | IQ | 0.16283827086331396 | 0.007446341178363202 | -0.20720524869294016 | 0.000626714908461577 |
| scz | Right.Crus.I | IQ | 0.14825121440049774 | 0.014947832706845838 | -0.18715294951608677 | 0.002052530348765224 |
| scz | Right.Crus.II | IQ | 0.21747836659206066 | 0.0003263858526637051 | -0.26692476981403646 | 9.067455615610008e-06 |
| scz | Right.VIIIB | IQ | 0.23549231967373055 | 9.644937856350603e-05 | -0.2788537650244462 | 3.4056675401948217e-06 |
| scz | Right.X | IQ | 0.09563020833245367 | 0.11764477874264888 | -0.22288003458443884 | 0.00022875655681068633 |

**Supplementary Table 14**. Association of extreme deviations and IQ in SZ using task-based atlas

| Diagnosis | ROI | Test | Correlation of % positive deviation | P value of % positive deviation | Correlation of % negative deviation | P value of % negative deviation |
|---|---|---|---|---|---|---|
| scz | 10: Autobiographical recall/visual letter recognition/interference resolution | IQ | 0.17026353564363927 | 0.005110279919824275 | -0.12861813511274664 | 0.03499331941304757 |
| scz | 3: Saccades/visual working memory/visual letter recognition | IQ | 0.16875847476939074 | 0.005522121072975453 | -0.2406071489351213 | 6.702816585976154e-05 |
| scz | 4: Action Observation/divided attention/motor planning | IQ | 0.2361898246854599 | 9.18228095604874e-05 | -0.27764061566113885 | 3.7702583486457227e-06 |
| scz | 5: Divided attention/active maintenance/mental arithmetic | IQ | 0.16189887566644381 | 0.00780135313846835 | -0.24470391696655225 | 4.979324129909756e-05 |
| scz | 6: Divided attention/verbal fluency/active maintenance | IQ | 0.2108585104778549 | 0.0004986961097378476 | -0.235023365087175 | 9.968205539670315e-05 |
| scz | 9: Verbal Fluency/word comprehension/mental arithmetic | IQ | 0.15547787816604489 | 0.010658511094188975 | -0.23346572214140368 | 0.00011116443666094431 |
| scz | 1: Left-hand presses/ motor planning/ interference resolution | IQ | 0.23249776779387646 | 0.00011891301188986983 | -0.22836105350764077 | 0.00015809346087094426 |
| scz | 2: Right-hand presses/ motor planning/ divided attention | IQ | 0.23633133204492515 | 9.090998489423844e-05 | -0.2642849758808288 | 1.1191876469958106e-05 |
| scz | 7: Narrative/ emotion processing/ language processing | IQ | 0.18030944132301463 | 0.002998553454517152 | -0.24984046907373228 | 3.405250351376514e-05 |
| scz | 8: Word comprehension/ language processing/ narrative | IQ | 0.1478659728655771 | 0.015213936829290411 | -0.20255383821441822 | 0.0008336843799427591 |

**Supplementary Table 15**. Association of extreme deviations and IQ in SZ using resting-state atlas

| Diagnosis | ROI | Test | Correlation of % positive deviation | P value of % positive deviation | Correlation of % negative deviation | P value of % negative deviation |
|---|---|---|---|---|---|---|
| scz | 13: Control C | IQ | 0.17736411259588913 | 0.0035158895692436578 | -0.20957475437592366 | 0.0005406270583983106 |
| scz | 16: Default C | IQ | 0.13749925930862464 | 0.024109706998784244 | -0.12227438284261095 | 0.04510924237500713 |
| scz | 8: Salience/ Ventral Attention B | IQ | 0.22908845809520884 | 0.00015042724023073628 | -0.18723049307737683 | 0.002043581346606773 |
| scz | 11: Control A | IQ | 0.17937545665213211 | 0.0031545932601349655 | -0.09252700673793973 | 0.13009235711845188 |
| scz | 7: Salience/Ventral Attention A | IQ | 0.2105541667361251 | 0.0005083548009837796 | -0.2786686527935065 | 3.4590388913539712e-06 |
| scz | 12 Control B | IQ | 0.17865184579367754 | 0.003280506403107873 | -0.23155193540442107 | 0.00012697135303104182 |
| scz | 17: Temporal Parietal | IQ | 0.14434974368028716 | 0.01784075668665967 | -0.2562842999074682 | 2.0897348154989935e-05 |
| scz | 6: Dorsal Attention B | IQ | 0.2035968341237589 | 0.000782431900130277 | -0.27366064654221445 | 5.245763263966628e-06 |
| scz | 9: Limbic B | IQ | 0.19454202730892803 | 0.0013433291576938893 | -0.24074578751077558 | 6.636290699055261e-05 |
| scz | 3: Somatomotor A | IQ | 0.23989567990309654 | 7.054203118503705e-05 | -0.25626014030965355 | 2.093614817980989e-05 |
| scz | 15: Default B | IQ | 0.1717157446092557 | 0.00473928823833297 | -0.2381719237900243 | 7.978729976290001e-05 |
| scz | 4: Somatomotor B | IQ | 0.2383176093035337 | 7.896389174206331e-05 | -0.20459088738823372 | 0.0007363063722505872 |
| scz | 14: Default A | IQ | 0.1633698530311745 | 0.007251895917687008 | -0.25185409311931295 | 2.9274416722512903e-05 |
| scz | 5: Dorsal Attention A | IQ | 0.15253781782670572 | 0.012250992460062119 | -0.2192127792735248 | 0.0002914577052380149 |
| scz | 2: Visual B | IQ | 0.16525619446379375 | 0.0065979469246841465 | -0.18377529046190005 | 0.002478813087804469 |
| scz | 10: Limbic A | IQ | 0.1537639214395918 | 0.011563023429580084 | -0.12361431749939499 | 0.04278937732411642 |
| scz | 1: Visual A | IQ | -0.024949935202876302 | 0.6837383283112604 | -0.038523428001570534 | 0.5292688836689003 |

**Supplementary Table 16**. Association of extreme deviations and PANSS in SZ using anatomical atlas

| Diagnosis | ROI | Test | Correlation of % positive deviation | P value of % positive deviation | Correlation of % negative deviation | P value of % negative deviation |
|---|---|---|---|---|---|---|
| scz | Left. Crus.I | symptomsco_score_p | 0.03642458170445016 | 0.5519618065328951 | 0.05359642125939896 | 0.38125697052610197 |
| scz | Left.Crus.II | symptomsco_score_p | -0.034158106833656936 | 0.5769888097348552 | 0.039229950539560084 | 0.5217374790340171 |
| scz | Left.VIIB | symptomsco_score_p | -0.04147066712821361 | 0.4982197323742572 | 0.0985006025280529 | 0.10697738819769828 |
| scz | Left.VI | symptomsco_score_p | 0.0806236758584995 | 0.18739793892313025 | 0.044803036934332886 | 0.4643089288445583 |
| scz | Left.VIIIA | symptomsco_score_p | 0.019899713327710202 | 0.7452629211574264 | 0.08918226033728499 | 0.14462164799835642 |
| scz | Left..VI | symptomsco_score_p | -0.0008606759395730081 | 0.9887897704924844 | 0.006041704349482726 | 0.9214314383863949 |
| scz | Left.X | symptomsco_score_p | -0.002652293943994943 | 0.9654637017448675 | -0.0026419637815571204 | 0.9655981307459279 |
| scz | Left.IV | symptomsco_score_p | 0.02912436498347816 | 0.6343938924547845 | 0.01550369454798386 | 0.8001827628079907 |
| scz | Left.VIIIA | symptomsco_score_p | 0.0013657733299313752 | 0.9822118042641219 | 0.053586642533104696 | 0.38134397951519394 |
| scz | Left.IX | symptomsco_score_p | -0.005016034432978998 | 0.9347369815567823 | 0.09139395976367988 | 0.13488241854948632 |
| scz | Left.I.III | symptomsco_score_p | -0.011563286417510758 | 0.8502690590960192 | -0.03317588690232747 | 0.5879974409449271 |
| scz | Vermis.VIII | symptomsco_score_p | 0.0698056682721982 | 0.25388704683706437 | 0.025493967413621837 | 0.6772251286996359 |
| scz | Vermis.IX | symptomsco_score_p | -0.07702389538593334 | 0.20793120978573468 | -0.018573520260567047 | 0.7617100551682924 |
| scz | Vermis.VII | symptomsco_score_p | -0.0328420668315096 | 0.591760728428397 | 0.06812630104828658 | 0.26551727478374243 |
| scz | Vermis.VI | symptomsco_score_p | 0.040634302190157234 | 0.5069318128718476 | 0.057382235118493635 | 0.3484866855244718 |
| scz | Vermis.X | symptomsco_score_p | 0.021027775172132213 | 0.7313629445306943 | 0.017699730178524962 | 0.7726059106673466 |
| scz | Right.V | symptomsco_score_p | 0.023694727460493448 | 0.6988546649407734 | 0.15518587138759093 | 0.010808044556794797 |
| scz | Right.IV | symptomsco_score_p | 0.008194155986048009 | 0.8935840165742549 | 0.09349019165999274 | 0.12612456706425695 |
| scz | Right.IX | symptomsco_score_p | 0.008124536868389976 | 0.8944828649242194 | 0.01822243464982977 | 0.7660824605932028 |
| scz | Right.I.III | symptomsco_score_p | 0.015416722561326862 | 0.8012802834668672 | 0.1498105773560242 | 0.013911917535641584 |

| | | | | | | |
|---|---|---|---|---|---|---|
| scz | Right.Crus.II | symptomsco_score_p | 0.04951753125441288 | 0.41859440336082443 | 0.13291750121072618 | 0.029291812240346722 |
| scz | Right.VIIB | symptomsco_score_p | 0.08308655536172631 | 0.1742336465524982 | 0.06632219970501721 | 0.27841213052716757 |
| scz | Right.VI | symptomsco_score_p | 0.06939110102992847 | 0.2567247533930334 | 0.0834440401519386 | 0.17238140042987427 |
| scz | Right.Crus.I | symptomsco_score_p | 0.061993565430387323 | 0.3110554493409521 | 0.1222394869940849 | 0.04517104027183643 |
| scz | Right.Crus.II | symptomsco_score_p | 0.035900110635940695 | 0.5577057384197497 | 0.007990230548431817 | 0.8962172656770354 |
| scz | Right.VIIIB | symptomsco_score_p | 0.014983444779764137 | 0.8067536933378386 | 0.09919244219880248 | 0.10452385548592093 |
| scz | Right.X | symptomsco_score_p | -0.03690042206940543 | 0.5467755924984989 | 0.11669959089469086 | 0.05592250919860756 |
| scz | Left. Crus.I | symptomsco_score_n | 0.06801015539096547 | 0.2663349008049267 | 0.10600177975835093 | 0.08267800005979432 |
| scz | Left.Crus.II | symptomsco_score_n | 0.010110893404513547 | 0.8688944281093153 | 0.16019153328196942 | 0.008485366703407072 |
| scz | Left.VIIB | symptomsco_score_n | -0.010506759843587338 | 0.8638102428544482 | 0.1701870453767982 | 0.005130525358003929 |
| scz | Left.VI | symptomsco_score_n | 0.10173835746471663 | 0.09587565141351466 | 0.06826787078794719 | 0.2645229985669846 |
| scz | Left.VIIIA | symptomsco_score_n | 0.040812315662499044 | 0.5050708731138642 | 0.10235279044508776 | 0.09387624102227533 |
| scz | Left..VI | symptomsco_score_n | 0.06316389091805692 | 0.30199174839278137 | 0.028660706118812593 | 0.6398005670938565 |
| scz | Left.X | symptomsco_score_n | 0.014885760596266787 | 0.8079890065114494 | 0.08472121945557853 | 0.16588349993194046 |
| scz | Left.IV | symptomsco_score_n | 0.11672369701191303 | 0.055871495514517686 | 0.07339901463997739 | 0.23019900245134745 |
| scz | Left.VIIIA | symptomsco_score_n | 0.048778004040130586 | 0.4255859998653415 | 0.15642366984047332 | 0.010186663395710763 |
| scz | Left.IX | symptomsco_score_n | 0.018143523060130118 | 0.767066248375728 | 0.11030467419927314 | 0.07088369073982244 |
| scz | Left.I.III | symptomsco_score_n | 0.013215064808383935 | 0.8291878717426309 | -0.025559740898853284 | 0.6764392946977602 |
| scz | Vermis.VIII | symptomsco_score_n | 0.033228011677800554 | 0.5874108120094967 | 0.061387722692062344 | 0.31581684052101683 |
| scz | Vermis.IX | symptomsco_score_n | 0.013470939474301488 | 0.8259327275024384 | 0.08084778154837804 | 0.18617070441067668 |
| scz | Vermis.VII | symptomsco_score_n | 0.03178579482458896 | 0.6037403375566651 | 0.13503665730548292 | 0.026787169400193385 |
| scz | Vermis.VI | symptomsco_score_n | 0.02692881455365469 | 0.6601628066274501 | 0.03348700047763435 | 0.5845000708325231 |
| scz | Vermis.X | symptomsco_score_n | -0.03776937156986032 | 0.5373672082419165 | 0.06786120496313991 | 0.2673859805340957 |
| scz | Right.V | symptomsco_score_n | 0.0032930091491897664 | 0.9571279721507029 | 0.05089600873316269 | 0.40574301077276353 |
| scz | Right.IV | symptomsco_score_n | 0.06393152046593244 | 0.29614263819083597 | 0.00922568598653628 | 0.8802825263072729 |
| scz | Right.IX | symptomsco_score_n | 0.01595366433398108 | 0.7945107938993217 | 0.09658598079451904 | 0.11400429271911232 |
| scz | Right.I.III | symptomsco_score_n | 0.04930444301672263 | 0.4206020444520945 | 0.05976061793574285 | 0.3288387179350838 |
| scz | Right.Crus.II | symptomsco_score_n | 0.06050627235939775 | 0.32282880147577864 | 0.16939953370439526 | 0.005343193665644524 |
| scz | Right.VIIB | symptomsco_score_n | 0.06548508826666907 | 0.28453701791801345 | 0.1990986923300708 | 0.0010264092587659705 |
| scz | Right.VI | symptomsco_score_n | 0.08582699524547248 | 0.16040714624507796 | 0.08869395544313596 | 0.14684239556451517 |
| scz | Right.Crus.I | symptomsco_score_n | 0.08906799758122146 | 0.14513899269407357 | 0.08921919390081298 | 0.14445472565163714 |
| scz | Right.Crus.II | symptomsco_score_n | 0.06737971083331981 | 0.2708030620028051 | 0.17188222022116978 | 0.004698346456588632 |
| scz | Right.VIIIB | symptomsco_score_n | 0.0561628483904107 | 0.3588418866329731 | 0.14221849291094651 | 0.019618339818306636 |
| scz | Right.X | symptomsco_score_n | -0.03360472567034341 | 0.5831791821119643 | 0.19082295849661682 | 0.001666010290619648 |

**Supplementary Table 17**. Association of extreme deviations and PANSS in SZ using task-based atlas

| Diagnosis | ROI | Test | Correlation of % positive deviation | P value of % positive deviation | Correlation of % negative deviation | P value of % negative deviation |
|---|---|---|---|---|---|---|
| scz | 10: Autobiographical recall/visual letter recognition/interference resolution | symptomsco_score_p | -0.012522734440987768 | 0.8380099247032362 | 0.07248577605694087 | 0.2360656396572851 |
| scz | 3: Saccades/visual working memory/visual letter recognition | symptomsco_score_p | 0.030993116816980967 | 0.6128009524560711 | 0.06511340060862038 | 0.2872853895934117 |
| scz | 4: Action Observation/divided attention/motor planning | symptomsco_score_p | 0.029539636347475674 | 0.6295678609590343 | 0.09180887264927738 | 0.13311284589190803 |
| scz | 5: Divided attention/active maintenance/mental arithmetic | symptomsco_score_p | 0.014071276684788966 | 0.8183072193171282 | 0.07525099800967827 | 0.21862009906456592 |

| | | | | | | |
|---|---|---|---|---|---|---|
| scz | 6: Divided attention/verbal fluency/active maintenance | symptomsco_score_p | 0.04626596110824808 | 0.4498355105498282 | 0.10126406054859616 | 0.0974421110680943 |
| scz | 9: Verbal Fluency/word comprehension/mental arithmetic | symptomsco_score_p | 0.08680114239018852 | 0.15569625178673108 | 0.13764089687878744 | 0.02396299023388808 |
| scz | 1: Left-hand presses/ motor planning/ interference resolution | symptomsco_score_p | 0.05785657755880346 | 0.3445100375409129 | 0.06892622991607891 | 0.2599327439770431 |
| scz | 2: Right-hand presses/ motor planning/ divided attention | symptomsco_score_p | 0.04617153598203712 | 0.45076195879241066 | 0.09115939581152438 | 0.1358907847051739 |
| scz | 7: Narrative/ emotion processing/ language processing | symptomsco_score_p | 0.060509720626333204 | 0.3228011752133778 | 0.04065126116993617 | 0.5067543706714589 |
| scz | 8: Word comprehension/ language processing/ narrative | symptomsco_score_p | 0.050064090409900716 | 0.413470631402412 | 0.09717144038073176 | 0.11181824556068545 |
| scz | 10: Autobiographical recall/visual letter recognition/interference resolution | symptomsco_score_n | 0.06534075697944182 | 0.2856021390863768 | 0.0938949886872144 | 0.12448526689813692 |
| scz | 3: Saccades/visual working memory/visual letter recognition | symptomsco_score_n | 0.04410203060335156 | 0.47133461566251256 | 0.14659637038649473 | 0.016120491174618272 |
| scz | 4: Action Observation/divided attention/motor planning | symptomsco_score_n | 0.06114185170587324 | 0.3177626743346849 | 0.15211859170204217 | 0.01249435962088616 |
| scz | 5: Divided attention/active maintenance/mental arithmetic | symptomsco_score_n | 0.057383986784053195 | 0.3484719472916966 | 0.1325736869642038 | 0.02971649912521723 |
| scz | 6: Divided attention/verbal fluency/active maintenance | symptomsco_score_n | 0.04998478234187059 | 0.4142118140348541 | 0.0922210025384777 | 0.13137288625042698 |
| scz | 9: Verbal Fluency/word comprehension/mental arithmetic | symptomsco_score_n | 0.1199286338297208 | 0.04942426475860095 | 0.15161514086703434 | 0.012792238318122033 |
| scz | 1: Left-hand presses/ motor planning/ interference resolution | symptomsco_score_n | 0.07075544526588796 | 0.24746789995399762 | 0.08166321462540743 | 0.18175512057710888 |
| scz | 2: Right-hand presses/ motor planning/ divided attention | symptomsco_score_n | 0.03848013471505654 | 0.5297321639304051 | 0.08495163810347767 | 0.1647309695454533 |
| scz | 7: Narrative/ emotion processing/ language processing | symptomsco_score_n | 0.05113602151824319 | 0.4035295995906839 | 0.13858146672426921 | 0.023008086704128485 |
| scz | 8: Word comprehension/ language processing/ narrative | symptomsco_score_n | 0.07426861293875889 | 0.2247092360785048 | 0.05862012948344983 | 0.33816954181001047 |

**Supplementary Table 18.** Association of extreme deviations and PANSS in SZ using resting-state atlas

| Diagnosis | ROI | Test | Correlation of % positive deviation | P value of % positive deviation | Correlation of % negative deviation | P value of % negative deviation |
|---|---|---|---|---|---|---|
| scz | 13: Control C | symptomsco_score_p | 0.0320078077796168 | 0.6012134159683146 | 0.10738133391581235 | 0.0787355952171325 |
| scz | 16: Default C | symptomsco_score_p | 0.03452134508599898 | 0.5729422749499853 | 0.10405540960722535 | 0.08850915486791715 |
| scz | 8: Salience/ Ventral Attention B | symptomsco_score_p | 0.00786539988459659 | 0.8978297425224212 | 0.10676345478174167 | 0.08048211019319737 |
| scz | 11: Control A | symptomsco_score_p | -0.03871786685105702 | 0.527190728967736 | 0.09315911816335946 | 0.1274776978376273 |
| scz | 7: Salience/Ventral Attention A | symptomsco_score_p | 0.06424522928889521 | 0.2937741162243715 | 0.0920105142142452 | 0.13225933764398887 |
| scz | 12 Control B | symptomsco_score_p | 0.08770190267404707 | 0.15143385827052167 | 0.04378710323638863 | 0.4745097864559378 |
| scz | 17: Temporal Parietal | symptomsco_score_p | 0.04028234649327761 | 0.5106216492849697 | 0.05966257101785927 | 0.32963429933662625 |
| scz | 6: Dorsal Attention B | symptomsco_score_p | 0.03928682847661 | 0.5211335632168441 | 0.0319860819909306 | 0.6014604862990874 |
| scz | 9: Limbic B | symptomsco_score_p | 0.02226137170622442 | 0.716262462311938 | 0.08466284661860295 | 0.16617643014596872 |
| scz | 3: Somatomotor A | symptomsco_score_p | 0.02721256284104749 | 0.6568090079062445 | 0.06882948667642738 | 0.2606038050764983 |
| scz | 15: Default B | symptomsco_score_p | -0.021318466796504138 | 0.7277949976242428 | 0.028897309253193518 | 0.6370391638561681 |

| | | | | | | |
|---|---|---|---|---|---|---|
| scz | 4: Somatomotor B | symptomsco_score_p | 0.035192470851857324 | 0.5655012498651761 | 0.0455107176551537 | 0.4572755342123901 |
| scz | 14: Default A | symptomsco_score_p | 0.014411550312406181 | 0.8139925782050369 | 0.05928229457007871 | 0.33273169121793966 |
| scz | 5: Dorsal Attention A | symptomsco_score_p | -0.0015239458047388713 | 0.9801521244625144 | 0.06651980833085479 | 0.276979403314549 |
| scz | 2: Visual B | symptomsco_score_p | 0.0007972052222990182 | 0.9896164228265518 | 0.0356605628378267 | 0.5603388188817748 |
| scz | 10: Limbic A | symptomsco_score_p | 0.004590005471571818 | 0.9402692062438238 | 0.037842608784149585 | 0.5365779579338648 |
| scz | 1: Visual A | symptomsco_score_p | 0.1047962342713585 | 0.0862520080748275 | 0.017662541455885868 | 0.7730706483995666 |
| scz | 13: Control C | symptomsco_score_n | 0.05609280870235348 | 0.35944245750969106 | 0.16578618230676218 | 0.006423960299802995 |
| scz | 16: Default C | symptomsco_score_n | 0.033999710188612746 | 0.578757558279449 | 0.11245528691524931 | 0.06552478911250538 |
| scz | 8: Salience/ Ventral Attention B | symptomsco_score_n | 0.03336731507243254 | 0.5858443702358969 | 0.10724528150544359 | 0.079117508542387 |
| scz | 11: Control A | symptomsco_score_n | 0.007346699730789314 | 0.9045343757334239 | 0.1050426543954807 | 0.0855115444524749 |
| scz | 7: Salience/Ventral Attention A | symptomsco_score_n | 0.07751847286852598 | 0.20501775466974417 | 0.13393909358705164 | 0.028060425852819704 |
| scz | 12 Control B | symptomsco_score_n | 0.09923044582991354 | 0.10439037813523133 | 0.09282443426878507 | 0.12885697118465297 |
| scz | 17: Temporal Parietal | symptomsco_score_n | 0.07632783567561517 | 0.2120819571772678 | 0.08871913440205725 | 0.14672725441095524 |
| scz | 6: Dorsal Attention B | symptomsco_score_n | 0.07325664275711916 | 0.23110675188298657 | 0.18655177350857063 | 0.00212313196969051 |
| scz | 9: Limbic B | symptomsco_score_n | 0.03592123944569783 | 0.5574737807307022 | 0.08567479599068899 | 0.16115274420600226 |
| scz | 3: Somatomotor A | symptomsco_score_n | 0.06853996788413817 | 0.2626191730724171 | 0.08317827835492111 | 0.17375699601075312 |
| scz | 15: Default B | symptomsco_score_n | 0.05004430573999243 | 0.4136554581650329 | 0.04761086780569746 | 0.4367570730446402 |
| scz | 4: Somatomotor B | symptomsco_score_n | 0.06280809427766748 | 0.3047285682914283 | 0.036655570811744274 | 0.5494412433593459 |
| scz | 14: Default A | symptomsco_score_n | 0.005906432590664573 | 0.9231850860730173 | 0.1344214885007757 | 0.027494560230466626 |
| scz | 5: Dorsal Attention A | symptomsco_score_n | 0.015670942335996103 | 0.79807332631468 | 0.140419709880286 | 0.021236380702205522 |
| scz | 2: Visual B | symptomsco_score_n | 0.04881257051678295 | 0.4252577008113826 | 0.04133666113988212 | 0.4996102706718907 |
| scz | 10: Limbic A | symptomsco_score_n | 0.035198051219948276 | 0.5654395722506353 | 0.11171120469360671 | 0.06734001840018941 |
| scz | 1: Visual A | symptomsco_score_n | 0.11194458525414268 | 0.06676629648143365 | 0.08473102858110794 | 0.1658343130955077 |
